# Image-based, pooled phenotyping reveals multidimensional, disease-specific variant effects

**DOI:** 10.1101/2025.07.03.663081

**Authors:** Sriram Pendyala, Katie Partington, Nicholas Bradley, Abbye E. McEwen, Gwenneth Straub, Hyeon-Jin Kim, Shawn Fayer, Daniel Lee Holmes, Katherine A. Sitko, Allyssa J. Vandi, Rachel L. Powell, Clayton E. Friedman, Evan McDermot, Nishka Kishore, Frederick P. Roth, Alan F. Rubin, Kai-Chun Yang, Lea M. Starita, William S. Noble, Douglas M. Fowler

## Abstract

Genetic variants often produce complex phenotypic effects that confound current assays and predictive models. We developed Variant in situ sequencing (VIS-seq), a pooled, image-based method that measures variant effects on molecular and cellular phenotypes in diverse cell types. Applying VIS-seq to ∼3,000 *LMNA* and *PTEN* variants yielded high-dimensional morphological profiles that captured variant-driven changes in protein abundance, localization, activity and cell architecture. We identified gain-of-function *LMNA* variants that reshape the nucleus and autism-associated *PTEN* variants that mislocalize. Morphological profiles predicted variant pathogenicity with near-perfect accuracy and distinguished autism-linked from tumor syndrome-linked *PTEN* variants. Most variants impacted a multidimensional continuum of phenotypes not recapitulated by any single functional readout. By linking protein variation to cell images at scale, we illuminate how variant effects cascade from molecular to subcellular to cell morphological phenotypes, providing a framework for resolving the complexity of variant function.

## INTRODUCTION

Genetic variants change the sequence of transcripts and peptides deterministically, but their effect on molecular or cellular phenotypes is difficult to predict. For example, protein-coding variants can alter protein structure, function or localization; disrupt macromolecular complexes and subcellular processes; or alter cell internal structure, morphology and behavior. Deep mutational scanning enabled the systematic, pooled evaluation of thousands of protein-coding variants in a single experiment for some of these molecular or cellular phenotypes^1^. However, the vast majority of deep mutational scans and other multiplexed experiments have focused on simple, one-dimensional phenotypes^2^ like cell growth^3,4^ or protein abundance^5,6^ in cultured human cell lines or non-human models. These experiments generally provide limited insight into variant pathomechanism, largely fail to parse complex gene-disease relationships and cannot inform on cell-type specific effects. Consequently, genetic variants are often viewed as affecting one phenotype and classified as either benign or pathogenic. In reality, variants have complex, multidimensional phenotypic consequences. Thus, scalable methods are needed that can enable variant effect measurements on generalizable, interpretable, and information-dense molecular and cellular phenotypes in specialized cell types.

Building on *in situ* sequencing methods for image-based, pooled CRISPR screens^7–11^, we massively expanded the scope of deep mutational scanning by enabling pooled measurement of variant effects using cell imaging. Because imaging can spatially resolve biomolecules, subcellular structures and cell morphology using fluorescent proteins, *in situ* hybridization, antibodies, and cell-staining dyes, it yields insights into the effects of diverse perturbations. For example, imaging of cells with dyes that stain subcellular compartments^12^ revealed the effects of chemical^13^, CRISPR^14^, and ORF overexpression^15^ perturbations and illustrated that image-based profiling is at least as powerful as RNA-based profiling for clustering these perturbations^16^. Arrayed, image-based measurement of the localization of ∼3,000 human pathogenic variant proteins expressed in a cell line also demonstrated that mislocalization is a common defect^17^. We overcame longstanding technical challenges to develop Variant *in situ* sequencing (VIS-seq), a method based on a circular RNA barcode that yields a thirteen-fold increase in reads per cell compared to a linear construct. We also created a transgene expression cassette that prevents silencing, enabling a ten-fold increase in scale compared to previous multiplexed experiments in human induced pluripotent stem (iPS) cells^18–21^ and their derivatives. Thus, VIS-seq is a scalable platform that uses imaging of cells to quantify the effect of variants on diverse phenotypes in disease-relevant cell types.

We used VIS-seq to derive morphological profiles for ∼3,000 synonymous, missense, nonsense and frameshift variants of *LMNA* and *PTEN* from ∼11.4 million cell images. We chose these genes because variants in each are associated with multiple diseases, and the underlying molecular and cellular mechanisms linking variants to each disease remain poorly understood. Morphological profiles of *LMNA* variants expressed in U2OS^22^ cells revealed extensive effects on the lamin A protein’s abundance, localization, aggregation, and ability to maintain nuclear shape. *LMNA* profiles enabled us to discover gain-of-function variants that increase nuclear circularity and revealed a direct connection between lamin A’s three-dimensional protein structure, localization and effect on cells. Morphological profiles of *PTEN* variants expressed in both iPS cells and derived neurons comprehensively mapped the interdependence of the PTEN protein’s abundance, activity, and localization, and revealed a nuclear localization defect enriched among autism spectrum disorder-associated variants. *LMNA* and *PTEN* morphological profiles identified known pathogenic variants with near-perfect accuracy and, in the case of *PTEN*, enabled us to discriminate between autism and tumor syndrome-associated variants. Thus, VIS-seq expands our ability to deeply phenotype genetic variants in different cell contexts, illuminating how variant effects propagate from molecules to subcellular structures to cells. More fundamentally, we show that variants perturb a phenotype space larger and more complex than previously explored, spanning a multidimensional continuum rather than a pathogenic-benign binary and highlighting the limitations of current one-dimensional experimental or predictive methods.

## RESULTS

### Barcoded circular RNAs and a universal expression cassette enable variant in situ sequencing

Variant *in situ* Sequencing (VIS-seq) is a platform for pooled, image-based profiling of thousands of transgenically expressed protein-coding variants (**Fig. 1a**). VIS-seq comprises a cassette with a promoter expressing the protein variant and a second promoter expressing an abundant circular RNA^23,24^ containing one or more barcodes that are sequenced *in situ* to reveal the identity of the variant expressed in each cell (**Fig. 1b**). The VIS-seq expression cassette includes flanking insulators^25^, a ubiquitous chromatin opening element^26^, and an intron^27,28^, along with a viral regulatory element^29^. Together, these elements prevent transgene silencing in iPS cells before and after differentiation, eliminating a longstanding barrier to transgene-based experiments in stem cells and their derivatives^30^. The cassette sustained expression of WT mEGFP^31^-tagged histone H1.4 over 14 days of passaging in human iPS cells and after 56 days in differentiated iPS-derived cardiomyocytes, whereas expression from a lentiviral construct was completely silenced (**Fig. 1c**). We initially employed a linear RNA barcode that yielded zero or one *in situ* sequencing reads in 56% of all cells and 28% of non-silenced cells, preventing effective barcode calling (**Fig. 1d**). We therefore developed a circularizing, *in situ* sequencing-compatible RNA barcode^23,24^ based on the Twister ribozyme. 93% of cells expressing this circularizing RNA barcode had two or more reads, and reads per cell were increased ∼13-fold in all cells and ∼5-fold in non-silenced cells.

**Figure 1:**
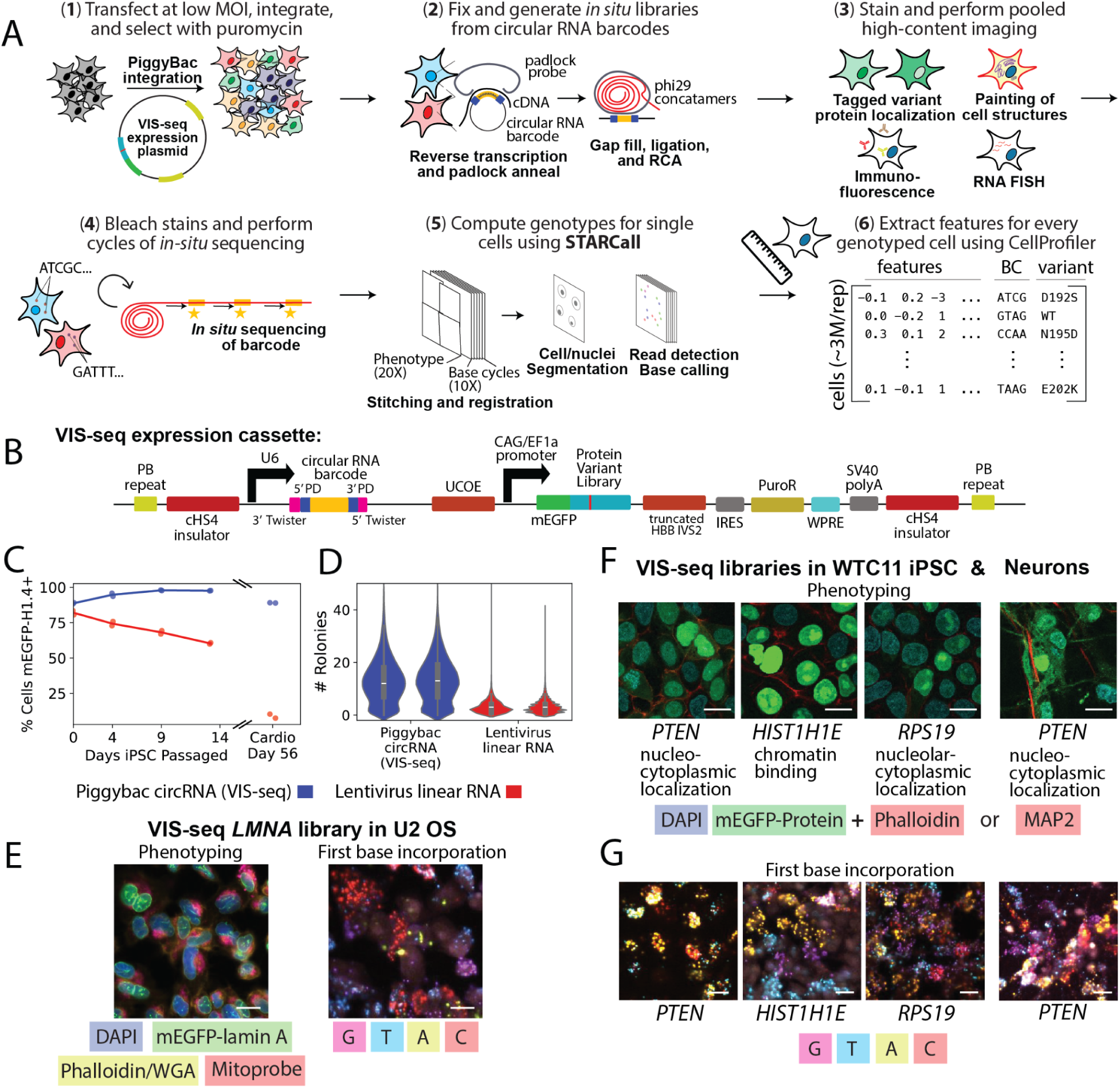
Transgenic variant expression and *in situ* sequencing with VIS-seq. (A) VIS-seq uses fluorescent *in situ* sequencing of abundant circular RNA barcodes to genotype cells expressing protein variants. (1) A variant library in the VIS-seq expression cassette is integrated into cells via *piggyBac*-ase. (2) Cells are fixed; barcodes are reverse transcribed, captured with a padlock probe and amplified; (3) cells are stained and imaged; (4) barcode is sequenced *in situ*; (5) single cell phenotype-genotype pairs are determined using STARCall; and (6) features for each cell are extracted using CellProfiler. (B) Schematic of VIS-seq expression cassette. PB = *piggyBac*^32^; PD = padlock probe^127^ binding site; UCOE = universal chromatin opening element^26^; HBB IVS2 = hemoglobin subunit beta intervening sequence 2^27,28^; IRES = internal ribosome entry site^128^; WPRE = woodchuck hepatitis virus post-transcriptional regulatory element^29^; cHS4 = chicken beta globin locus control region hypersensitive site 4^25^. (C) Time-series plot of mEGFP positivity of human WTC11 iPS cells and derived cardiomyocytes. mEGFP-tagged WT histone H1.4 is expressed via the VIS-seq cassette (blue) or lentivirus (red). Two replicates are plotted. (D) Violin plot of the number of rolling circle amplification colonies per cell after one cycle of sequencing for VIS-seq expression cassette (blue) and lentivirus (red) cells from 1F at day 14. Silenced cells were excluded. Two replicates are shown. (E) Images of VIS-seq library of mEGFP-tagged lamin A variants in U2OS cells, stained with DAPI, phalloidin, WGA, and mitoprobe (left). First base *in situ* sequencing of the same field-of-view; magenta=guanine, blue=thymidine, yellow=adenine, and red=cytosine (right). Some cells express lamin A variants that mislocalize into aggregates. Scale bar indicates 20 μm. (F) Images of libraries of mEGFP-tagged PTEN in human WTC11 *PTEN*-KO inducible-NGN2 iPS cells and mEGFP-tagged H1.4 and RPS19 in human WTC11 iPS cells, stained with DAPI and phalloidin-CF568 (left). Imaging of mEGFP-tagged PTEN library in derived neurons, seven days after induction of NGN2, stained with DAPI and anti-MAP2 AF568 (right). Some cells express variants with localization defects (PTEN=nucleocytoplasmic, HIST1H1E=chromatin binding, RPS19=nucleolar-cytoplasmic). Scale bar indicates 20 μm. (G) Imaging after first base incorporation of VIS-seq libraries from (F). *PTEN* library contains two 8bp barcodes per variant in each cell, whereas other libraries have one 12bp barcode per cell. Nucleobase coloring identical to (C). Scale bar indicates 20 μm.

VIS-seq starts with integration of a library-bearing expression cassette via PiggyBac transposition^32^ at low multiplicity of integration (MOI) (**Fig. 1a**, **Supplementary Fig. 1a**). Then, cells are fixed and barcodes are reverse transcribed, captured with a padlock probe and amplified^7,33^. Cells are stained with the desired combination of fluorescent antibodies, RNA FISH probes, and dyes for subcellular structures^12^. Images are collected and stains and probes are removed or bleached^10,34,35^. A sequencing primer is hybridized to the amplified barcodes which are sequenced *in situ*^7^. Images from each well are aligned and stitched in a single step using STitching, Alignment and Read Calling for *in situ* sequencing (STARCall), software we developed to reduce the need to maintain precise registration between successive rounds of phenotyping and *in situ* sequencing images. Barcode sequences are extracted and mapped to protein variants, then cell and nuclear borders are segmented^36,37^. Lastly, morphological features are extracted using CellProfiler^38^, generating a single cell matrix of image features (**Fig. 1a**).

To test the VIS-seq workflow, we constructed and integrated mEGFP-tagged variant libraries of four genes with variant mislocalization phenotypes,^17,39^ *LMNA, PTEN, RPS19*, and *HIST1H1E*, into U2OS or human iPS cells. These libraries elicited expected mislocalization phenotypes, including in derived neurons, and we could sequence the first base of the barcodes *in situ* (**Fig. 1e-g**, **Supplementary Fig. 1b**). Thus, VIS-seq enables protein variant expression and *in situ* sequencing in diverse cell types.

### VIS-seq reveals morphological impact of thousands of LMNA variants

We used VIS-seq to phenotype variants in the rod domain of the intermediate filament protein lamin A, encoded by *LMNA*. Lamin A assembles into a poorly understood meshwork lining the inner nuclear membrane maintaining nuclear shape and mechanical stability^40,41^ while connecting the nucleus to the cytoskeleton^42,43^ and regulating gene expression at the lamina^44,45^. *LMNA* missense variants cause a wide range of diseases, collectively called laminopathies, including Emery-Dreifuss muscular dystrophy^46^ and dilated cardiomyopathy^47^ or, less commonly, Dunnigan-type familial partial lipodystrophy^48^, Charcot-Marie-Tooth neuropathy^49^, or progeroid syndromes^50,51^. Immunostaining of lamin A in patient fibroblasts^52^ and transient transfection of lamin A variants into cell lines^39,53^ revealed altered localization and morphology, with some pathogenic rod domain variants leading to lamin A aggregates, changes in nuclear shape, and other nuclear defects. However, not all pathogenic *LMNA* variants form aggregates, and the relationship between these molecular and cellular phenotypes and the diverse clinical phenotypes remains poorly understood^39^.

We created a library of >1,700 mEGFP-tagged *LMNA* synonymous, missense and frameshift variants of the central rod domain between amino acid positions 178 and 273, a hotspot for pathogenic variants. We integrated this lamin A library into U2OS *LMNA*-KO cells using the VIS-seq expression cassette and imaged ∼10 million fixed, library-expressing cells after staining chromatin (DAPI), microfilaments (phalloidin), plasma membrane (wheat germ agglutinin, WGA)^12^, and mitochondria (anti-12S and 16S mitochondrial rRNA FISH probes; **Fig. 2a**, **Supplementary Fig. 1c**)^10^. *In situ* sequencing of the 12-base variant barcode followed by filtering to remove cells with two or more barcodes yielded ∼3,000 cells for each synonymous or missense variant as well as ∼900,000 cells expressing wild type (WT) lamin A (**Fig. 2b, Supplementary Fig. 2a-c**). We used CellProfiler^38^ to extract 1,077 morphological features for each cell that collectively captured the intensity, distribution, texture, and granularity of the mEGFP-lamin A and stain channels, as well as correlations between channels (**Fig. 2c**). Thus, we obtained image and feature information for 1,767 variants expressed in ∼6.6 million cells over two replicate experiments. The number of genotyped cells (Pearson’s r=0.98) and median nuclear lamin A intensity (r=0.94) for each variant were highly concordant across replicates (**Supplementary Fig. 2d,e**).

**Figure 2:**
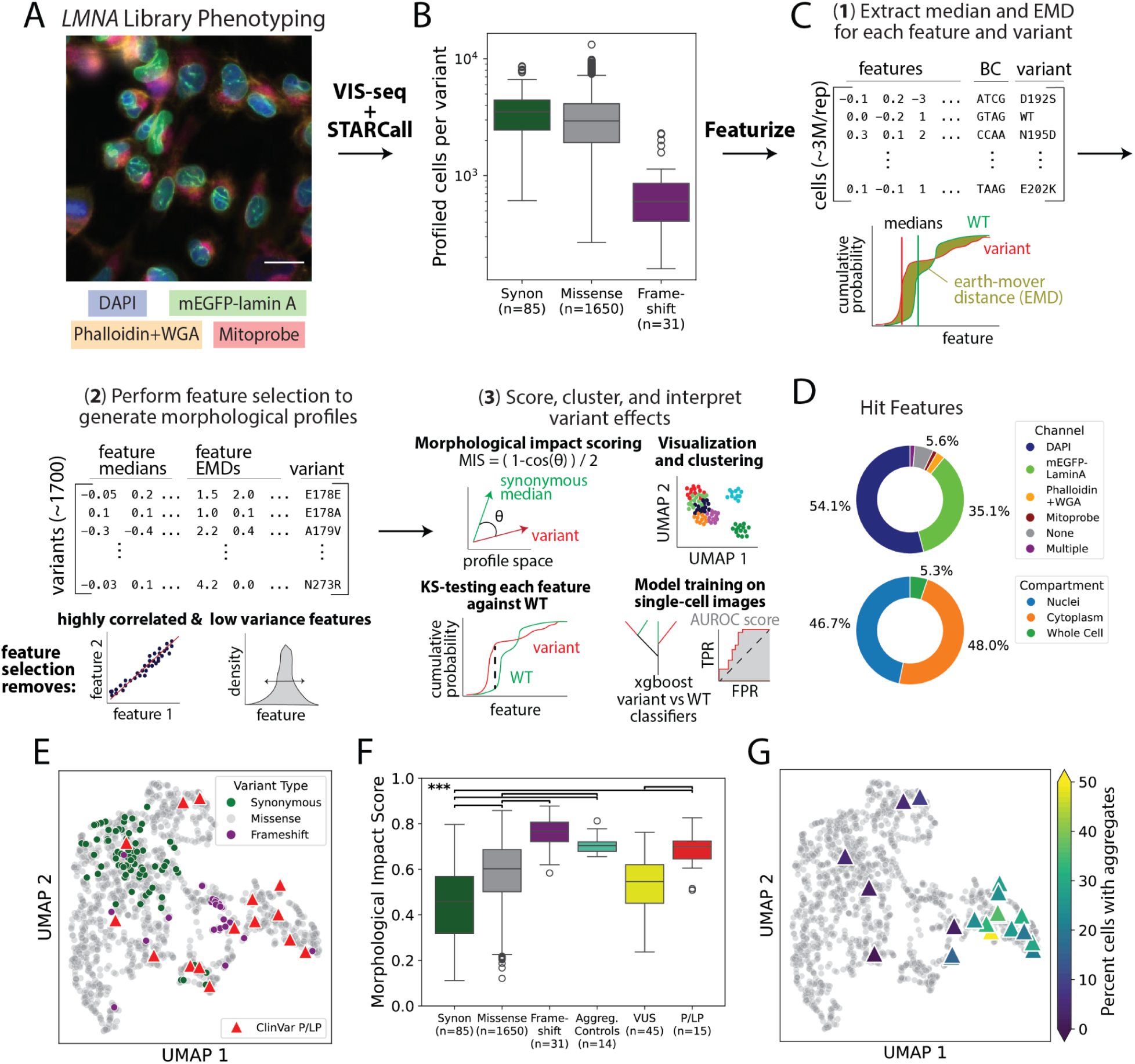
Morphological profiles of >1,700 *LMNA* variants identify variants that alter function, including aggregating and pathogenic variants. (A) Representative image of a library of mEGFP-tagged lamin A variants in U2OS cells, stained with DAPI, phalloidin and WGA (wheat germ agglutinin), and mitoprobe^10^. Scale bar indicates 20 μm. (B) Number of profiled single cells per variant. (C) Morphological profile analysis workflow. (1) A matrix of ∼6.6 million single cells each with 1,077 CellProfiler-derived features, a sequenced barcode and associated lamin A variant was generated. (2) Feature medians and earth-mover distances were computed among all cells expressing each variant; feature selection removed features that were highly correlated, with low variance, or that are biologically irrelevant. (3) Variant embeddings were visualized using UMAP following dimensionality reduction with PCA. Morphological impact score for each variant was computed using cosine distance. Variant single-cell feature distributions were KS-tested against WT. AUROC scores for each variant reflect the ability of a model trained to distinguish variant from WT using single cell feature profiles. (D) Hit features (Bonferroni-corrected KS-test p<0.001 for >=25 *LMNA* variants), shown by imaging channel (top) or compartment (bottom). (E) UMAP visualization of PCA-transformed morphological profiles for each variant. Circles indicate synonymous (green), missense (grey) and frameshift (purple) variants, triangles indicate ClinVar likely pathogenic/pathogenic (LP/P, red). (F) Boxplots of morphological impact scores for variants that aggregated in >15% of HEK 293T cells^39^ (ClinVar variants of uncertain significance=VUS and likely pathogenic/pathogenic variants=LP/P). *** indicates Mann-Whitney p-value < 0.001. (G) UMAP visualization from (E). Colored triangles indicate previously measured fraction of cells with lamin A aggregates in HEK 293T cells^39^.

To summarize phenotype heterogeneity across single cells, we computed the median and earth-mover distance (EMD)^54,55^ from WT for each feature for every variant (**Fig. 2c**). Then, we removed features that had low variance, high correlation to other features, reflected biologically irrelevant information (e.g., pixel coordinates, cell orientation)^56^, or had EMDs with low reproducibility^55^ (**Supplementary Fig. 2f**). We retained 333 z-score scaled feature medians and EMDs for each profiled variant, which we refer to as “variant morphological profiles”. Synonymous variants, which do not change the protein’s sequence and are therefore expected to negligibly impact morphology, had fewer altered features than missense or frameshift variants (**Supplementary Fig. 2g**). Features from both nuclear and cytoplasmic compartments derived from the DAPI and mEGFP-lamin A channels were most likely to be altered (**Fig. 2d**). *LMNA* variant features and morphological profiles can be explored at https://visseq.gs.washington.edu/.

### Morphological profiles detect variant aggregation and summarize impact and pathogenicity

Dimensionality reduction and visualization^57^ of variant morphological profiles revealed that synonymous variants formed a distinct group (**Fig. 2e**). To summarize the dissimilarity between each variant’s profile and WT, we developed two metrics. The first is a morphological impact score describing the size of a variant’s effect on its morphological profile (**Fig. 2c, Supplementary Table 1**). The second is the area under a receiver operating characteristic curve (AUROC), describing the performance of a model trained to classify morphological profiles of single cells expressing the variant versus profiles of cells expressing WT (**Fig. 2c**, **Supplementary Fig. 3a, Supplementary Table 1**). The AUROC score is thus a single-cell measure of a variant’s distinguishability from WT rather than the magnitude of its effect on cells. Morphological impact (Pearson’s r=0.88) and AUROC scores (r=0.92) were highly concordant between replicates (**Supplementary Fig. 2h, Supplementary Fig. 3b**). Both impact and AUROC scores were significantly lower for synonymous variants than for missense and frameshift variants, with AUROC scores being more sensitive at discriminating different classes of variants (**Fig. 2f**, **Supplementary Fig. 3c**).

Next, we validated the morphological profiles, impact scores and AUROC scores using two types of control variants. 14 variants known to form aggregates in HEK293T cells^39^ segregated from synonymous variants on the UMAP and had significantly higher morphological impact and AUROC scores, illustrating the power of unbiased morphological profiling to detect this known lamin A phenotype (Mann-Whitney U p<0.001; **Fig. 2f,g**, **Supplementary Fig. 3c**). We also curated 15 pathogenic (P) or likely pathogenic (LP) variants from ClinVar^58^, removing variants predicted to affect splicing^59^. All P/LP variants except p.Leu204Arg, a single-submitter ClinVar variant in an individual with dilated cardiomyopathy, occupied regions of the UMAP distinct from nearly all synonymous variants and had markedly elevated morphological impact and AUROC scores compared to synonymous variants and variants of uncertain significance (VUS; Mann-Whitney U p<0.001; **Fig. 2e,f**, **Supplementary Fig. 3c**). Thus, unbiased summarization of morphological profiles at both variant- and single-cell resolution differentiated aggregating and pathogenic control variants from synonymous variants, with near perfect accuracy in the case of AUROC scores (**Supplementary Fig. 3f**).

### LMNA variant morphological profiles form clusters with explainable effects

Next, we organized variants by morphological profile similarity into ten clusters using the Louvain algorithm (**Fig. 3a**). We ranked features in the morphological profiles by their ability to drive cluster separation and chose four interpretable, highly-ranked “landmark features” that spanned molecular phenotypes, subcellular localization, and cell morphology: lamin A intensity in the nucleus, lamin A intensity at the nuclear boundary, lamin A nuclear granularity, and nuclear circularity (**Fig. 3b-d**, **Supplementary Table 1**). Each cluster was characterized by distinct effects on one or more of the four landmark features (**Fig. 3b**). 76 (90%) synonymous variants occupied clusters 5, 6, and 7, which, along with 519 (31%) missense variants, had little effect on landmark features and were similar to WT (**Fig. 3b,e**). The known aggregating variants were exclusively present in clusters 8, 9 and 10, along with 380 (24%) other missense variants^39^ (**Fig. 2g, 3b**). Variants in these clusters had low nuclear and boundary lamin A abundance and high granularity, presumably as a consequence of relocalization from the nuclear boundary to aggregates (**Fig. 3b,d,e**). These variants also exhibited the largest decrease in nuclear circularity, leading to both more elliptical and more concave nuclei and indicating that these variants abrogate lamin A’s ability to maintain nuclear shape^40,43,60^ (**Supplementary Fig. 4a,b**). The pathogenic variant p.Asn195Lys, along with six other pathogenic variants (47% total) were also found in these clusters (**Fig. 3c**). The 454 (28%) missense variants in clusters 2, 3 and 4 had smaller decreases in nuclear and boundary abundance and nuclear circularity, suggesting a hypomorphic phenotype (**Fig. 3b,e**). The pathogenic and known partially-aggregating^39^ variant p.Arg190Gln, along with five other pathogenic variants (40% total), were also found in these clusters (**Fig. 3c**). Surprisingly, the 292 (18%) missense variants grouped in cluster 1 drove large increases in nuclear circularity, suggesting that these variants, including p.Glu228Val,^61^ a pathogenic variant associated with multiple overlapping clinical phenotypes, may exert their effects via a gain-of-function mechanism (**Fig. 3b,c,e**).

**Figure 3:**
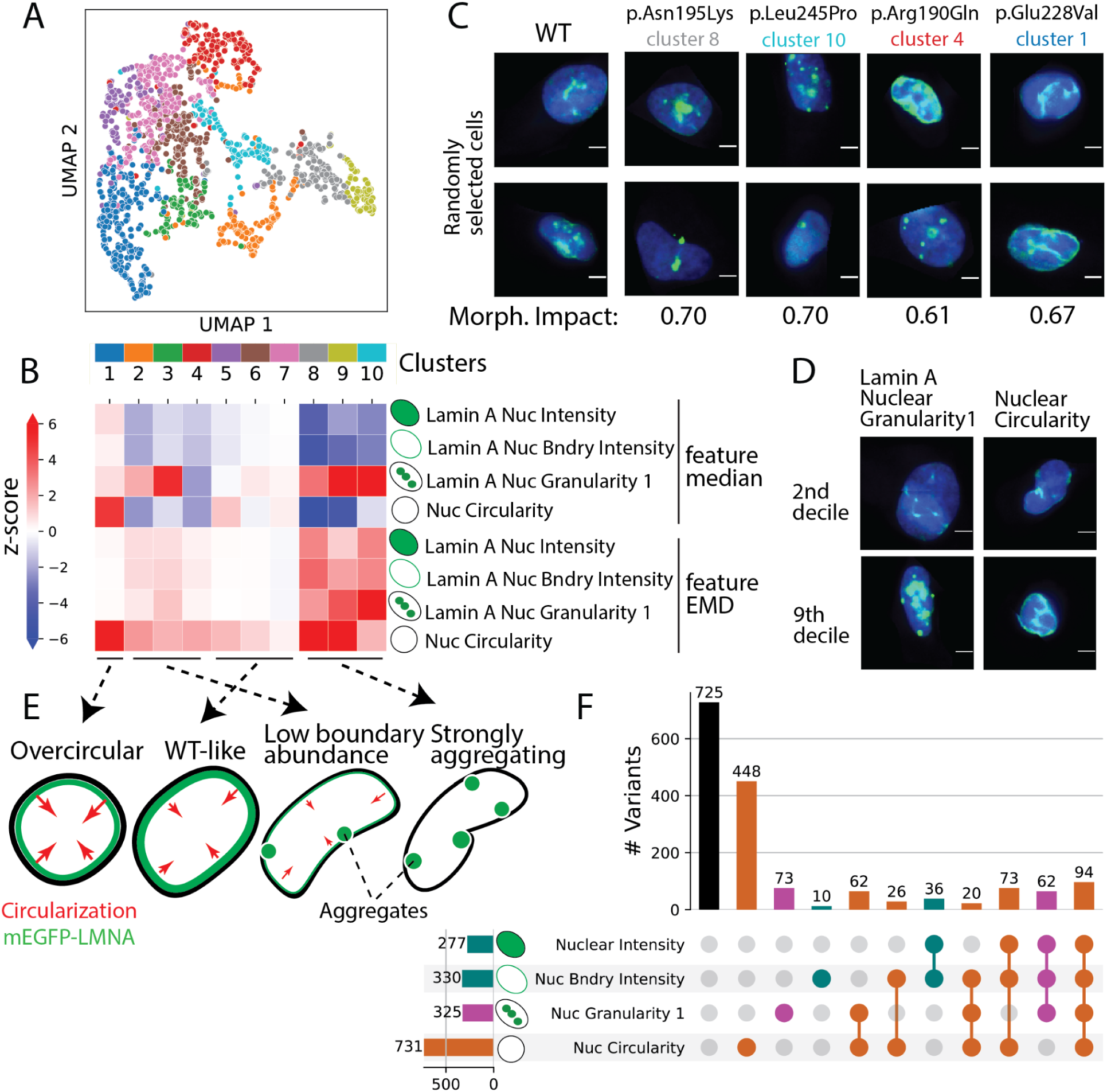
Clustering *LMNA* variant morphological profiles yields rich, interpretable phenotypes. (A) UMAP of *LMNA* variant profiles (2E) colored by Louvain^124^–derived clusters. Colors legend at bottom. (B) Landmark feature medians and EMDs for each cluster are colored by z-score versus the synonymous variant distribution. The nuclear mEGFP-lamin A intensity feature measures the average lamin A pixel intensity in the nucleus, the boundary mEGFP-lamin A intensity feature measures the average lamin A pixel intensity within two pixels of the nuclear boundary. The nuclear mEGFP-lamin A granularity 1 feature is related to the presence of lamin A aggregates, with high values indicating aggregates. The nuclear circularity feature is the normalized ratio of nuclear perimeter squared to nuclear area, with high values indicating a circular nucleus. (C) Images of two randomly-selected cells expressing the indicated variants in clusters 1, 4, 8 and 10, with morphological impact scores (mEGFP-tagged lamin A=green, DAPI=blue). Scale bar indicates 5 μm. (D) Images of randomly-selected cells from the second and ninth deciles of the distribution of lamin A nuclear granularity 1 and nuclear circularity feature values (mEGFP-tagged lamin A=green, DAPI=blue). Scale bar indicates 5 μm. (E) Graphical representation of different localization and nuclear morphologies associated with lamin A variant clusters. (F) UpSet plot showing the number of missense *LMNA* variants that impact each combination of landmark features. Features were considered impacted if EMD z-score>2.5 and KS-test Bonferroni-corrected p<0.01). Sets, represented by bars and dots in the plot, are colored by the most frequently impacted feature in the set (no impacted features=black, circularity=orange, granularity=purple, intensity/boundary intensity=teal).

In addition to defining clusters based on variant morphological profiles, we explored the relationship between landmark features (**Fig. 3e,f**). 44% of profiled missense variants perturbed nuclear circularity, making it the most frequently perturbed landmark feature. Only 20% drove changes in nuclear lamin A granularity, despite aggregation being readily appreciable in images of patient-derived cells^39,53,62^. 24% of missense variants impacted multiple landmark features, and 5.7% impacted all four features. Thus, VIS-seq segregated variants into distinct groups by their effects on lamin A intensity, localization, aggregation, and nuclear shape and revealed a new class of variants that produce a gain of nuclear circularization function.

### VIS-seq reveals the structural basis of LMNA variant effects

Next, we compared lamin A morphological profiles to prior crystal structures^63–65^ and models^66^ to understand the structural basis for the segregation of variants into clusters. Lamin A forms a soluble dimer in the cytoplasm that assembles into a multimeric filament and creates a meshwork at the lamina. The region of lamin A we mutagenized spans the rod domain comprising two ⍺-helical coiled coil subdomains, coils 1B and 2A, separated by the linker 12 subdomain (**Fig. 4a**). In other intermediate filament protein structures, linker 12 is flexible and adopts non-helical conformations, but in crystal structures of partial lamin A filaments, linker 12 is ⍺-helical^63,64,67^. An alternate model suggests that the functional state of linker 12 is flexible and non-⍺-helical^66^. To explore the relationship between these lamin A structures and function, we created variant effect maps for each landmark feature alongside UMAPs colored by feature scores (**Fig. 4a,b**, **Supplementary Fig. 4c-h**). The two most notable features of these heatmaps are the differences between variant effects at proposed coil and linker positions and the profound effects of proline substitutions, which disrupt ⍺-helices. Moreover, coil and linker substitutions had similar morphological impact scores but different morphological effects (**Supplementary Fig. 5a,b**). In coil 1B and coil 2A, 190 (25%) missense substitutions resided in clusters 8, 9 and 10 and were characterized by relocalization of lamin A to aggregates and corresponding low nuclear circularity. However, in linker 12, 370 (45%) missense substitutions resided in cluster 1, characterized by increased circularity and normal lamin A abundance. Proline substitutions had the highest morphological impact scores of all, but their morphological effects depended on their location (Mann-Whitney U p<0.001; **Fig. 4a**, **Supplementary Fig. 5c**). To further explore the effects of proline substitutions, we clustered them by their morphological profiles, yielding three distinct clusters that separated by subdomain (**Fig. 4c-e**). The clusters aligned nearly perfectly with the alternate structural model^66^, suggesting that the linker 12 subdomain is functionally disordered and providing support for flexible linker positions, linker-proximal positions, and ⍺-helical coil positions where variants have distinct effects.

**Figure 4:**
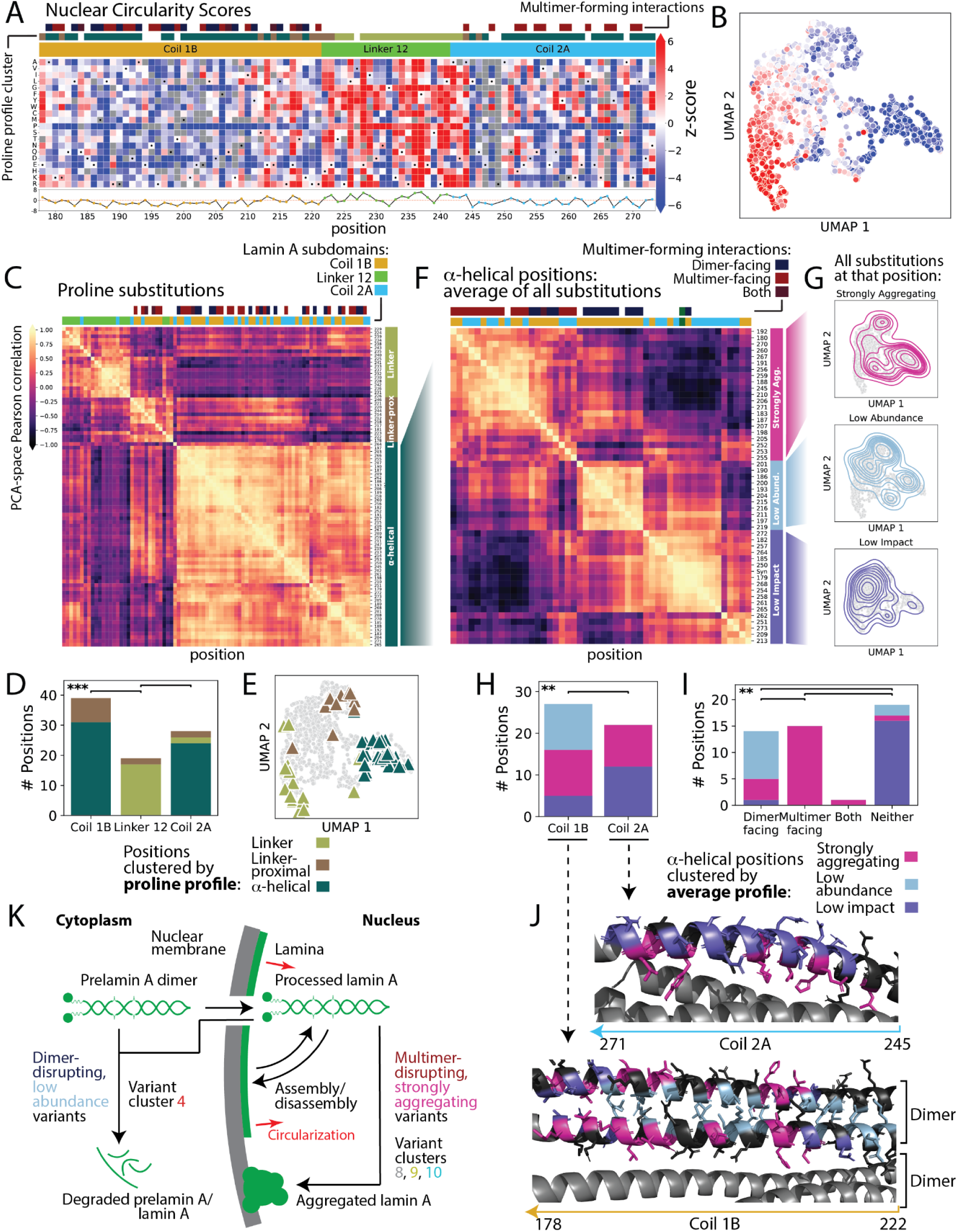
Disruption of lamin A multimerization dictates effects on aggregation and abundance. (A) Map of lamin A missense variant effects on the nuclear circularity feature (grey boxes=missing variants, black dots=synonymous substitutions). Blue to red coloring indicates the nuclear circularity feature z-score versus the synonymous variant distribution. Positions are annotated by their participation in multimer contacts (top row, defined in (F)), proline-substitution profile (middle row, defined in (C)) and coil or linker subdomain (bottom row^63^). (B) UMAP of variant profiles (2E) colored by nuclear circularity score as in (A). (C) Pearson’s correlation coefficients between *LMNA* proline substitution morphological profiles in PCA space (light color=positive correlation, dark=negative correlation). Bars on top indicate each position’s subdomain and participation in multimer contacts (see Methods, color legend shown in (F)). Bars on right indicate dendrogram-derived grouping into three clusters: linker, linker-proximal and ⍺-helical. (D) The number of proline substitutions is plotted for each subdomain^63^ colored by clusters defined in (C), with legend below. *** indicates Mann-Whitney U p-value < 0.001. (E) Proline substitutions are shown as triangles on the UMAP shown in (2E), colored by clusters defined in (C) with legend below. (F) Pearson’s correlation coefficients between the ⍺-helical (defined in (C)) PCA space, position-averaged variant morphological profiles (light color=positive correlation, dark=negative correlation). Synonymous variants were averaged into a single PCA vector and included (green annotation, top). Bars on top indicate each position’s subdomain^63^ (color legend shown in (C)) and participation in multimer contacts^63,64^ (see Methods). Bars on right indicate dendrogram-derived grouping into three clusters: strongly aggregating, low abundance, and low impact. (G) Contour plots indicate location on the UMAP shown in (2E) for missense substitutions at each group of positions defined in (F), labeled according to their effects on lamin A. (H) The number of ⍺-helical positions (defined in (C)) plotted by subdomain^63^ and colored by clusters defined in (F), with legend below. ** indicates Mann-Whitney U p-value < 0.01. (I) The number of ⍺-helical positions (defined in (C)) plotted by their participation in contacts (see Methods), colored by cluster as defined in (F). Color legend shown below. ** indicates Mann-Whitney U p-value < 0.01. (J) A11 tetramer structure 6JLB^63^ (bottom) and the A22 tetramer structure^64^ (top) with positions colored by their group as defined in (F) (position not clustered=black, magenta=strongly aggregating, light blue=weakly aggregating, dark blue=low impact). (K) Graphical representation of lamin A processing, localization, aggregation, and degradation, annotated with their relationship to dimerization, multimerization and phenotypic effects.

We further investigated the coil positions. Helices in the coiled coils can make contacts with adjacent helices across a dimer-forming interface in coil 1B and multimer-forming interfaces in both coil 1B and 2A^63,64^. To probe the nature of these contacts, we clustered ⍺-helical positions based on the average morphological profile across all substitutions at each position (**Fig. 4f**). We identified three groups of positions where substitutions were either strongly aggregating, low abundance or low impact, with corresponding significant differences in morphological impact scores and variant cluster segregation (Mann-Whitney U p<0.001; **Fig. 4f,g**, **Supplementary Fig. 5d,e**). The disparate morphologies characterizing substitutions in each of the three groups of positions were dictated by whether variants occurred at dimer or multimer interfaces (**Fig. 4h-j**, **Supplementary Fig. 5f-h**). For example, 67% of dimer–facing positions, which occur only in coil 1B, were in the low abundance group, whereas 100% of multimer-facing positions, which occur in both coil 1B and 2A, were in the strongly aggregating group. 86% of non-interacting residues, as well as the synonymous variant average profile, were in the low impact group. (**Fig. 4f**)

Thus, unsupervised clustering of *LMNA* variant morphological profiles yielded insight into lamin A structure. The effects of variants in the linker 12 subdomain on lamin A abundance and localization, as well as nuclear circularity, suggest that linker 12 is not a static coiled coil but, as postulated by the alternate structural model, is flexible^66^. The effects of variants at ⍺-helical positions were largely dictated by their location in dimer or multimer interfaces. Here, disruptive variants at dimer interfaces caused low abundance but apparently did not prevent formation of lamin A multimers needed to circularize the nucleus (**Fig. 4k**). Disruptive variants at multimer interfaces resulted in lamin A aggregation and profound loss of nuclear circularity. Rather than a simple continuum of loss of function based on protein folding or stability, disruption of different aspects of lamin A structure and assembly led to diverse effects. VIS-seq enabled us to trace these effects as they cascade from molecular phenotypes like abundance and localization to cellular phenotypes like nuclear morphology.

### Morphological profiling of PTEN variants in iPS cells and derived neurons

We next focused on Phosphatase and TENsin homolog (PTEN), a lipid and protein phosphatase that negatively regulates the PI3K-AKT-mTOR pathway and plays roles in cell growth, cell migration, and genome stability^68–73^. Pathogenic germline *PTEN* variants give rise to a spectrum of autosomal dominant tumor predisposition phenotypes^74^ and are a cause of autism spectrum disorder, often co-occurring with macrocephaly and developmental delay^75^. Somatic *PTEN* mutations occur frequently in cancers, including glioblastoma, prostate cancer, melanoma, and endometrial carcinoma^68,76^. Patients with *PTEN* variants can present with a complex mix of these phenotypes, and the relationship between changes in PTEN molecular function and disease remains unclear^77,78^. To explore this relationship, we collected morphological profiles for 1,228 PTEN catalytic domain synonymous, missense, nonsense, 3-nt deletion and frameshift variants in >2.8 million human iPS cells and >2 million iPS cell-derived neurons in two replicate *PTEN*-knockout clonal cell lines^79^ (**Fig. 5a**, **Supplementary Fig. 6a-d**, **Supplementary Fig. 1d**). iPS cells were imaged with dyes staining chromatin (DAPI), microfilaments (phalloidin), and endoplasmic reticulum (concanavalin-A)^12^ as well as mEGFP-tagged PTEN and an antibody staining phospho-Thr308-AKT (pAKT) which reports PTEN lipid phosphatase activity^80^. Neurons were imaged with the same dyes and antibodies except that a marker of neuronal differentiation^81^ (anti-MAP2 antibody) replaced phalloidin. We refined the VIS-seq protocol to include two barcodes^82^ sequenced simultaneously in each cell, reducing the number of *in situ* sequencing cycles from 12 to 8 (**Supplementary Fig. 6a-c**). The number of *in situ* genotyped cells per variant was highly concordant between iPS cells and the corresponding neuronal population after each differentiation (Pearson’s r=0.97, 0.90; **Supplementary Fig. 6e**).

**Figure 5:**
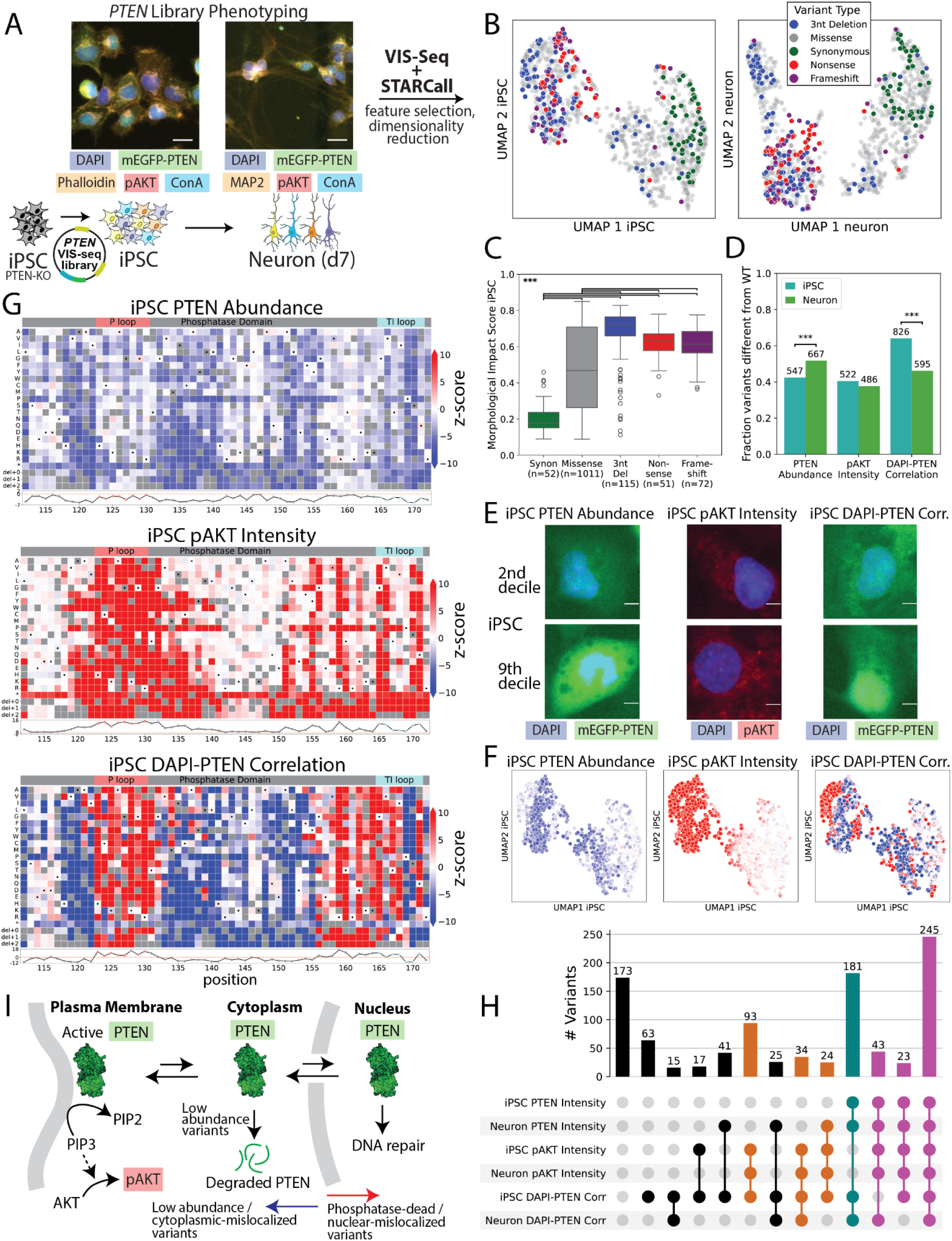
Morphological profiles of >1,200 *PTEN* variants in iPS cells and neurons. (A) Image of *PTEN* library at positions 112-172 expressed in human *PTEN*-KO iPS cells carrying dox-inducible NGN2 (left) and derived neurons 7 days after induction of NGN2 (right). MAP2=anti-MAP2 antibody, ConA=concanavalin A. Scale bar indicates 10 μm. (B) UMAP visualization of 1,228 PTEN variant morphological profiles collected in iPS cells (left) and derived neurons (right). Point color indicates variant type: synonymous variants (green), missense (grey), 3-nt deletions (blue), nonsense (red), and frameshift variants (purple). (C) *PTEN* iPS cell variant morphological impact scores plotted by variant type, colored as in (B). (D) Barplot of fraction of variants that perturb landmark features in iPS cells and neurons. For each variant, a feature was considered perturbed if |z-score|>2.5 and KS-test Bonferroni-corrected p<0.01. *** indicates *χ*2 p-value < 0.001. (E) Images of randomly-selected iPS cells expressing *PTEN* variants with feature values in the 2nd decile (top) or 9th decile (bottom) for the three landmark features. Scale bar indicates 2 um. (F) UMAPs of *PTEN* variant morphological profiles in iPS cells colored by landmark features. Blue indicates low and red indicates high z-score versus the synonymous variant distribution, colored as in (G). Scale bar indicates 5 μm. (G) Missense variant effect maps for the PTEN landmark features (grey boxes=missing variants, black dots=synonymous substitutions). Blue to red coloring indicates feature z-score versus the synonymous variant distribution. Positional average of scores (bottom) and P-loop and TI-loop residues (top) are shown. (H) UpSet plot showing the number of *PTEN* missense variants that perturb each combination of landmark features in iPS cells and neurons. Landmark features for each variant were considered perturbed if feature |z-score|>2.5 and KS-test Bonferroni-corrected p<0.01. Sets, represented by bars and dots in the plot, are colored by combinations of impacted features (neither pAKT nor PTEN abundance impacted in both cell types = black, pAKT impacted in both cell types = orange, PTEN abundance impacted in both cell types = teal, both pAKT and PTEN abundance impacted in both cell types = purple). (I) Graphic depicting the relationship between PTEN localization (cell membrane, cytoplasm, or nucleus), activity and protein abundance, annotated with changes to landmark features.

Feature selection led to variant morphological profiles comprising 1,174 feature medians or EMDs for iPS cells and 1,036 feature medians or EMDs^54,55^ for neurons (**Supplementary Fig. 6f**). Features derived from the phalloidin channel in iPS cells and DAPI, mEGFP-PTEN, and pAKT channels in both cell types were most likely to be altered (**Supplementary Fig. 7**). UMAP visualization of the iPS cell and neuron variant morphological profiles demonstrated clear separation of synonymous from 3-nt deletion and nonsense variants (**Fig. 5b**). Likewise, *PTEN* variant morphological impact and AUROC scores measuring distinguishability from WT were higher for 3-nt deletion, frameshift, and nonsense variants than for synonymous and missense variants in both iPS cells and neurons (Mann-Whitney U p<0.001), and replicate impact scores were strongly correlated (Pearson’s r=0.88, 0.89; **Fig. 5c**, **Supplementary Fig. 8a-c**, **Supplementary Table 2**). Morphological impact scores (Pearson’s r=0.95) and AUROC scores (r=0.90) were highly concordant between iPS cells and neurons, suggesting that most variants drove the same overall degree of differences from WT in both cell types (**Supplementary Fig. 8e,f, Supplementary Table 2**). *PTEN* variant features and morphological profiles can be explored at https://visseq.gs.washington.edu/.

### PTEN variants alter abundance, lipid phosphatase activity, and localization in a cell type-specific manner

To explore variant-specific differences between cell types, we focused on three landmark features: Steady-state PTEN abundance as reflected by mEGFP-PTEN intensity, lipid phosphatase activity as reflected by pAKT intensity, and nucleocytoplasmic localization as reflected by DAPI-PTEN pixel-level correlation (**Fig. 5d-f, Supplementary Table 2**). Both PTEN abundance and lipid phosphatase activity form a steep gradient across variants in the UMAP in both iPS cells and neurons (**Fig. 5f**, **Supplementary Fig. 9**). VIS-seq-derived PTEN abundance and activity measurements were correlated with previous multiplexed variant effect measurements of activity in a yeast fitness assay (Pearson’s r=-0.84 versus iPS cell pAKT intensity and -0.86 versus neuron) and abundance in a human cell line (r=0.78 versus iPS cell mEGFP-PTEN intensity, 0.77 versus neuron; **Supplementary Fig. 10**)^5,83^. As expected, abundance-sensitive PTEN positions were concentrated in the core of the protein^69^ (e.g., β-sheet positions 118-121, α-helix positions 137-141; **Fig. 5g, Supplementary Fig. 11**). However, some variants had cell type-specific abundance effects, with 43% of variants having altered PTEN abundance in iPS cells versus 52% in neurons (*χ*^2^ p<0.001; **Fig. 5d**, **Supplementary Fig. 12**). For example, one population of variants was high abundance in neurons and WT-like abundance in iPS cells (**Supplementary Fig. 12g,h**). A second population had moderately reduced abundance in iPS cells and low abundance in neurons (**Supplementary Fig. 12g,h**). Activity-sensitive positions were concentrated at the catalytic surface (e.g., P-loop^69^ positions 123-130 and 171) and in second/third-shell positions (e.g., 155 and 160; **Fig. 5g**, **Supplementary Fig. 11**). Unlike for abundance, variants did not appear to have cell type-specific effects on activity (**Fig. 5d**, **Supplementary Fig. 12a**).

The third landmark feature, DAPI-PTEN correlation, is a sensitive readout of nucleocytoplasmic localization and was strongly correlated with the PTEN nucleus-to-cytoplasm (N/C) intensity ratio (**Supplementary Fig. 12c-e**). WT PTEN is found in the nucleus, where it dephosphorylates nuclear proteins and may have other, non-enzymatic functions^70,72,73,78^. The DAPI-PTEN correlation feature formed a gradient across variants in the UMAP, albeit with less consistency than PTEN abundance or activity (**Fig. 5f**, **Supplementary Fig. 9f**). Many variants had cell type-specific effects on DAPI-PTEN correlation, with 67% having altered DAPI-PTEN correlation in iPS cells versus 48% in neurons (*χ*^2^ p<0.001; **Fig. 5d**, **Supplementary Fig. 12h**). However, PTEN nucleocytoplasmic localization was related to changes in both abundance and lipid phosphatase activity: Low abundance variants had low DAPI-PTEN correlation levels and thus cytoplasmic localization compared to WT, whereas low lipid phosphatase activity variants had high DAPI-PTEN correlation levels and thus nuclear localization (**Fig. 5g**, **Supplementary Fig. 12f-i**). This relationship between localization, abundance and activity obscured the identification of variants that impacted localization alone, so we used regression to remove the effects of both abundance and activity from the DAPI-PTEN correlation feature. The resulting mislocalization score quantified each variant’s effect on localization relative to WT independent of abundance and activity. The mislocalization score enabled us to identify variants that shifted PTEN into the nucleus without altering pAKT levels (**Supplementary Fig. 12j**). We hypothesized that such variants would occur at the membrane-binding positions 163 and 164^84^ and the substrate specificity-determining positions 167 and 168^85^. Indeed, at these positions, 43 (72%) profiled variants had elevated mislocalization scores yet WT-like lipid phosphatase activity (**Fig. 5g**, **Supplementary Figs. 11e,f, 12j**). By contrast, only 18 (27%) variants at four adjacent activity-sensitive positions (165, 166, 170, and 171) had elevated mislocalization scores (**Fig. 5g, Supplementary Figs. 11e,f, 12j**). Thus, analyzing the relationships between multiple interpretable VIS-seq features enabled us to isolate variant effects on specific phenotypes.

Overall, PTEN missense variants impacted diverse combinations of landmark features in human iPS cells and neurons, most frequently altering all three features in both cell types (20%; **Fig. 5h**). Other common combinations included altering DAPI-PTEN correlation in iPS cells only (5.1%); DAPI-PTEN correlation in iPS cells and pAKT intensity in both cell types (7.6%); and abundance and DAPI-PTEN correlation in both cell types (15%). Only 14% of variants had no effect on any landmark features in either cell type. Thus, morphological profiles in combination with simple models enabled us to dissect the relationship between PTEN abundance, lipid phosphatase activity, and nucleocytoplasmic localization in a cell-type-specific manner (**Fig. 5i**).

### PTEN variant morphological profiles are validated by clinical phenotypes and reveal disease-specific variant effects

To validate *PTEN* variant morphological profiles, we curated 5 likely benign (LB) and 62 likely pathogenic or pathogenic (LP/P) missense variants from ClinVar^58^. These clinical control variants were separated on the UMAP in both iPS cells and neurons (**Fig. 6a**, **Supplementary Fig. 13a**). LP/P variant morphological impact scores were significantly elevated as compared to both synonymous and LB/B variant impact scores (Mann-Whitney U p<0.001; **Fig. 6b**, **Supplementary Fig. 13b**). AUROC scores from iPS cells or neurons nearly perfectly separated LP/P variants from a control set composed of gnomAD^86^ variants, outperforming prior yeast activity and human cell abundance variant effect data (**Fig. 6c**, **Supplementary Fig. 13c**). To explore the relationship between *PTEN* variant morphological profiles and autism spectrum disorder/developmental delay (ASD/DD), PTEN Hamartoma Tumor Syndrome (PHTS), or somatic cancer, we curated 66 germline variants in 541 probands from 170 publications for PHTS and ASD/DD phenotypes (**Supplementary Table 3**) as well as 22 variants from the Catalogue Of Somatic Variants In Cancer (COSMIC) database^87^ (**Fig. 6d**). 31 variants in this combined set were associated with only ASD/DD, PHTS or somatic cancer, and 41 were associated with multiple clinical phenotypes. AUROC scores separated curated variants associated with all three clinical phenotypes from gnomAD^86^ controls better than yeast fitness or human cell abundance scores (**Fig. 6c**, **Supplementary Fig. 13c**). Additionally, morphological impact scores in both iPS cells and neurons were significantly increased for variants associated with each clinical phenotype when compared to synonymous or gnomAD^86^ variants (Mann-Whitney U p<0.001; **Fig. 6e**, **Supplementary Fig. 13d**). However, variants associated with ASD/DD had significantly lower morphological impact scores compared to variants associated with PHTS or somatic cancer (Mann-Whitney U p<0.01-0.001). Thus, morphological profiles were superior to single phenotype measurements in capturing overall *PTEN* missense variant pathogenicity and in parsing the relationship between variants and clinical phenotypes.

**Figure 6:**
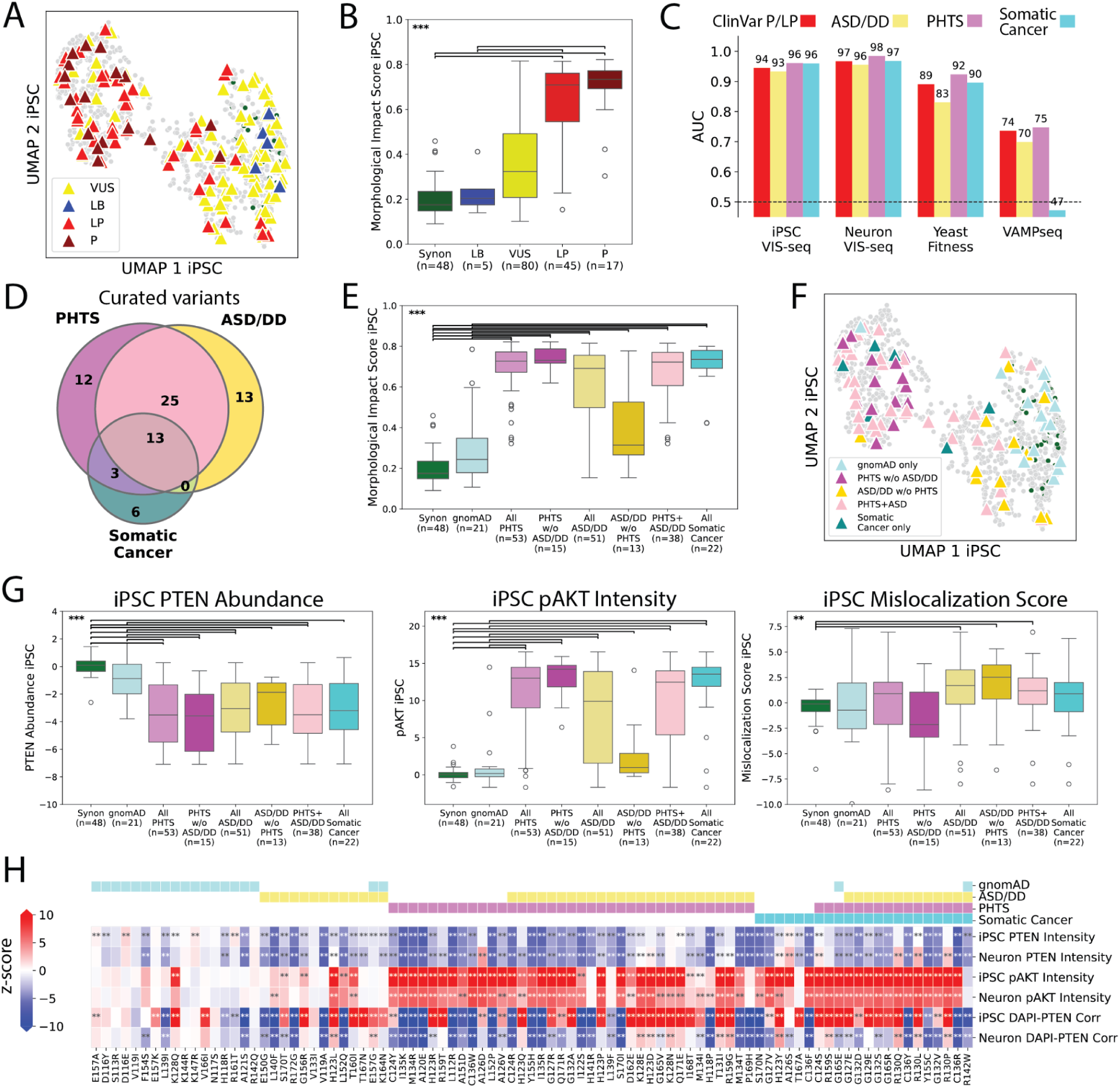
P*T*EN variant morphological profiles predict clinical phenotypes and reveal disease-specific variant effects. (A) UMAP of *PTEN* iPS cell variant profiles from (5B). Triangles indicate missense variants in ClinVar classified as likely benign (LB, blue), likely pathogenic (LP, red), pathogenic (P, dark red), or variant of uncertain significance (VUS, yellow). Circles represent other variants, colored green (synonymous) or grey (non-synonymous). (B) Morphological impact scores for *PTEN* variants in iPS cells plotted by ClinVar class, colored as in (A). Synonymous variants (Synon; green) are included for comparison. (C) AUC are shown for predictions from univariate zero-shot models discriminating between ClinVar LP/P variants or curated clinical variants associated with each phenotype (ASD=autism spectrum disorder, DD=developmental delay, PHTS=*PTEN* hamartoma tumor syndrome) and gnomAD v4.1 variants. VIS-seq models are based on morphological-profile derived AUROC scores (2C) from iPS cell or neuron VIS-seq, and are compared to yeast fitness scores^83^ or VAMPseq scores^5^. (D) Venn diagram of clinical phenotypes associated with curated variants (see Methods). Each clinical phenotype was associated with a variant if it occurred in at least one proband with that variant. (E) Morphological impact scores for *PTEN* variants in iPS cells plotted by association with clinical phenotypes (colors match (D)). gnomAD v4.1 (blue) and synonymous (green) variants are plotted for comparison. *** indicates Mann-Whitney U p<0.001. (F) UMAP of iPS cell *PTEN* variant profiles from (5B). Triangles indicate association with clinical phenotypes (colors match (D)). gnomAD v4.1 (blue) variants are also plotted. Circles represent other variants, colored green (synonymous) or grey (non-synonymous). (G) Boxplots of PTEN abundance as read out by mEGFP-PTEN intensity (left), pAKT intensity (center) and mislocalization (right, see Methods) z-scores versus the synonymous variant distribution for *PTEN* variants in iPS cells. Clinical phenotype category coloring matches (D). gnomAD v4.1 (blue) and synonymous (green) variants are included for comparison. *** indicates Mann-Whitney U p<0.001 and ** indicates p<0.01. (H) Heatmap of *PTEN* variant effects on iPS cell and neuron landmark feature z-scores, annotated by each variant’s association with clinical phenotypes (top). Blue indicates low and red indicates high z-score versus the synonymous variant distribution for each feature. ** indicates KS-test Bonferroni-corrected p<0.01.

We hypothesized that, rather than being points on a single dimension of pathogenicity, ASD/DD, PHTS, and cancer-associated variants might instead perturb distinct molecular and cellular phenotypes. Indeed, variants associated with each clinical phenotype occupied distinct regions of the UMAP, implying that they had disparate molecular and cellular effects (**Fig. 6f, Supplementary Fig. 13e**). Examination of variant effects on landmark features revealed that nearly all disease-associated variants significantly decreased PTEN abundance, with ASD/DD variants producing the smallest decreases^5,88^ (Mann-Whitney U p<0.001; **Fig. 6g, Supplementary Fig. 13f**). Likewise, most disease-associated variants significantly increased pAKT intensity, with ASD/DD variants producing the smallest increases^83,89^ (Mann-Whitney U p<0.001; **Fig. 6g**, **Supplementary Fig. 13f**).

Unlike for PTEN abundance and pAKT intensity, we found a nucleocytoplasmic localization defect enriched among ASD/DD-associated variants, which shifted aberrantly into the nucleus in iPS cells (Mann-Whitney U p<0.01; **Fig. 6g**). ASD/DD-associated variants have been previously described as hypomorphic^89,90^, and they do have lower overall morphological impact as well as milder abundance and lipid phosphatase activity phenotypes compared to PHTS and somatic cancer-associated variants. But, most profiled ASD/DD-associated variants had elevated mislocalization scores that distinguished them from PHTS and cancer-associated variants on the molecular level. Based on this observation, we trained a model that could accurately discriminate between gnomAD control, PHTS-associated, and ASD/DD-associated variants given the landmark features (VIS-seq macro-averaged AUROC=0.92 versus 0.76 for yeast fitness, 0.62 human cell abundance; **Supplementary Fig. 13h**). Thus, we dissected the relationship between *PTEN* variant molecular, cellular, and clinical phenotypes, identifying features that differentiate PHTS-from ASD/DD-associated variants and revealing that ASD/DD-associated variants cannot be understood solely as hypomorphic.

### Variant effect predictors perform poorly on many molecular and cellular phenotypes

State-of-the-art variant effect predictors leverage conservation and protein structure to make accurate inferences of overall variant effects. We compared three leading predictors, AlphaMissense^91^, EVE^92^, and REVEL^93^, to our morphological profiles. Variant effect predictions correlated modestly with lamin A morphological impact scores as compared to impact score replicability (Pearson’s r=0.33 to 0.36 for impact score versus predictions, r=0.88 impact score replicability; **Supplementary Fig. 2h**, **Supplementary Fig. 14a,b**) and more strongly with PTEN impact scores (Pearson’s r=0.69 to 0.83 for impact score versus predictions in iPS cells and 0.62 to 0.77 in neurons; r=0.88 for impact score replicability in iPS cells and 0.89 in neurons; **Supplementary Fig. 8a**, **Supplementary Fig. 14c-f**). These correlations are similar to those observed in large analyses of the relationship between variant effect experimental data and predictions^91–94^.

However, the correlation between individual lamin A or PTEN landmark features and predictions was more variable and poorer. For example, predictions were moderately correlated with lamin A variant abundance (Pearson’s r=-0.36 to -0.47) but poorly correlated with lamin A variant aggregation as assessed by mEGFP-lamin A granularity (r=0.19 to 0.21), and with function as assessed by nuclear circularity (r=-0.27 to -0.39; **Supplementary Fig. 14a**). Similarly, predictions were moderately correlated with PTEN lipid phosphatase activity as assessed by pAKT intensity (Pearson’s r=0.56 to 0.72) but poorly correlated with PTEN abundance (r=-0.27 to -0.48) or PTEN localization (r=-0.09 to -0.32; **Supplementary Fig. 14c,e**). AlphaMissense predictions performed similarly to VIS-seq AUROC scores in separating *PTEN* pathogenic variants from gnomAD controls (**Supplemental Fig. 13c**). However, one-dimensional AlphaMissense scores performed poorly compared to VIS-seq landmark features at discriminating between gnomAD control, PHTS-associated, and ASD/DD-associated variants (VIS-seq macro-averaged AUROC=0.92 versus 0.76 for AlphaMissense; **Supplemental Fig. 13h**). These results highlight the strengths and limitations of current predictors and illustrate how the multimodal data provided by VIS-seq could be used to develop or constrain a next generation of phenotype-aware predictors.

## DISCUSSION

We introduce Variant *in situ* Sequencing (VIS-seq), a scalable single-cell method that uses *in situ* sequencing of circular RNA barcodes to simultaneously measure the effects of thousands of protein variants on features derived from cell images. VIS-seq includes an expression cassette engineered to prevent silencing and enable transgene assays in specialized or hard-to-transduce cell types where silencing and low expression limit current approaches^30^. Thus, VIS-seq expands multiplexed assays of variant effect beyond simple cell growth, protein abundance or reporter readouts in cancer cell lines or non-human cells by enabling morphological profiling that captures numerous biologically relevant, spatially-resolved molecular and cellular phenotypes.

We used VIS-seq to create morphological profiles for ∼3,000 variants in *LMNA* and *PTEN*, genes we chose for their complex relationship to disease. Variant morphological profiles for these genes comprise a high-dimensional space which can be interpreted using multiple landmark features with biological meaning. For example, *LMNA* variant profiles related lamin A dimer and multimer assembly, abundance and aggregation to nuclear shape, enabling discovery of a subset of gain-of-function variants that increase nuclear circularity. *PTEN* variant profiles in both human iPS cells and derived neurons mapped the interdependence of PTEN abundance, lipid phosphatase activity, and nucleocytoplasmic localization, revealing a mislocalization phenotype enriched among autism-associated variants. Morphological profiles, which comprise a large set of simple measurements of the intensity, distribution and shape of different markers in cells, outperformed single phenotype measurements at discerning variant pathogenicity. They also illuminated how variant effects cascade from molecules to subcellular structures to cells in a fashion not captured by current experimental or predictive methods. Thus, our work shows that variant effects span a multidimensional continuum rather than a pathogenic-benign binary or a one-dimensional spectrum of function and highlights the power of multimodal variant effect assays.

Because VIS-seq can measure a vast array of imageable phenotypes in human cell lines and iPS-derived cell types, it will be broadly applicable, particularly to the ∼5,400 disease-associated human genes^95^. While other generalizable multiplexed assays of variant effect can measure cell growth^3,4,96^, protein abundance^5,6^ or transcriptomic changes^97,98^, none offer the spatial information or multimodality of VIS-seq. For example, most genes are incompatible with cell growth assays^99–101^, and growth assays do not generally provide mechanistic insight. Loss of protein abundance is a primary mechanism of missense variant pathogenicity^5,6,102,103^, but fails to capture variants that perturb function via other mechanisms like mislocalization or loss of catalytic activity^104^. The transcriptome is important, but missense variants do not always produce large transcriptomic changes, particularly in utilitarian cancer cell lines^97^, and often exert their primary effect on proteins or cells. VIS-seq leverages the extraordinary catalog of imaging-compatible stains and affinity reagents to overcome these limitations by enabling measurement of a vast swath of molecular phenotypes like protein abundance, mislocalization, and protein function as well as organelle and cell phenotypes. Thus, because many aspects of variant dysfunction can be made visible in cells, VIS-seq can be applied broadly.

Despite its promise, VIS-seq has some important limitations. Although our platform provides robust expression in many cell types, such exogenous expression systems create the possibility of expression-related artifacts. We used fluorescent protein fusions to visualize *LMNA* and *PTEN*, which is known to change variant stability and other phenotypes^105^. The choice of markers and stains is critical, because they determine the content of the morphological profile. Customization of these markers to the protein’s putative function or the cell type of interest allows deeper and more informative morphological profiling, but requires optimization of each new marker. Lastly, automated image analysis remains a difficult problem. For example, while image-derived features are useful for morphological profile clustering, these features require manual interpretation.

These limitations suggest productive avenues for future work. VIS-seq could be combined with time-lapse imaging or 3D organoid models^106^ to capture dynamic or tissue-like phenotypes. Incorporating protein-tagging strategies for endogenous loci^107,108^ with barcodes in the 3’-UTR would mitigate expression-related artifacts. Untaggable proteins could be immunostained for visualization. And, with rapidly advancing deep-learning-based pipelines for embedding image phenotypes^109,110^, the bottleneck of measuring and interpreting morphology at scale will continue to diminish. In summary, VIS-seq represents a significant step forward in our capacity to assess variant effects on complex phenotypes. By uniting multiplexed assays with the power of imaging, our work serves as a blueprint for mapping how variant effects propagate from molecules to cells.

## METHODS

### Cell Culture Reagents

DMEM (Gibco, 11965-092)

DMEM/F-12 (Gibco, 11320-033)

Fetal Bovine Serum (FBS; Hyclone, SH30071.03)

Penicillin/Streptomycin (Pen/Strep; Gibco, 15140-122)

Sodium Pyruvate (Gibco, 11360-070)

OPTI-MEM (Gibco, 31985-070)

DPBS (Gibco, 14190-144)

Versene (Gibco, 15400-054)

Trypsin-EDTA, 0.05%, Phenol-Red (Gibco, 25300-054)

TransIT-293 (Mirus, MIR 2700)

Dimethyl Sulfoxide (DMSO; Sigma, D8418-50mL)

CryoStor CS10 Cell Freezing Medium (STEMCELL Technologies, 100-1061)

Matrigel Matrix (Corning, 354234)

mTeSR Plus (STEMCELL Technologies, 100-0276)

ROCK inhibitor Y-27632 (SelleckChem, S1049)

Lonza 4D Nucleofector SE Kit S (Lonza, V4XC-1032)

Lonza 4D Nucleofector P3 Kit S (Lonza, V4XP-3032)

Lonza 4D Nucleofector P3 Kit X (Lonza, V4XP-3024)

SpCas9 2NLS Nuclease (Synthego, 300 pmol)

Puromycin Dihydrochloride (Gibco, A11138-03)

0.45 μm Syringe Filter (VWR, 76479-012)

### Neuron Differentiation Reagents

Poly-L-ornithine (Sigma, P3655)

Accutase (STEMCELL Technologies, 07922)

Doxycycline hyclate (Sigma, D9891)

Recombinant Human NT-3 (Peprotech, 450-03)

Recombinant Human BDNF (Peprotech, 450-02)

GlutaMAX Supplement 100X (Gibco 35050-061)

MEM Non-Essential Amino Acid Solution 100X (Gibco 11140-050)

B-27 Supplement 50X (Gibco 17504-044)

N-2 Supplement 100X (Gibco 17502-048)

KnockOut DMEM/F-12 (Gibco 12660-012)

Neurobasal-A Medium (Gibco 12349-015)

Laminin Mouse Protein (Gibco 23017-015)

Anti-NCAM1 Rabbit mAb conjugated to Alexa647 (CellSignal, 50831S)

### Cardiomyocyte Differentiation Reagents

RPMI (Invitrogen, 11875135)

DMEM, no glucose (Gibco, 11966-025)

Sodium-L-Lactate (Sigma, 71718)

Vitronectin XF (STEMCELL Technologies, 100-0763)

CHIR-99021 (TOCRIS, 4423)

Wnt-C59 (Selleck, S7037)

Bovine Serum Albumin (BSA; Sigma, A9418)

Ascorbic Acid (Sigma, A8960)

B27 + Insulin (Invitrogen, 17504044)

TrypLE Select 10X (Gibco, A12177-01)

### VIS-seq Reagents

Paraformaldehyde 20% Solution (Electron Microscopy Sciences, 15713-S)

Triton-X 100 (Sigma, Y8787-50ML)

Tween-20 (Sigma, P1279-500ML)

Glutaraldehyde 25% Solution (Electron Microscopy Sciences, 16210)

Deoxynucleotide (dNTP)

Solution Mix 10mM (New England Biolabs, N0447L)

Ultrapure BSA (Thermo Scientific, AM2616)

Ribolock RNase Inhibitor (Thermo Scientific, EO0381)

Revertaid H Minus Reverse Transcriptase (Thermo Scientific, EP0451)

5X Reaction Buffer for RT (Thermo Scientific, EP0451)

Ampligase Buffer (Biosearch Technologies, SS000015-D3)

Taq-IT polymerase (Qiagen, P7620L)

Ampligase (Biosearch Technologies, E0001-100D2)

Rnase H (New England Biolabs, M0297L)

Phi29 DNA Polymerase (Thermo Scientific, EP0094)

Glycerol (Fisher Scientific, G33-1)

Formamide (Fisher Scientific, BP227-500)

Nuclease-Free Water (Thomas Scientific, C001X09)

20X SSC (Invitrogen, 15557036)

10X HBSS (Gibco, 14065-056)

Bovine Serum Albumin (Sigma, A2153-100G)

Sodium Azide (Sigma, S-8032)

Ascorbic Acid (Sigma, A92902-100G)

Illumina MiSeq Incorporation Buffer (Illumina, MS-103-1003)

Illumina MiSeq v2 Nano 500 kit (Illumina, MS-103-1003)

Glass 6-well Plate (Cellvis, P06-1.5H-N)

4’,6-diamidino-2-phenylindole (DAPI; Fisher Scientific, D1306)

Sodium borohydride (Sigma, 452882)

Phalloidin-CF568 conjugate (Biotium 00044-T)

Wheat Germ Agglutinin (WGA)-CF555 conjugate (Biotium 29076-1)

Concanavalin A-Alexa Fluor 750 conjugate (Thermo Scientific, C56127)

Anti-MAP2 Mouse mAb conjugated to Alexa Fluor 594 (BioLegend, 801802)

Anti-phospho-AKT (Thr308)

Rabbit mAb (CellSignal, 13038T)

Goat Anti-Rabbit IgG H&L cross-adsorbed secondary-Alexa Fluor 647 conjugate

(Thermo Scientific, A-21244)

### Cloning Reagents

Corning Untreated 245mm Square BioAssay Dishes (Fisher Scientific, 07-200-600)

Carbenicillin Disodium, USP Grade (GoldBio, C-103-250)

Kanamycin (Sigma, K1537-25G)

NEB 10-beta electrocompetent cells (New England Biolabs, C3020K)

Q5 High-Fidelity 2X Master Mix (New England Biolabs, M0492L)

Klenow Fragment, 3’-5’ exo-minus (New England Biolabs, M0212L)

NEBuilder HiFi DNA Assembly Master Mix (New England Biolabs, E2621L)

T4 DNA Ligase (New England Biolabs, M0202M)

BsaI-HF-v2 (New England Biolabs, R3733L)

SapI (New England Biolabs, R0569L)

Esp3I (New England Biolabs, R0734L)

BsmBI-v2 (New England Biolabs, R0739S)

NheI-HF (New England Biolabs, R3131L)

MluI-HF (New England Biolabs, R3198S)

EcoRI-HF (New England Biolabs, R3101S)

NotI-HF (New England Biolabs, R3189S)

SacII (New England Biolabs, R0157S)

SmrtBell Prep Kit 3.0 (Pacific Biosciences, 102-182-700)

### Western Blot Reagents

RIPA buffer (Abcam, ab156034)

Halt protease inhibitor cocktail (ThermoFisher, 1860932)

BCA assay (ThermoFisher Pierce BCA protein assay kit, 23225)

Laemmeli sample buffer (BioRad, 1610737)

β-mercaptoethanol (BioRad, 1610710)

8–16% Criterion TGX Stain-Free Protein Gel (BioRad, 5678103)

Tris/Glycine/SDS Running Buffer (BioRad, 1610732)

Trans-Blot Turbo Midi 0.2 µm PVDF Transfer Packs (BioRad, 1704157)

Bovine Serum Albumin (BSA; Sigma-Aldrich, A9418)

Anti-phospho-AKT (Thr308) Rabbit mAb (CellSignal, 13038T)

Anti-pan-AKT mouse mAb (Cell Signaling, 2920)

Anti-PTEN rabbit mAb (Cell Signaling, 9559)

Anti-lamin A/C mouse mAb (BioLegend, 600001)

StarBright blue 520 goat anti-mouse IgG (BioRad, 12005866)

StarBright blue 700 goat anti-rabbit IgG (BioRad, 12004162)

hFAB rhodamine anti-actin primary antibody (BioRad, 12004164)

### Plasmids

pHSG299 (Takara Bio, 3299)

pHR-UCOE-SFFV-Zim3-dCas9-P2A-EGFP (Addgene, 188899)

piggyFlex (Addgene, 218234)

lentiGuide-Puro (Addgene, 52963)

pCAG-NLS-HA-Bxb1 (Addgene, 51271)

Hyperactive *piggyBac* transposase expression vector^111^ (hyPBase; gift from Jay Shendure lab)

pMD2.G (Addgene, 12259)

psPAX2 (Addgene, 12260)

### Cloning Oligos

mEGFP-HBBIVS2trunc-IRES-v1/v2N/v2C gene fragments (Twist, inquire for full sequence)

PD-BsmBIsites gene fragment (Twist):

gggagggggcgggaaaccgcctaaccatgccgagtgcggccgcGAGAAGGTCGGGTCCAGATATTGTAACTG TACGAAGACTGGGAGACGAGTGCCTGCAGGCATACGTCTCTCGATACTTGTTCGATCCTTC TAAGCTTGGCGTAACTAGATCTTGAGACTAGCTTTAAGGCCGGTCCTAGCAACGTATCTGTC GAGTAGAGTGTGGGCTCGTGGccgcggtcggcgtggactgtagaacactgccaatgccggtcccaagcccggataa aagtggagggtacags

Prelamin A site-saturation library (positions 178-273; Twist SSVL, inquire for full sequence)

PTEN site-saturation oligo library (positions 112-172; Twist, inquire for full sequence)

PTEN_tile3_s1/2 gene blocks (Twist, inquire for full sequence)

H1.4 oligo library (Twist, inquire for full sequence)

H1.4_s1/2 gene blocks (Twist, inquire for full sequence)

RPS19 oligo library (Twist, inquire for full sequence)

RPS19_s1/2 gene blocks (Twist, inquire for full sequence)

VISseq_GG_BColigo (IDT):

GTATAGTTTGTGCGGTGGTCCGTCTCGACTGNNNNNNNNNNNNCGATCGAGACGCGAAGG TGTAGGGGATTGAT

VISseq_GG_BColigoX2 (IDT): ACTTCCGTCTCGACTGNNNNNNNNCGATACTTGTTCGATCCTTCTAAGCTTGGCGTAACTAG ATCTTGATATCCAACTGTACGAAGACTGNNNNNNNNCGATCGAGACGCGAAC

Lenti_GG_BColigo (IDT): ACTCGTCTCGGCTTAACTGTACGAAGACTGNNNNNNNNNNNNCGATACTTGTTCGATCCTT CTAAGCTTGGCGTAACTAGATCTTGACTGCGAGACGCTA

*LMNA*-KO gRNA (Synthego): GGAGCUCAAUGAUCGCUUGG-(gRNA scaffold)

*PTEN*-KO gRNA (Synthego): CCAAUUCAGGACCCACACGA-(gRNA scaffold)

### VIS-seq Oligos

BestMIP1 (IDT): /5Phos/CGATACTTGTTCGATCCTTCacctcttcCTCCTtcaactcctccctcccaacatactctcctccctACTGTA CGAAGACTG

BestMIP2_3ntdel (IDT): /5Phos/CGATACTTGTTCGATCCTcaacctcttcATctcttaatccttacccaccatacctccctccattccacctccacAC TGTACGAAGACTG

CROP_RT2_LNA_5Am (IDT): /5AmMC12/C+AA+GA+TC+TA+GT+TA+CGCCAAGCTTA

ISS1 (IDT): tccctcccaacatactctcctccctACTGTACGAAGACTG

ISS2 (IDT): tacctccctccattccacctccacACTGTACGAAGACTG Mitoprobe 12S (IDT): /5Cy5/TACTGTTTGCATTAATAAATTAA Mitoprobe 16S (IDT): /5Cy5/CTCTATATAAATGCGTAGGG

### VIS-seq expression construct cloning

VIS-seq expression constructs for expressing N-terminal mEGFP-tagged prelamin A (PBv2b-laminA) and PTEN (PBv2c-NSapI-PTEN) libraries were prepared by Gibson assembly^112^ using NEBuilder HiFi DNA Assembly Master Mix (New England Biolabs, E2621L) followed by Golden-Gate^113^ insertion of barcodes and then variant libraries. These plasmids are being made available for collaborators on Addgene.

First, an intermediate plasmid was cloned which contains BsmBI sites for installation of a padlock-BC sequence in a future step. We cloned this plasmid by digesting piggyFlex^23^ (Addgene, 218234) with NotI-HF (New England Biolabs, R3189S) and SacII (New England Biolabs, R0157S) and performing Gibson assembly with the PD-BsmBIsites gene fragment (Twist). Then, the PBv2b expression plasmid was prepared by Gibson assembly of the following five fragments: intermediate backbone from the previous step digested with NheI-HF (New England Biolabs, R3131L) and MluI-HF (New England Biolabs, R3198S), UCOE amplified from pHR-UCOE-SFFV-Zim3-dCas9-P2A-EGFP^114^ (Addgene, 188899), EF1a promoter/intron and PuroR-WPRE-SV40 terminator fragments amplified from lentiGuide-Puro^115^ (Addgene, 52963), and a gene fragment from Twist containing mEGFP, MluI and EcoRI restriction sites, a truncated portion of the *HBB* intron 2, and IRES (mEGFP-HBBIVS2trunc-IRES-v1, Twist). All PCR amplifications were performed using Q5 Master Mix (New England Biolabs, M0492L), using 98C for the denaturation temperature, 72C for the extension temperature, and 60C for the annealing temperature and were pulled prior to saturation.

PBv2b-NSapI, PBv2b-CSapI, and PBv2c-NSapI expression plasmids were cloned using similar Gibson assemblies. For all three plasmids, a gene fragment was used containing SapI sites either upstream (C-terminal tag: PBv2b-CSapI) or downstream (N-terminal tag: PBv2b-NSapI) of mEGFP^31^ (mEGFP-HBBIVS2trunc-IRES-v2N/C respectively, Twist). For the PBv2c-NSapI plasmid, we substituted a CAG promoter amplified from pCAG-NLS-HA-Bxb1^116^ for the EF1a fragment as well. All VIS-seq backbone plasmids (PBv2b, PBv2b-NSapI, PBv2b-CSapI, and PBv2c-NSapI) were barcoded using Golden-Gate assembly. Klenow fill-in (New England Biolabs, M0212L) of VISseq_GG_BColigo (IDT) yielded a double strand oligo with a 12bp degenerate barcode used for PBv2b barcoding (all plasmids), and PCR of VISseq_GG_BColigoX2 (IDT) with primers attaching BsmBI sites yielded a double strand oligo used for PBv2c-NSapI barcoding (with two 8bp barcodes). The barcoding reaction was performed in eight simultaneous reactions each with 500ng of input plasmid, a 2:1 molar ratio of barcode, 1.5 μL of Esp3I (New England Biolabs, R0734L), 1 μL of T4 ligase (New England Biolabs, M0202M), and 2.5 μL of 10X T4 ligase buffer with water added to 25 μL. These reactions were cycled between 37 °C and 16 °C 45 times, post-digested with 1μL of BsmBI-v2 (New England Biolabs, R0739S) and then pooled, concentrated and electroporated into NEB 10-beta electrocompetent *E. coli* (New England Biolabs, C3020K). Bacteria were plated onto BioAssay dishes (Fisher Scientific, 07-200-600) containing LB + 100 μg/mL carbenicillin (GoldBio, C-103-250), and then scraped and prepped the following day. 3 electroporations were used for each plate. Each barcoded VIS-seq expression vector had a minimum of 25 million colonies per plate for 2 plates, which were pooled 1:1.

The *LMNA* (prelamin A sequence, derived from NM_170707.4) variant cDNA library was purchased from Twist as an SSVL library and delivered in an arrayed format; positions 178 to 273 were mixed together to generate the library used in this paper. *PTEN* (positions 112-172, derived from NM_000314.8), *RPS19* (positions 52-109, derived from NM_001022.4), and *HIST1H1E* (positions 54-113, derived from NM_005321.3) variant cDNA libraries were cloned in-house using BsaI Golden-Gate assembly, and then inserted into the VIS-seq expression plasmids with SapI Golden-Gate assembly, similar to a previously published method^117^. First, the plasmid pHSG299 (Takara Bio, 3299) was modified to contain two BsaI sites. The oligo libraries containing a tile of gene variants were amplified from the purchased Twist oligo pool and then assembled with the flanking gene blocks coding for the remainder of the gene cDNA (PTEN_tile3_s1/2 or RPS19_s1/2 or H1.4_s1/2 from Twist) and the modified pHSG299-BsaI. This Golden-Gate reaction was performed using 100 ng of pHSG299-BsaI, a 2:1 molar ratio of gene blocks (Twist) to backbone, a 2:1 molar ratio of the amplified oligo library to backbone, 1μL of BsaI-HF-v2 (New England Biolabs, R3733L), 0.5 μL of T4 ligase, and 2.5 μL of 10X T4 ligase buffer with water added to 25 μL total volume. WT H1.4 cDNA was cloned by substituting the amplified oligo tile with a wildtype gene block (Twist). These reactions were incubated at 37 °C for 18 hrs, and then concentrated and electroporated into NEB 10-beta electrocompetent *E. coli*, followed by overnight grow-out in liquid culture and prep.

Lastly, full cDNA variant libraries were inserted into corresponding VIS-seq expression plasmids. The prelamin A variant library (positions 178 to 273; Twist) and barcoded PBv2b from the prior step were digested with MluI-HF and EcoRI-HF (New England Biolabs, R3101S) and then ligated using 0.5 μL T4 ligase at 16 °C overnight in T4 ligase buffer at a final concentration of 1X in a 20 μL reaction. Other genes were assembled into barcoded PBv2b-NSapI (*HIST1H1E*), PBv2b-CSapI (*RPS19*), or PBv2c-NSapI (*PTEN*) using Golden-Gate assembly: 100ng of barcoded plasmid and a 2:1 molar ratio of variant library plasmid, 1.5 μL of SapI (New England Biolabs, R0569L), 0.5 μL of T4 ligase, and 2.5 μL of 10X T4 ligase buffer in a 25 μL aqueous reaction followed by cycling between 16 °C and 37 °C for 3 hours. After assembly, reactions were post-digested with an additional 1 μL of SapI.

All reactions were concentrated and then electroporated into NEB 10-beta electrocompetent *E. coli* followed by growth of dilutions in LB overnight. For both libraries, the dilution that corresponded most closely to ∼20-fold barcode coverage over variants was selected the following day and prepped.

### PacBio sequencing

Barcoded PBv2b-Prelamin A and PBv2c-NSapI-PTEN libraries were digested with NotI-HF. Then, the SmrtBell Prep Kit (SmrtBell Prep Kit 3.0, Pacific Biosciences, 102-182-700) instructions were followed to generate PacBio libraries. Libraries were sequenced using the Pacific Biosciences Revio sequencer. Pacybara^118^ was used to process reads to generate variant-barcode association tables (*LMNA*) or variant-double barcode association tables (*PTEN*).

### Cell passaging and plating

U2OS cells were cultured in media composed of DMEM (Gibco, 11965-092) with 10% fetal bovine serum (FBS; Hyclone, SH30071.03) and 1% Pen/Strep (Gibco, 15140-122). Cells were passaged in the following way: lifted by washing with DPBS (Gibco, 14190-144) and then incubating with Trypsin 0.25% (Gibco, 25300-054) for 15 minutes, then Trypsin quenched with media and cells were spun down at 300g for 5 minutes, and lastly cells were then resuspended in media and replated. For VIS-seq experiments, cells were plated at a density of 800k per well on all wells of a glass-bottom 6-well plate (Cellvis, P06-1.5H-N) and fixed ∼12 hrs after plating. iPS cells were cultured on matrigel-coated six-well plates in complete mTeSR Plus (STEMCELL Technologies, 100-0276) media. The media was changed at least every other day. The plates were coated by diluting matrigel (Corning, 354234) to a final concentration of 80 μg/mL in DMEM-F12 (Gibco, 11320-033), adding 1 mL to each well (or 5mL to a 10-cm plate), and incubating the plates at room temperature for 1 hour. iPS cells were passaged in the following way: lifted by washing each well with DPBS and then incubating each well with Versene (Gibco, 15400-054) for 3-5 minutes until cells began to lift off the plate, then cells were resuspended in mTeSR Plus with 10 μM Y-27632 (SelleckChem, S1049) and replated. Media changes without re-plating used mTeSR Plus only. For VIS-seq experiments, iPS cells were plated on all wells of a matrigel-coated glass-bottom 6-well plate (Cellvis, P06-1.5H-N) at 600k per well and fixed 6 hours after they were plated.

### CRISPR/Cas9 knockout and cell-line preparation

For knockout of *LMNA/C,* U2OS C11 cells^119^ were electroporated on the Lonza 4D Nucleofector using protocol CM104 in a single well of a Lonza 4D-Nucleofector SE Kit S (Lonza, V4XC-1032). The electroporation mix contained 1 million cells resuspended in 30 μL of supplemented Lonza SE buffer with 1 μL of SpCas9 (Synthego) and 1 μL of *LMNA*-KO gRNA (Synthego). Cells were then recovered and seeded into 96-well plates at a dilution of 1 cell per 5 wells. Clones were grown over a month, passaged, and *LMNA*-knockout was confirmed by Western blot (see below). U2OS C3 *LMNA*-KO was the clone selected for both replicates of VIS-seq on *LMNA* variants.

For knockout of *PTEN,* NWTC11.G3-WT^120^ iPS cells were electroporated on the Lonza 4D Nucleofector using protocol CM137 in a single well of a Lonza 4D-Nucleofector P3 Kit S (Lonza, V4XP-3032). The electroporation mix contained 200,000 cells per well resuspended in 30 μL of supplemented Lonza P3 buffer with 1 μL of SpCas9 and 1 μL of *PTEN*-KO gRNA (Synthego). Cells were then recovered and seeded into 96-well plates at a dilution of 1 cell per 5 wells. Clones were grown over 10 days, passaged, and then *PTEN*-knockout was confirmed by Western blot. *PTEN*-KO C18 was used for the first replicate of iPS cell and neuron VIS-seq on *PTEN* variants, and *PTEN*-KO C2 was used for second replicates.

### Low-MOI piggyBac integration of VIS-seq libraries

For low-MOI integration of the VIS-seq *LMNA* library, U2OS C3 *LMNA*-KO cells were electroporated using the Lonza 4D Nucleofector with protocol CM104 in 6 separate wells of a Lonza 4D-Nucleofector SE Kit S. Each cuvette contained 1 million cells per well resuspended in 30 μL of supplemented Lonza SE buffer with 200 ng of VIS-seq *LMNA* variant library and 50ng of hyPBase^111^ expression plasmid. Cells were then recovered in a 6-well plate in media (DMEM + 10% FBS + 1% Pen/Strep), using one well per electroporation well, for 4 days and then pooled and integrants were selected with 2ug/mL puromycin (Gibco, A11138-03) for 7 days. After selection, cells were recovered for at least 2 days prior to freezing in media with 5% v/v added DMSO (Sigma, D8418-50 mL). Library-expressing cells were thawed, washed and resuspended in media and recovered for 2 days prior to VIS-seq experiments.

For low-MOI integration of VIS-seq libraries into iPS cells, cells were electroporated in 2 cuvettes of Lonza 4D-Nucleofector P3 Kit X (Lonza, V4XP-3024) using the Lonza 4D Nucleofector with protocol CM104. The *PTEN* library was integrated into *PTEN*-null clones of NWTC11.G3-WT^120^ and other libraries were integrated into WTC11 iPS cells. Each cuvette contained 3 million cells resuspended in 100 μL of supplemented Lonza P3 buffer with 400 ng of VIS-seq expression plasmid and 100 ng of hyPBase^111^ expression plasmid. Cells were then recovered for 4 days in a 10-cm plate per cuvette and then pooled and integrants were selected with 1ug/mL puromycin for 3 days or until cells were >90% mEGFP expressing. After selection, cells were recovered for at least 2 days in mTeSR Plus and then frozen in liquid nitrogen after resuspension in CryoStor CS10 Cell Freezing Medium (STEMCELL Technologies, 100-1061). Library-expressing cells were thawed, washed and resuspended in mTeSR Plus with 10 μM Y-27632 and recovered for 2 days prior to VIS-seq experiments or neuronal differentiation.

### Neuronal differentiation of iPSC using Dox-inducible NGN2 expression

To differentiate NWTC11.G3-WT^120^ iPS cells into neurons using doxycycline-inducible NGN2 expression, we followed a protocol reported previously^81^. On day -3, iPS cells were dissociated using Versene and then resuspended in Pre-Differentiation medium with 10μM Y-27632. Pre-Differentiation medium contains: KnockOut DMEM/F-12 (Gibco 12660-012), 1X MEM Non-Essential Amino Acids (Gibco 11140-050), 1X N-2 Supplement (Gibco 17502-048), 10ng/mL NT-3 (Peprotech, 450-03) and BDNF (Peprotech, 450-02), 1 μg/mL Laminin protein (Gibco 23017-015), and 2 μg/mL doxycycline (Sigma, D9891). Cells were plated at 800k/well using 1mL of media per well in 12-well plates coated in matrigel, and media was changed to Pre-Differentiation medium on days -2 and -1. On day -1, a glass-bottom 6-well plate was coated using 15 μg/mL poly-L-ornithine (Sigma, P3655) overnight and then washed with DPBS 3 times the following day followed by air drying at room temperature. On day 0, pre-differentiated cells were dissociated using Accutase (STEMCELL Technologies, 07922) and then pelleted at 200 g for 5 minutes, followed by resuspension in Maturation medium. Maturation medium contains: 50% Neurobasal-A medium base (Gibco 12349-015), 50% DMEM/F-12 base, 1X MEM Non-Essential Amino Acids, 0.5X GlutaMAX Supplement (Gibco 35050-061), 0.5X N-2 Supplement, 0.5X B-27 Supplement (Gibco 17504-044), 10 ng/mL NT-3 and BDNF, 1 μg/mL Laminin protein, and 2 μg/mL doxycycline. Cells were plated at 600k/well into poly-L-ornithine coated 6-well plates in 2 mL of Maturation medium per well. Neurons were fixed and processed for VIS-seq on day 4 (day 7 after doxycycline addition).

To confirm differentiation, neurons were fixed (see below, VIS-seq neuron fixation protocol) and stained using DAPI (Fisher Scientific, D1306) and rabbit anti-NCAM1 primary antibody conjugated to Alexa647 (CellSignal, 50831S) using the VIS-seq staining mix (see below, Cell Painting Buffer + 0.75% Triton-X 100) and imaged using the same settings as used in the VIS-seq phenotype imaging (**Supplementary Fig. 1d**).

### Cardiomyocyte differentiation of iPS cells

WTC11 iPS cells were cultured on 10 μg/ml Vitronectin XF (STEMCELL Technologies, 100-0763) prior to cardiomyocyte differentiation. Small molecule directed differentiation was performed as previously described with some modifications^20^. iPS cells were plated in 15 μg/ml Vitronectin XF-coated 24 well plates at 7.5×10^4^ cells per well in mTeSR Plus supplemented with 10 μM Y-27632. Media was changed to fresh mTeSR Plus without 10 μM Y-27632 the next day. Directed differentiation was initiated (D0) when cells reached 60-70% confluency by aspirating mTeSR Plus and replacing media with RBA media: RPMI (Invitrogen, 11875135), 0.5 mg/mL bovine serum albumin (Sigma, A9418), 0.213 mg/mL ascorbic acid (Sigma, A8960) supplemented with 4.5 μM CHIR-99021 (TOCRIS, 4423). After two days (D2), CHIR-99021 containing media was removed and replaced with RBA supplemented with 2 μM Wnt-C59 (Selleck, S7037). On D4, Wnt-C59-containing media was replaced with RBA. On D6 (and every other day afterwards), media was replaced with cardiomyocyte media: RPMI, B27 plus insulin (Invitrogen, 17504044). Cardiomyocyte cultures typically begin beating by D6. After D12, cardiomyocyte media was removed and cardiomyocytes were dissociated to single cells with TrypLE Select 10X (Gibco, A12177-01) and replated on Matrigel-coated 6 well plates at 3×10^6^ cells per well in cardiomyocyte media supplemented with 10% FBS and 10 μM Y-27632. At D20, cardiomyocytes were purified by replacing cardiomyocyte media with lactate selection media for four days: DMEM, no glucose (Gibco, 11966-025) with 4mM sodium-L-lactate (Sigma, 71718). Lastly, cardiomyocytes were dissociated to single cells using TrypLE Select 10X on day 56 (D56) after start of differentiation and then mEGFP-positivity was measured by flow cytometry (see end of next section).

### VIS-seq expression cassette silencing and *in situ* sequencing compared to lentiviral construct

The lentiviral plasmid pLenti_EF1a_mEGFP-H1.4 was cloned via a Golden-Gate assembly of four fragments: backbone derived from PCR of lentiGuide-Puro^115^, EF1a-mEGFP-H1.4 fragment from amplifying the assembled PBv2b-NSapI-H1.4(WT) plasmid (see VIS-seq Cloning), a double stranded 12-bp barcode flanked by padlock sequences generated by Klenow fill-in (see above, Lenti_GG_BColigo (IDT)), and an IRES-Puro-WPRE-SV40 terminator fragment from amplifying the assembled PBv2b-NSapI-H1.4(WT) plasmid. The plasmid was subsequently sequence-confirmed and prepped.

To generate lentivirus, HEK293T cells cultured in DMEM supplemented with 10% FBS, Pen/Strep, and sodium pyruvate (Gibco, 11360-070), were seeded into a 6 well plate at 900,000 cells per well in the evening. The next morning, TransIT-293 (Mirus, MIR 2700) suspensions were made by adding 0.625 μg pMD2.G (Addgene, 12259), 1.25 μg psPAX2 (Addgene, 12260), and 0.625 μg pLenti_EF1a_mEGFP-H1.4 to 250 μl OPTI-MEM (Gibco, 31985-070) for every well to be transfected. After vortexing, 7.5 μl Transit reagent was added per well and mixed by gently flicking the tube, followed by a 15 minute incubation at room temperature. After the incubation, the suspension was distributed dropwise onto the plated cells and allowed to sit for 24 hours. The following day the media was replaced with fresh DMEM and placed back in the incubator for 48 hours. Following the incubation, the media was removed from the cells and spun at 1000 g for 5 minutes. The supernatant was filtered through a 0.45 μm syringe filter (VWR, 76479-012) and 1 mL aliquots frozen at -80 °C until use.

To generate iPS cells with integrated lentivirus expressing mEGFP-H1.4 and linear RNA barcodes, 10 μL of filtered lentiviral supernatant was added to a 6-well plate of WTC11 iPS cells plated at a concentration of 100,000 per well for two replicates, followed by puromycin selection after 4 days until >90% of cells were GFP-positive. To generate iPS cells with integrated VIS-seq cassette expressing mEGFP-H1.4 and circular RNA barcodes, the protocol for low-MOI integration of the VIS-seq plasmid PBv2b-NSapI-H1.4(WT) was followed (see above) for two replicates.

After selection, both iPS cell populations were passaged and monitored for mEGFP-positivity using the BD FACSSymphony A3 Analyzer over 14 days. Cells were taken to flow cytometry on days 0 (immediately after selection), 4, 9 and 14. mEGFP expression was measured for at least 10,000 cells using the 488 nm laser and the BB515 filter. Nontransduced iPS cells were used as a control to set the mEGFP-positive gate.

Day 14 transduced and VIS-seq cassette-expressing iPS cell populations were fixed, *in situ* libraries were generated, and the first base of *in situ* sequencing was performed (see below for protocol). Only mEGFP-expressing cells were analyzed and STARCall was used to count the number of rolling circle colonies (rolonies) for each condition.

Cardiomyocytes were differentiated from transduced and VIS-seq-expressing iPS cell populations after selection (see above for protocol). On day 56 after start of differentiation, mEGFP-positivity of cardiomyocytes was measured using the BD Symphony S6 Cell Sorter with 488 nm laser and BB515 filter.

### Western blots

Approximately 1 million cells per sample was incubated with 50 μl 1x RIPA buffer (Abcam, ab156034) supplemented with Halt protease inhibitor cocktail (ThermoFisher, 1860932) for 30 minutes on ice. Samples were spun down at 10,000g and the supernatant moved to new chilled tubes. The supernatants were assayed by BCA assay (ThermoFisher, Pierce BCA protein assay kit 23225) to determine the total protein concentration. Lysates were normalized and mixed with 2x Laemmeli sample buffer (BioRad, 1610737) supplemented with fresh β-mercaptoethanol (BioRad, 1610710) and boiled for 5 minutes at 95 °C. Boiled samples are spun at 10,000g for 1 minute right before loading 30μg total protein per well for each sample into 8–16% Criterion TGX Stain-Free Protein Gel (BioRad, 5678103). Gels were run with 10x Tris/Glycine/SDS Running Buffer (BioRad, 1610732) diluted to 1x with type I ultrapure water for 35 minutes at 200V. After running, the gels were imaged for total protein on a BioRad Chemidoc MP. Transfer was done using Trans-Blot Turbo Midi 0.2 µm PVDF Transfer Packs (BioRad, 1704157) on the Trans-Blot Turbo Transfer System using the TURBO settings. Membrane blocking was done in 5% BSA (Sigma-Aldrich, A9418) in Tris buffered saline with Tween 20 (TBST; 50mM Tris-HCL, 150mM NaCl, 0.05% Tween-20) for rocking for 1 hour at room temperature. Primary antibody staining was done in TBST with 1% BSA rocking overnight at 4°C. Primary antibodies used were anti-phospho-(Thr308)-AKT rabbit mAb (Cell Signaling, 13038), anti-pan-AKT mouse mAb (Cell Signaling,2920), anti-PTEN rabbit mAb (Cell Signaling, 9559), and anti-lamin A/C mouse mAb (BioLegend, 600001) all used at a 1:1000 dilution from stock. Following primary, blots were washed three times for 10 minutes with TBST rocking at room temperature. Secondary staining was done in TBST with 0.01% SDS and 1% BSA. Secondary antibodies used were StarBright blue 520 goat anti-mouse IgG (BioRad, 12005866) and StarBright blue 700 goat anti-rabbit IgG (BioRad, 12004162) both used at a 1:5000 dilution. hFAB rhodamine anti-actin primary antibody (BioRad, 12004164) was used to stain for beta-actin as a loading control in the secondary solution at a dilution of 1:5000. Secondary staining was done on a rocker for one hour at room temperature followed by three 10 minute washes in TBST and developing on the BioRad Chemidoc MP.

### VIS-seq molecular biology

*In situ* sequencing protocols are similar to those published previously^33^. However, we detail them here, documenting the changes we made. All *in situ* sequencing experiments were performed using all wells of a glass-bottom #1.5H 6-well plate with cells plated at 800,000 cells per well (U2OS) or 600,000 cells per well (iPS cell/neuron), and unless indicated otherwise wash steps contain 2mL per well.

U2OS and iPS cells were first washed 3 times with DPBS prior to fixation. Then 1 mL of 4% PFA (Electron Microscopy Sciences, 15713-S) in DPBS was added to each well. Neurons were fixed in media, by removing 50% of media (leaving 1 mL remaining in well) and adding 250 μL of 20% PFA dropwise for a final concentration of 4% PFA, followed by swirling gently to mix PFA into media. All cells were incubated during fixation in the dark for 30 minutes at room temperature and then washed 3X with DPBS prior to enzyme steps.

Cells were permeabilized with 1 mL per well of 0.1% Triton-X 100 (Sigma, Y8787-50ML) in DPBS for 15 minutes at room temperature. This was followed by 2 washes of 0.1% Tween-20 (Sigma, P1279-500ML) in DPBS (henceforth called PBS-T). A 5’-amino-linked RT primer containing linked nucleic acid (LNA) bases was then hybridized by incubation with 600 μL per well of 1 μM CROP_RT2_LNA_5Am^7,9^ (IDT) for 15 minutes at room temperature followed by 3 PBS-T washes. Then cells were fixed with 1 mL per well of 3% PFA and 0.1% glutaraldehyde (Electron Microscopy Sciences, 16210) in PBS-T for 30 minutes at room temperature, followed by 3 PBS-T washes. 750 μL per well of reverse transcription reaction mix (Table 1) was added to each well. Water was added to the spaces between wells and the plate was sealed with foil. The plate was incubated at 37 °C while shaking at 80 RPM overnight.

**Table 1.**

| Reagent | Volume (μL) |
| --- | --- |
| Revertaid RT buffer (5x) | 150 |
| dNTPs (10 mM, New England Biolabs, N0447L) | 18.75 |
| Ultrapure BSA (50 mg/mL, Thermo Scientific, AM2616) | 6 |
| RiboLock (40 U/μL, Thermo Scientific, EO0381) | 9.375 |
| Revertaid enzyme (200 U/μL, Thermo Scientific, EP0451) | 18 |
| Nuclease-free water (Thomas Scientific, C001X09) | 547.875 |
| *total* | 750 |

Following reverse transcription, cells were washed three times with PBS-T and then fixed with 1 mL per well of 3% PFA and 0.1% glutaraldehyde in PBS-T for 30 minutes at room temperature. Cells were then washed five times with PBS-T to reduce the dNTP concentration. A previously reported optimized padlock probe^9^ is annealed onto the sample, the gap in the padlock probe is filled in, and the circle is ligated shut using 600 μL per well of the gap-fill mix described in Table 2. The plate was incubated at 37 °C for 5 minutes and then 45 °C for 90 minutes.

**Table 2.**

| Reagent | Volume (μL) |
| --- | --- |
| Ampligase buffer (10X, Biosearch Technologies, SS000015-D3) | 60 |
| Ultrapure BSA (50mg/mL) | 2.4 |
| Padlock probe (BESTMIP1 (IDT) for *PTEN* or BESTMIP2_3ntdel (IDT) for *LMNA*) at 10 μM | 6 |
| Taq-IT polymerase (50 U/μL, Qiagen, P7620L) | 0.24 |
| dNTP mix (10 μM, prepare fresh dilution each time) | 3 |
| Ampligase (5 U/μL, Biosearch Technologies, E0001-100D2) | 30 |
| RNaseH (5 U/μL, New England Biolabs, M0297L) | 12 |
| Nuclease-free water | 486.36 |
| *total* | 600 |

The cells are washed twice with PBS-T and then 600 μL of RCA mix (Table 3) are added to each well. The plate was incubated at 30 °C while shaking at 80 RPM overnight.

**Table 3.**

| Reagent | Volume (μL) |
| --- | --- |
| Phi29 Buffer (10X) | 60 |
| dNTPs (10 mM) | 15 |
| Ultrapure BSA (50 mg/mL) | 2.4 |
| Glycerol (50%, Fisher Scientific, G33-1) | 60 |
| Phi29 enzyme (10 U/μL, Thermo Scientific, EP0094) | 15 |
| Nuclease-free water | 447.6 |
| *total* | 600 |

The cells were washed twice with PBS-T and then stained. Different staining mixes were used for U2OS cells, iPS cells, and neurons. U2OS cells were first stained with 600 μL of mitoprobe staining mix^10^, a FISH-based alternative to MitoTracker (Table 4) per well. They were incubated at 37 °C for 20 minutes.

**Table 4.**
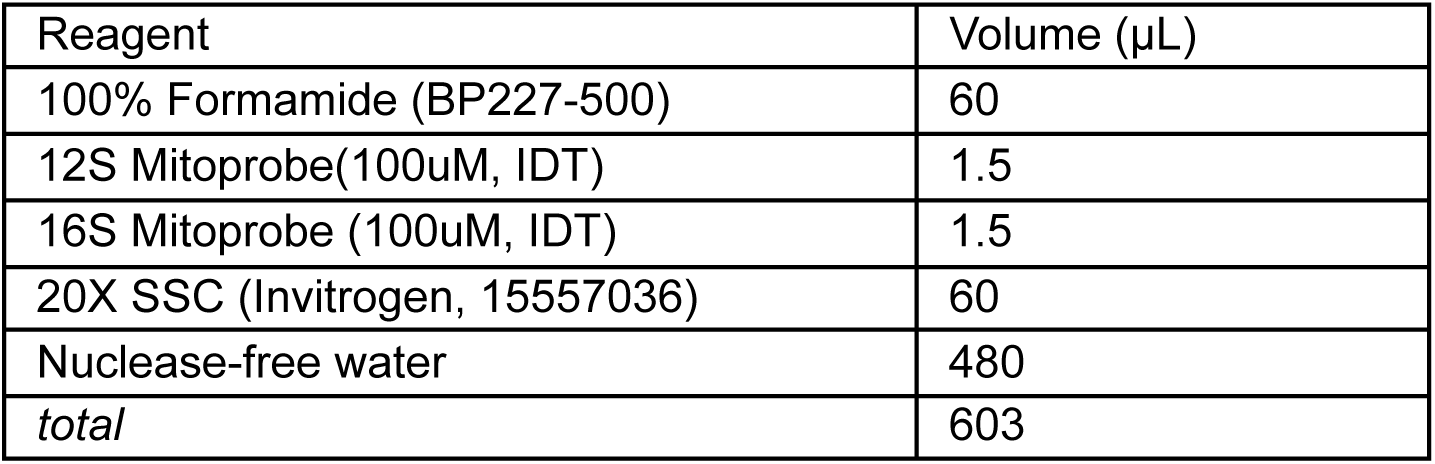

Following mitoprobe staining, U2OS cells were then washed twice with PBS-T and 600 μL of Cell Painting^12^ mix (Table 5) was added to each well. Cells were incubated at room temperature for 30 minutes. Then U2OS cells were washed five times with PBS-T.

**Table 5.**

| Reagent | Volume (μL) |
| --- | --- |
| DAPI (200 ng/μL, Fisher Scientific, D1306) | 0.6 |
| WGA CF555 (Biotium 29076-1) | 0.9 |
| Phalloidin CF568 (Biotium 00044-T) | 0.75 |
| RiboLock (40 U/μL) | 4.5 |
| Cell Painting Buffer: 1X HBSS (Gibco, 14065-056) + 1% BSA (Sigma-Aldrich, A2153-100G) + 0.01% NaN<sub>3</sub> (Sigma, S-8032) | 595 |
| *total* | 601.75 |

iPS cells and neurons were first stained with a modified cell painting/immunostaining mix that included concanavalin-A and an anti-phospho-AKT rabbit antibody and then stained with a secondary goat anti-rabbit Alexa-647-labeled antibody (Thermo Scientific, A-21244). The cell painting mix for iPS cells (Table 6) did not contain WGA and the cell painting mix for neurons (Table 7) did not include WGA or phalloidin and instead contained an anti-MAP-2 Alexa594-labeled primary antibody (BioLegend, 801802). The cell painting/immunostaining mix was incubated at room temperature while shaking at 80 RPM for 1 hour.

**Table 6.**

| Reagent | Volume (μL) |
| --- | --- |
| DAPI | 0.75 |
| Concanavalin A AF750 (Thermo Scientific, C56127) | 0.75 |
| Phalloidin CF568 | 0.9375 |
| anti-(Thr308)-pAKT Antibody (CellSignal, 13038T) | 1.5 |
| 20% Triton-X 100 | 9.375 |
| Cell Painting Buffer: 1X HBSS + 1% BSA + 0.01% NaN<sub>3</sub> | 750 |
| *total* | 763.3125 |

**Table 7.**

| Reagent | Volume (μL) |
| --- | --- |
| DAPI | 0.75 |
| Concanavalin A AF750 | 0.75 |
| anti-MAP-2 Antibody AF594 (BioLegend, 801802) | 1.5 |
| anti-(Thr308)-pAKT Antibody | 1.5 |
| 20% Triton-X 100 | 9.375 |
| Cell Painting Buffer: 1X HBSS + 1% BSA + 0.01% NaN<sub>3</sub> | 750 |
| *total* | 763.875 |

After painting and primary staining, iPS cells or neurons were washed three times with PBS-T. Then they were incubated with 750 μL per well of a goat anti-rabbit Alexa-647 secondary antibody at a concentration of 0.5 U/μL in Cell Painting^12^ buffer (1X HBSS + 1% BSA + 0.01% NaN_3_) with 0.75% Triton-X 100 for 1 hour while shaking at 80 RPM. Then cells were washed five times with PBS-T. Cells were kept in the dark during and after staining and prior to imaging.

Once the U2OS cells, iPS cells, and iPS-derived neurons had been stained, 2 mL of imaging mix (Table 8) were added to each well prior to imaging. We used ascorbic acid at a final concentration of 10mM in the imaging mix to reduce photodamage.

**Table 8.**
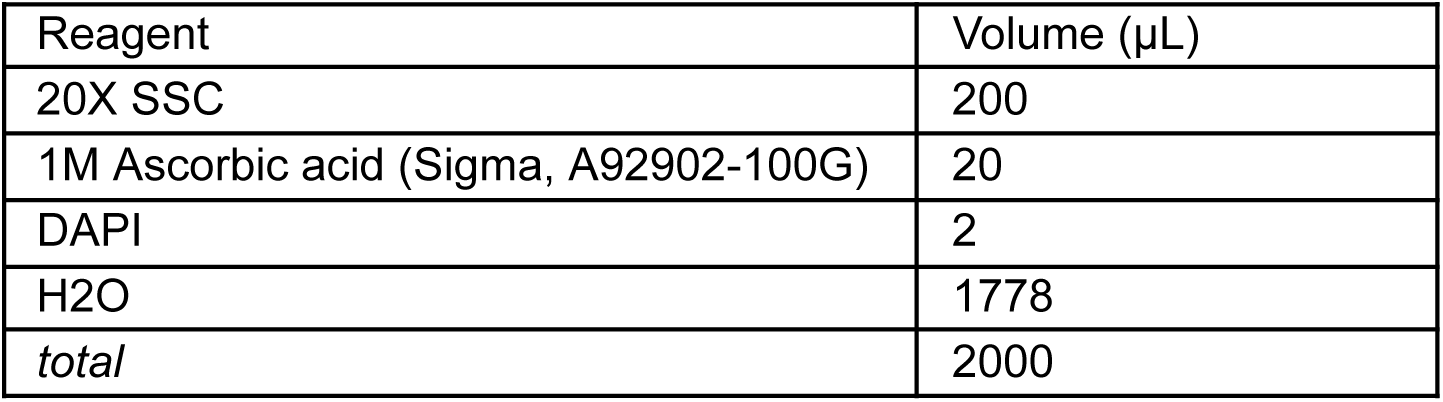

After phenotype imaging, dyes were removed differently for U2OS cells (*LMNA*) or iPS cells and neurons (*PTEN*). U2OS cells were permeabilized with 2 mL 70% ethanol per well for 15 minutes at room temperature, which was sufficient to remove phalloidin; mitoprobe was removed later during sequencing steps by heating^10^. 1 mL PBS-T was added and removed five times to titrate the ethanol concentration below 1%. Then U2OS cells were washed twice with 2 mL PBS-T per well.

iPS cells and neurons were washed twice with 2 mL PBS-T per well prior to chemical bleaching with 1 mL of 1.74 mg/mL sodium borohydride (Sigma, 452882) in DPBS incubated for 15 minutes at room temperature. The borohydride solution was used immediately after mixing. Then, cells were washed three times with PBS-T. After phenotyping dyes were removed or chemically bleached, we proceeded with *in situ* sequencing.

We then added 600 μL of 0.5 μM sequencing primer (using previously reported ISS1 (IDT) for *PTEN* experiments and ISS2 (IDT)^9^ for *LMNA* experiments) diluted in hybridization buffer of 2X SSC + 10% formamide to each well and incubated at room temperature for 30 minutes. We washed cells twice with PBS-T and then once with MiSeq Incorporation Buffer (IB, Illumina, MS-103-1003). We then proceeded to the base cycles: 600 μL of Illumina MiSeq reagent #1 (Illumina, MS-103-1003) was added to each well and the 6-well plate was incubated at 60 °C for 3 minutes. We then added and removed 1 mL IB four times until the incorporation mix was diluted >50-fold. We washed each well with 5X SSC + 0.5% Tween-20 (a wash-step replacement for Incorporation Buffer) and incubated wash steps at 55C (reduced temperature to prevent amplicon unbinding) for 5 minutes while shaking at 300 RPM. We repeated this wash and incubation four more times for a total of 5 washes to remove background base incorporation. We then performed base (genotyping) imaging (see below) in the imaging mix (Table A).

After imaging was completed, we added 700 μL Illumina MiSeq reagent #4 (Illumina, MS-103-1003) to each well and incubated at 60 °C for 6 minutes, washed once with 2 mL IB per well. We then added 2 mL 5X SSC with 0.5% Tween-20 to each well and incubated at 55C for 1 minute while shaking at 300 RPM. We repeated this wash and incubation two more times for a total of 3 washes. We then washed once with 2 mL IB before adding Illumina MiSeq reagent #1 to incorporate another base. We repeated this process until 8 (*PTEN*) or 12 (*LMNA*) bases were imaged.

### VIS-seq imaging

All VIS-seq imaging was performed using a Nikon Ti2-E inverted epifluorescence microscope with a motorized stage and motorized objective and filter cube turrets, similar to previously reported configurations for *in situ* sequencing^33^. A Hamamatsu ORCA-Fusion scientific CMOS camera was used to collect images. A Lumencor Celesta light engine with 7 laser lines (408, 445, 473, 518, 545, 635, and 750 nm) was used for fluorescence illumination. We also used a Finger Lakes Instrumentation HS-625 high speed emission filter wheel with 5 bandpass filters (499-529, 553-577, 604-644, 662-691, and 698-766 nm), and enabled hardware-level triggering between these filters. The phenotype imaging was performed using a 0.75 NA Plan Apochromat Lambda 20X objective, whereas base cycles were performed using a 0.45 NA Plan Apochromat Lambda 10X objective. Laser power was set to 1% for 405nm laser (DAPI), 20% for other phenotyping lasers, and 100% for *in situ* sequencing lasers, and exposure time was maintained at 50ms for both phenotyping and *in situ* sequencing imaging.

### Confocal imaging of protein variant libraries in iPS cells and neurons

Imaging of *PTEN*, *HIST1H1E*, *RPS19* and *LMNA* libraries in iPS cells, neurons, or U2OS cells for **Fig. 1e-g** was performed using a Nikon A1R HD25 laser scanning confocal microscope. 405, 488, and 561 nm Nikon LU-N4 lasers were used to image DAPI-stained, mEGFP-tagged protein libraries, Phalloidin-CF568-stained cells, respectively. Imaging was performed using an Nikon Apochromat Lambda S LWD 40x water-immersion objective (NA 1.15). A Nikon galvano scanner and Nikon A1-DUG-2 GaAsP photomultiplier tubes were used to acquire 1024×1024 12-bit color images of single z-planes.

### Stitching and alignment of VIS-seq images

Stitching, alignment, and read calling was performed using STARCall. Stitching and alignment is required for processing of VIS-seq data due to the small (<3 pixel) features being detected and the need for them to align to one another across all imaging cycles. STARCall stitches the raw microscope images across positions and cycles simultaneously using an extended MIST algorithm^121^ incorporating improvements from ASHLAR^122^.

Using stage positional information, a graph was constructed representing images as nodes with edges connecting every overlapping image pair. This graph contained all imaging cycles, with edges between the same tile in one cycle to the respective tile in all other cycles. For every edge on this graph, both inter-cycle and intra-cycle, we performed the phase cross-correlation algorithm^123^ to find the alignment that maximizes the correlation between the two images. The alignment for each edge was scored by calculating the normalized cross-correlation (NCC) on the overlapping region of the two images. The NCC was computed by normalizing the overlapping pixel intensities to unit vectors and computing their dot product. To remove erroneous offsets, we calculated a lower-bound for the NCC scores of offsets, using the 95th percentile of NCC scores calculated on a random selection of non-overlapping images. Any pair of images that had a NCC score below this threshold was removed from the graph. Removing these pairs caused some portions of the graph to become disconnected, especially if cells in some regions of the well were sparsely plated thus yielding few features for alignment. To reconnect these portions, we trained a linear model on the remaining offsets, predicting their offset given their stage positions. This model was then used to replace edges on disconnected images.

We then constructed a linear system of equations on the global (x,y) positions of the images, where each offset in the graph creates two equations, relating the difference in position of the two images to the offset. This system was then solved using least-squares, finding the optimal global position for each image. In particular, the system of equations was solved minimizing the mean absolute error (MAE), the solution for which can be found quickly with linear programming, with an additional benefit being that the influence of outliers is limited when compared to minimizing the mean squared error. All images were then combined to form composite images for each cycle, with overlapping regions between images combined using a weighted average. The code for STARCall and instructions on how to use it can be found on GitHub (see Data and code availability section).

### Read calling of genotyping images

We defined the base reads as bright, high frequency features present in only one of the channels and changing frequently across bases, best characterized by a small point spread function. In addition, we characterized common image features that should be filtered out: low-intensity cell background, diffuse features that are present in most channels and increase monotonically across cycles, and cell debris, which present as extremely bright, high frequency features present in all channels and not fluctuating between cycles. With the objects of interest characterized, we developed a procedure for selecting only the fluorescent reads similar to previous approaches used for *in situ* sequencing datasets^7^: First we subtracted a small Gaussian blur (*σ*=4), which left only small, high frequency features, filtering out general and cell background. We then z-score normalized the resulting image, which corrected for differences between channels in intensity. Next we subtracted the second-maximal channel on a per-pixel basis, which did not affect the fluorescent dots present in any one channel but greatly reduced the intensity of cell background and cell debris present in all channels. After these filtering steps we applied a small (*σ*=1) Gaussian blur to smooth out noise we introduced. Finally we calculated the standard deviation on a per-pixel level across imaging cycles, which resulted in higher values for features that change between bases, filtering out background and debris that doesn’t change between cycles. After summing across channels we obtained a grayscale image containing the features we wanted to extract and believed to only be fluorescent reads. The Laplacian of Gaussian blob detection algorithm was applied to find these features at the expected size (1≤*σ*≤3). The positions of the blobs were used across all cycles to extract the base reads, which are defined using the highest channel intensity in each cycle for that position. We extracted these values from the images after we subtracted the small Gaussian blur and z-score normalized them, additionally applying a dilation morphological filter with a footprint of 2 pixels to ensure the maximum value of each dot is sampled. We associated barcodes (*LMNA*) or pairs of barcodes (*PTEN*) to the nearest genotype, making no call if two genotypes are equally close by edit distance. Later, we filtered cells on their least edit distance barcode.

### Segmentation of cells and nuclei

Cell segmentation was performed using Cellpose version 2.2.1^36^ for cell border segmentation and StarDist version 0.8.5^37^ for nuclear segmentation. CellPose requires the estimated diameter of cells to be specified to rescale images before inference. The method for cell segmentation may be adjusted depending on the cell type and imaging setup. We used diameter values of d=50 for U2OS cells and d=35 for iPS cells and neurons. For U2OS cells we provided DAPI as the nuclear channel and WGA + Phalloidin as the cytoplasmic channel to CellPose. For iPS cells we provided DAPI as the nuclear channel and WGA as the cytoplasmic channel. Due to difficulties segmenting neurons, we used only nuclear segmentation, converting it into an estimated cell segmentation by expanding the nuclear labels out to neighboring cells, with a maximum expansion distance of 25 pixels.

### Intensity normalization and feature extraction

We performed intensity normalization for each well’s phenotype imaging separately to adjust for uneven illumination of the field of view by subtracting the 5th percentile value of that pixel across all such fields. After stitching and registering phenotype images with genotypes, we then partitioned each well’s phenotype image into a 20-by-20 grid and passed images, cell segmentations from Cellpose^36^, and nuclear segmentations from StarDist^37^ into CellProfiler version 4.2.6^38^ for all grid positions that have genotyped cells. We then merged all of these grid positions to generate a well-level cells-by-features matrix. In the case of the *PTEN* VIS-seq experiments, we adjusted the pAKT channel (stained with AF647) for bleed-through from the ConA channel (stained with AF750) by subtraction. The CellProfiler^38^ pipelines for feature extraction used in *PTEN* and *LMNA* VIS-seq experiments are available in the STARCall GitHub (see Data and code availability section).

### Generation and analysis of variant-level morphological profiles

We filtered out cells whose consensus barcodes were >1 (*LMNA*) or >2 (*PTEN*) edit distance from a barcode in the barcode-variant lookup table, had no variant called, contained barcodes (*LMNA*) or combinations of 2 barcodes (*PTEN*) that were present in <10 cells, or contained variants with <5 barcodes. We then extracted variant-level feature median and earth-mover distance (EMD) values^54,55^ from WT for all variants and features. We filtered out feature EMDs which were non-reproducible in a manner similar to Pearson *et al*^55^. Briefly, we randomly partitioned WT cells into two batches and computed scores equal to EMD values between the batches for each feature normalized to median absolute deviation. After repeating this 25 procedure times, we removed features above an average score threshold of 1.5 times the interquartile range plus the lower quartile average score.

We then used pycytominer version 1.2.1^56^ to perform variant-level median aggregation and variant-level median/EMD normalization and selection, for generating morphological profiles. This normalization was a z-scoring for each feature median using synonymous feature means and standard deviation in the *LMNA* VIS-seq experiment, and a z-scoring over all variants for the *PTEN* VIS-seq experiments. For feature selection, we removed non-biologically-relevant features like orientational or positional features, and then we ran pycytominer^56^ feature selection using both “correlation threshold” and “variance threshold” to remove features. The features that remained were the variant-level morphological profiles.

Next, we performed principal component analysis (PCA) for dimensionality reduction and uniform manifold approximation and projection (UMAP)^57^ for visualization of morphological profiles. For PCA, we used the minimum number of PCA dimensions that could explain 60% (*PTEN*) or 70% (*LMNA*) of the dataset variance. For *PTEN* neuron profiles, we noticed that three variants with poor inter-replicate correlation were outlying in PC1, and they were removed prior to rerunning PCA and further analysis. We then performed UMAP on PCA-reduced variant profiles using cosine similarity to visualize the local structure of the phenotype manifold.

To generate variant-level feature p-values, we performed KS-testing between each variant and WT for each feature. We then performed Bonferroni correction for multiple hypothesis testing to generate p-values which we used to annotate variants for landmark feature differences. Hit features were defined as features that were differential (at Bonferroni-corrected p<0.001) in at least 25 variants.

To generate morphological impact scores, we computed the cosine similarity between a variant’s morphological profile and the median synonymous variant morphological profile, computed by taking the median over each selected feature. We then defined the morphological impact score as ½ x (1 minus the cosine similarity).

To perform Louvain clustering^124^ on the *LMNA* profiles, we generate an unweighted k-nearest-neighbor graph using k=25 of the PCA-reduced variant profiles using a cosine similarity-based distance and then perform the Louvain algorithm on that graph.

### Training of variant-specific classifiers and AUROC scores

To characterize the morphological distinguishability of each variant from WT using single-cell image data, we trained a collection of binary classifiers, each tasked with distinguishing a single variant from the wild type in a binary classification setting (**Supplementary Fig. 3a**). The input to each variant-specific model was the list of CellProfiler^38^ features, after non-biologically-relevant features were removed (see above), extracted from all wild type cells and the cells corresponding to that variant. The output was a binary label indicating whether each cell was wild type or variant. In order to prepare the training data, we first removed any variants that had fewer than 150 samples, after which we divided the filtered training data into a stratified 8:1:1 train/test/validation split. For each variant, we used the train and validation sets to train a single binary classifier using the Extreme Gradient Boosting (XGBoost) algorithm^125^, where the validation set was used to determine when to stop training based on an early stopping criterion that monitors validation loss. Finally, after training each classifier, we evaluated its performance using the area under the receiver operating characteristic curve (AUROC) on the held-out test set. This test set was not used during training, providing an unbiased estimate of how well each model generalizes to unseen data. We used the resulting AUROC scores to quantify the morphological impact of each variant, as we hypothesize that variants with greater morphological differences from wild type cells would be more easily distinguished by a binary classifier (**Supplementary Tables 1,2**).

### Landmark feature analysis

For *LMNA* landmark feature analysis, Louvain clustering was performed on morphological profiles to generate cluster annotations (see above). Then, Mann-Whitney U-testing was performed on feature-cluster pairs to determine which features best separate each cluster from all other clusters. Both feature medians and EMDs were tested. For each feature-cluster pair, two metrics of effect size were computed. First, a robust z-score equal to the difference between the median (over variants) value of the feature in the tested cluster and the median of all variants, divided by the median absolute deviation over all variants. Second, a robust z-score between the median (over variants) value of the feature in the tested cluster and the median of synonymous variants, divided by the median absolute deviation of synonymous variants. For a given feature-cluster pair, these z-scores summarize the degree to which the feature values differ amongst cluster variants relative to all variants and to synonymous variants. Feature-cluster pairs were filtered by both |robust z-scores|>1.5, yielding ∼3,800 pairs. Lastly, feature-cluster pairs were ranked by increasing Mann-Whitney U p-value, the top 15% were inspected (with p-values<10^-7^), and landmark features were manually selected. The nuclear circularity (Nuclei_AreaShape_FormFactor) feature associated with cluster 9 (containing aggregating variants^39^) was the 1st ranked feature-cluster pair in the analysis. After selection, *LMNA* landmark features for each variant were called as perturbed if feature EMD z-score>2.5 and feature Bonferroni-corrected KS-test p<0.01. EMD scores were used here due to inherent cell-level heterogeneity in landmark features (especially aggregation) for every variant.

For *PTEN*, we selected landmark features based on prior those chosen in previous high-throughput experiments^5,83^ or those highlighted by previous experiments focused on single variants^72,78,126^. *PTEN* landmark features for each variant were called as perturbed if feature median synonymous |z-score|>2.5 and feature Bonferroni-corrected KS-test p<0.01.

### Structural analysis

We used the lamin A structures of A11 tetramer 6JLB^63^ and recently published A22 four-coil interaction^64^, and PTEN structure 1D5R^69^ to perform structural analysis of morphological profiles. Structures were visualized and residues colored with position-averaged feature z-scores using PyMOL v3.1. Lamin A rod subdomains were defined using Ahn *et al.*^63^ and PTEN P-loop and TI-loop annotations were defined using Lee *et al.*^69^

To call lamin A dimer-facing and multimer-facing positions, we plotted the inter-chain minimum beta-carbon to beta-carbon distance for each lamin A position (except glycines) on a given chain. We identified residues with inter-chain local minima distances, and any residues within 1.5 Angstroms of this inter-chain local minimum distance, as interacting with an opposing chain (**Supplementary Fig. 5f**). Interactions were labeled as dimer-facing or multimer-facing depending on the geometry of the chains: coil 1B forms the A11 lateral structure and therefore contains both types of interacting residues whereas coil 2A forms the A22 four-helix bundle and therefore only contains multimer-facing residues.

### Clinical variant curation and pathogenicity classification

All clinical variant analysis excluded variants with predicted splicing impact defined by SpliceAI^59^ scores>0.2, since these variants would not be expected to correlate with our assay. *LMNA* ClinVar^58^ variants were accessed on 6/18/2024 and LP/P variants were curated. Conflicting variants were labeled as variants of uncertain significance (VUS). *PTEN* ClinVar variants were accessed on 12/19/2024; conflicting variants were treated as their most recent classification to generate a putative list of profiled VUS, LB/B and LP/P variants.

*PTEN* missense variants on gnomAD v4.1 were accessed on 5/14/2025. gnomAD v4.1^86^ variants that were not ClinVar LP or P or present in the clinical variant-phenotype associations we curated (see below) were used as controls for either zero-shot prediction of pathogenicity or clinical phenotypes (**Fig. 6c, Supplementary Fig. 13c,h**). Otherwise, all gnomAD v4.1 variants that were not ClinVar LP or P were used (**Fig. 6e-h, Supplementary Fig. 13d-g**).

*PTEN* missense variants present in 10 or more patient cancers on the Catalogue Of Somatic Mutations In Cancer (COSMIC)^87^, accessed on 12/12/2024, were used to define somatic *PTEN* cancer-associated variants (**Fig. 6d**).

For curation of *PTEN* clinical variant-phenotype associations for PHTS and ASD/DD, 170 publications containing a total of 541 probands with *PTEN* missense variants were identified using HGMD and ClinVar (accessed 5/13/2025). The clinical features for each individual with a *PTEN* missense variant were curated manually by a molecular genetic pathologist blinded to the functional assay results (AEM) and included demographics, clinical diagnosis (Cowden Syndrome (including Cowden-like Syndrome), Bannayan-Riley-Ruvalcaba syndrome, or PTEN hamartoma tumor syndrome, and the presence of autism spectrum disorder (ASD) or developmental delay/intellectual disability (DD/ID)). For the analysis, probands reported to have ASD, DD, or ID were considered to be affected by neurodevelopmental phenotype. These phenotypes were combined due to inconsistencies in reporting practices. A large language model (Claude Sonnet 4, Anthropic) was used to assemble the data table (**Supplementary Table 3**), with the output verified manually. Any variant linked to a clinical phenotype in at least one affected proband was then associated with that clinical phenotype (**Fig. 6d**).

Variant-level AUROC scores (VIS-seq, iPS cell and neuron), as well as yeast fitness^83^ scores, VAMP-seq^5^ scores, and AlphaMissense^91^ scores were used for zero-shot classification of pathogenicity. ROC curves and corresponding AUC values from this zero-shot classification are reported in **Fig. 6c, Supplementary Fig. 13c**. For multiclass prediction between gnomAD controls, PHTS variants not associated with ASD/DD, and ASD/DD variants not associated with PHTS, a linear support vector classifier (SVC) was trained on VIS-seq landmark features in iPS cells, or neurons, or variant-level scores used for single phenotype classification (**Fig. 6c, Supplementary Fig. 13c,h**). Then the ROC curves for classification were computed by macro-averaging model sensitivity and specificity over the three classes, treating each class with equal weight (**Supplementary Fig. 13h**).

## Data and code availability

Variant morphological impact scores, AUROC scores and landmark feature values can be found on MaveDB (*LMNA:* https://mavedb.org/experiments/urn:mavedb:00001243-a, *PTEN* iPS cells: https://mavedb.org/experiments/urn:mavedb:00001244-b, and *PTEN* neurons: https://mavedb.org/experiments/urn:mavedb:00001244-a). All code necessary to reproduce our analyses, starting at the cell by features matrix, can be found at https://github.com/FowlerLab/visseq. STARCall code used to generate the cells by features matrix from phenotyping and genotyping images can be found at https://github.com/FowlerLab/starcall-workflow. Code used to generate trained XGBoost models and variant-level AUROC scores can be found at https://github.com/FowlerLab/fisseqtools. Tabular data is provided as a supplement or, for larger files, is available via Zenodo at doi.org/10.5281/zenodo.15787684. Single-cell feature profiles will be made available via Amazon Web Services S3 and raw image data via the BioImage Archive upon publication.

## Acknowledgements

We would like to acknowledge Brian Beliveau, Andrew Stergachis, Hao Yuan Kueh, and Sanjay Srivatsan for their scientific guidance. For assistance with microscopy and executing VIS-seq experiments, we acknowledge Wai Pang-Chan at the University of Washington Biology Imaging Facility as well as Tony Cooke. For assistance with flow cytometry, we acknowledge scientists and staff at the DLMP Flow Cytometry Core at the University of Washington. Funding was provided by the National Institutes of Health (RM1HG010461 to DMF, LMS, and FPR; R35GM152106 to DMF; R01HG013025 to LMS, DMF, AEM, and AFR; K99HL177347 to CEF; R01HL171174 to KCY; R01HL164675 to FPR), Chan Zuckerberg Initiative (CZIF2024-010284 to DMF and LMS; CP-2-1-Fowler to DMF) the Brotman Baty Institute for Precision Medicine (CC28 to AEM), the Department of Veterans Affairs Biomedical Laboratory Research and Development Service (I01BX006428 and IK2BX004642 to KCY), the Novo Nordisk Foundation (DMF). AEM was supported by an Early Career Award from the Alex’s Lemonade Stand for Childhood Cancer and RUNX1 foundation (21-25037).

## Author contributions

Assigned according to the Contributor Role Taxonomy (CRediT), https://credit.niso.org/

Conceptualization: DMF and SP (equal contributors)

Data curation: AEM (lead), AFR, SP

Funding acquisition: DMF, LMS

Formal analysis: GS, NB, SP (lead), WN

Investigation: AJV, CEF, DLH, KAS, KCY, KP, RLP, SF, SP (lead)

Methodology: FPR, HJK, SP (lead)

Software: NB (lead), SP Visualization: GS, NB, SP (lead)

Writing (original draft): DMF and SP (equal contributions)

Writing (review and editing): all authors

## Declaration of interests

FPR is an advisor and shareholder in Constantiam Biosciences.

## SUPPLEMENTAL FIGURES

**Supplementary Figure 1:**
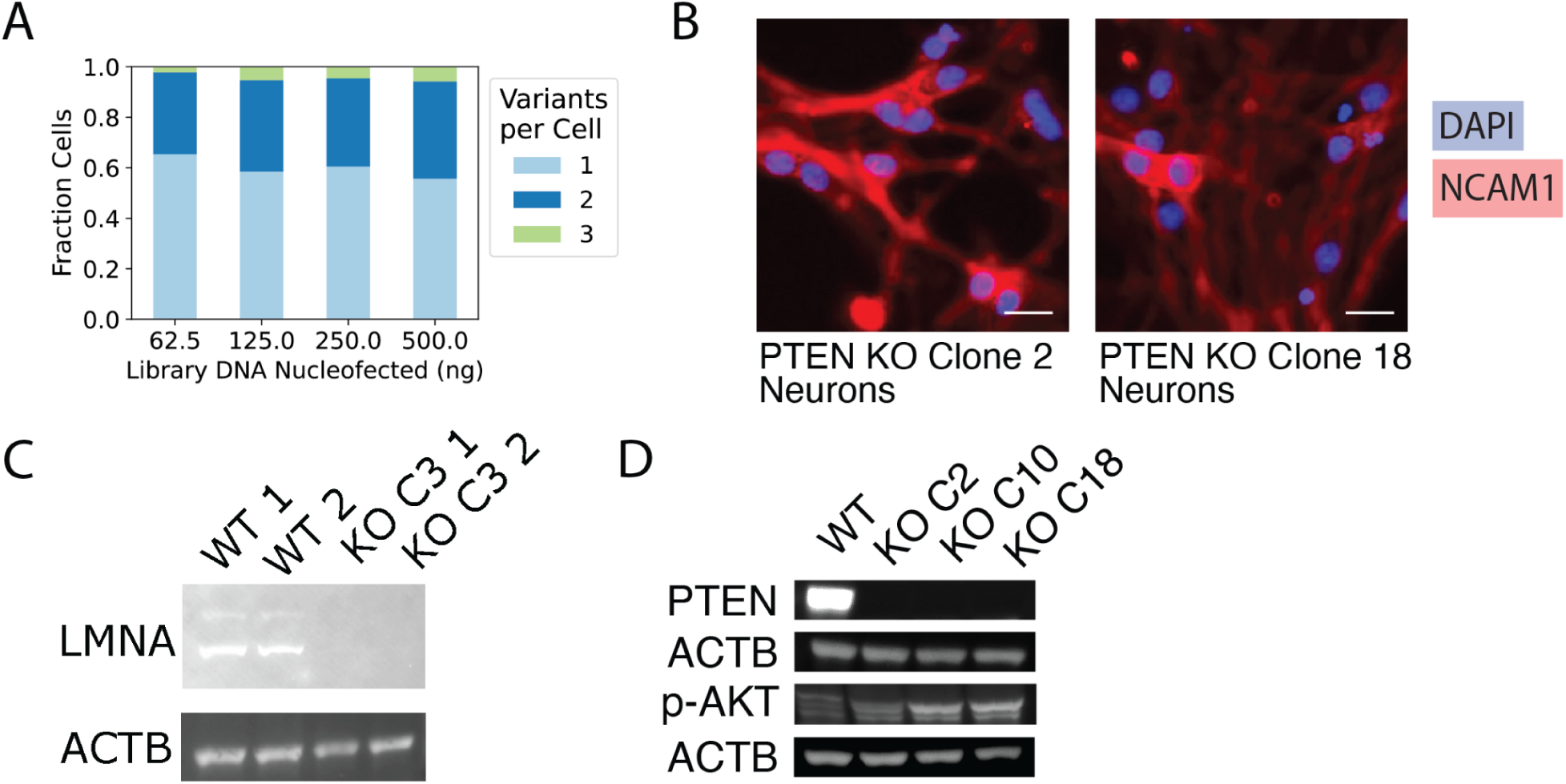
*piggyBac* MOI titration and knockout Western blots. (A) Fraction of cells with 1, 2, or 3 integrations, determined by 4-base *in situ* sequencing, after co-transfection of *LMNA* VIS-seq library DNA at different quantities (in nanograms) with plasmid encoding Piggybac-ase (at a 4-fold lower mass). (B) Imaging of day-7 neurons derived from clonal NGN2-inducible *PTEN* knockout lines stained for DAPI (blue) and NCAM1 (red). Scale bar indicates 20 μm. (C) Western blot of U2OS cells showing parental line (left) and *LMNA* knockout lines (right) stained for lamin A protein. (D) Western blot of NGN2-inducible iPS cells showing parental line (left) and *PTEN* knockout lines (right) stained for PTEN and phospho-AKT protein. Clonal lines C2 and C18 were used for *PTEN* VIS-seq experiments.

**Supplementary Figure 2:**
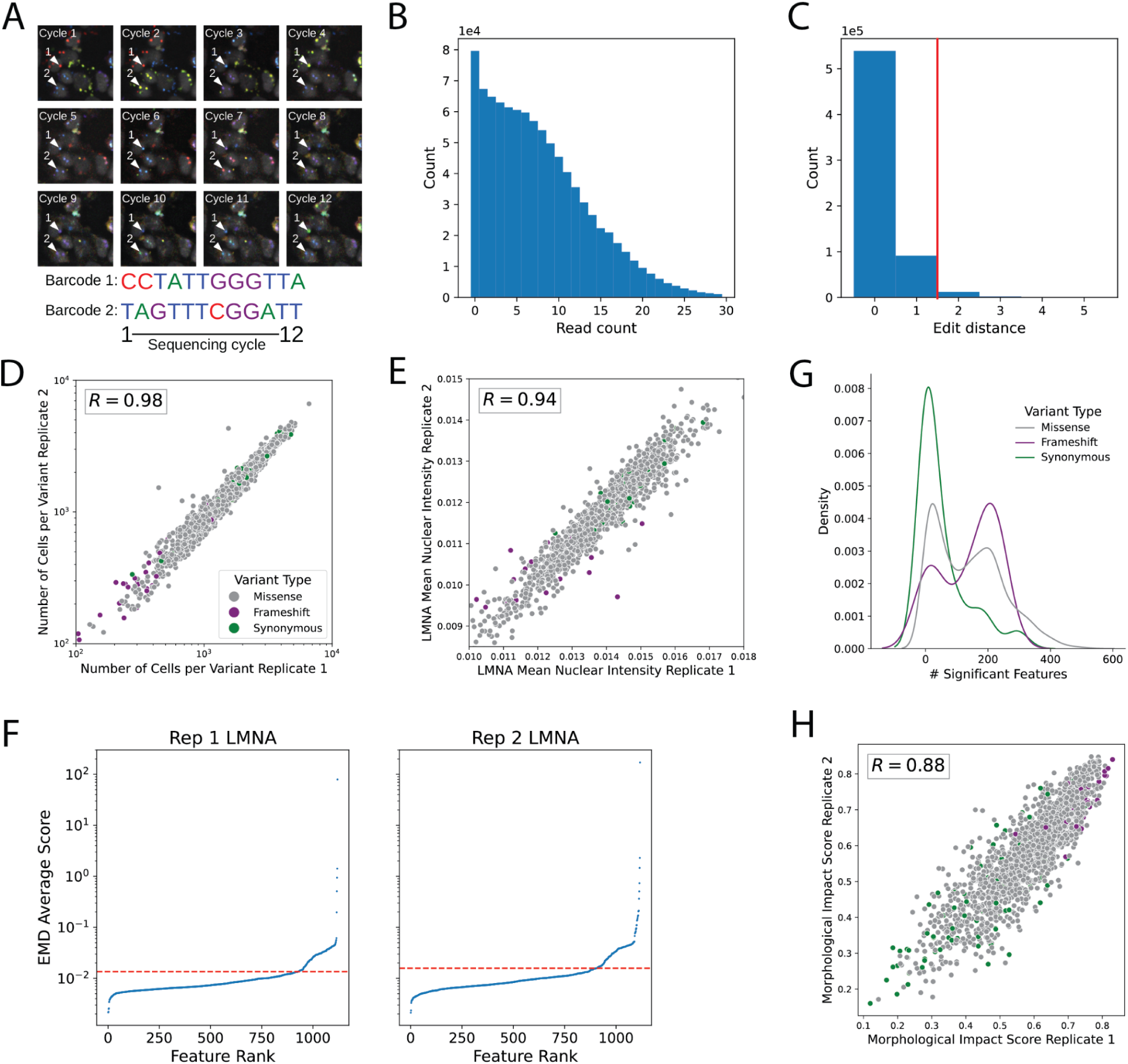
*LMNA* VIS-seq replication and visualization. (A) 12-base pair barcode sequences on circular RNAs were read by *in situ* sequencing by synthesis in each cell, and then mapped to corresponding *LMNA* variants using a barcode-to-variant dictionary made by long-read sequencing. Example cells with reads in all 12 cycles are shown. (B) Histogram of total number of 12-base pair reads per sequenced cell in single well of *LMNA* replicate 2 experiment. (C) Edit distance between consensus cell-level 12-base pair read and nearest library barcode in a single well of *LMNA* replicate 2 experiment. Red line indicates that cells with edit distance < 2 were used if they matched to a unique barcode. (D) Number of cells genotyped for each *LMNA* variant in both replicates of VIS-seq screen colored by variant type, with Pearson’s r shown. (E) Mean nuclear intensity in the mEGFP-lamin A channel (median over cells per variant) for variants in both replicates of VIS-seq screen colored by variant type as in (D), with Pearson’s r shown. (F) EMD reproducibility scores derived from 30 random partitions of wild-type *LMNA* expressing cells ranked by feature for replicate 1 (left) and replicate 2 (right). Low scores indicate high reproducibility. Threshold is drawn at 1.5 times the IQR added to the first quartile. Feature EMDs above this threshold in either replicate were removed in the feature selection step. (G) Number of significant features for each variant class determined by KS-test against wild-type cells, colored by variant type. Significance thresholded by Bonferroni-corrected p<0.001. (H) Morphological impact score of variants in both replicates of VIS-seq screen colored by variant type as in (D), with Pearson’s r shown.

**Supplementary Figure 3:**
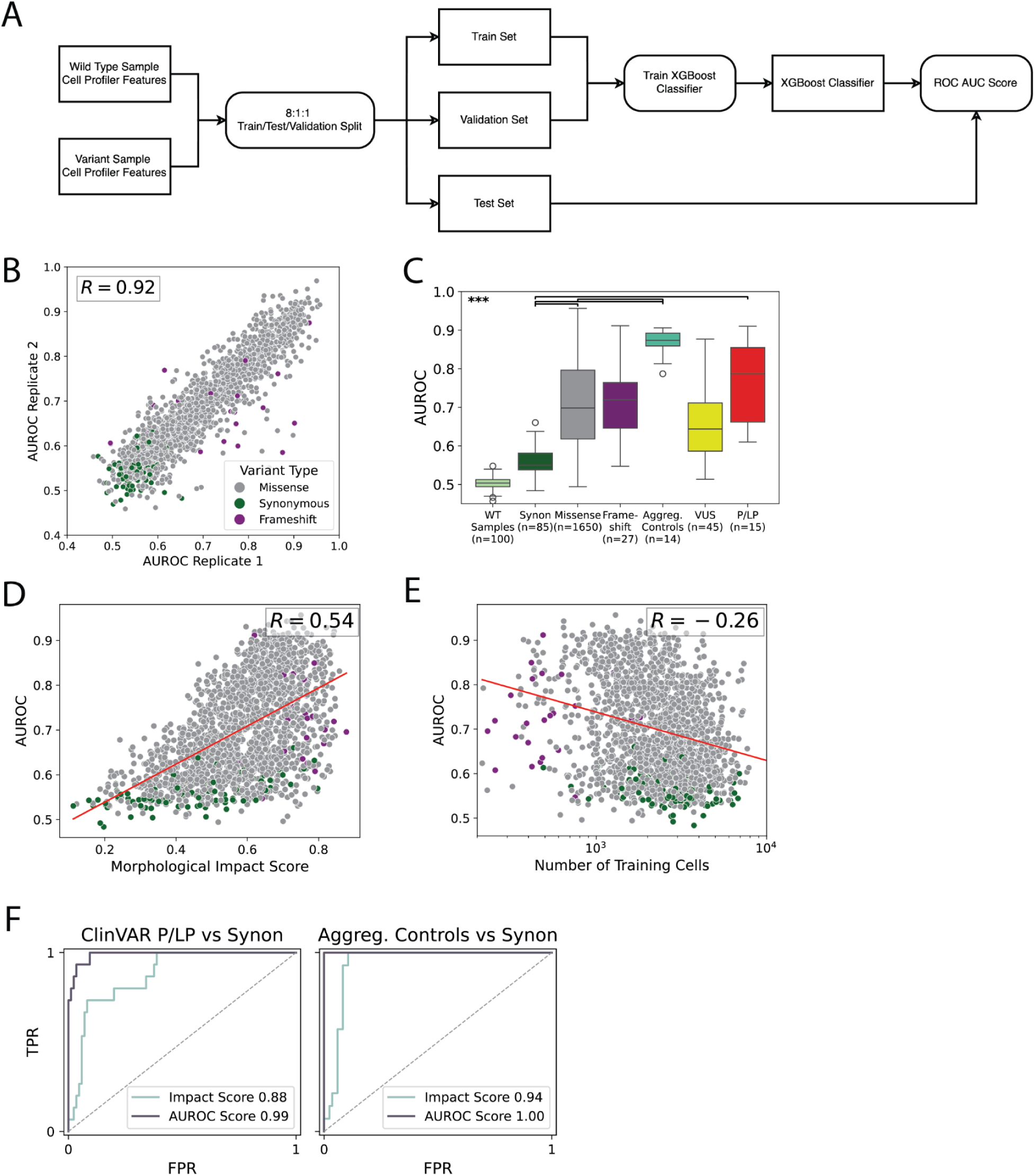
Training models to distinguish profiled *LMNA* variants from WT. (A) Flow-chart describing training binary classifiers for each variant to distinguish single cell images of that variant from corresponding WT images. For each variant as well as bootstraps of 1000 WT cells, an AUROC score summarizing classifier performance was computed on a test set of single cells. 0.5 indicates random classifier performance and 1 indicates perfect discrimination between variant and WT single cells. For a full description, see Methods. (B) AUROC of variants from both replicates of VIS-seq screen colored by variant type, with Pearson’s r shown. (C) AUROC scores for *LMNA* variants are plotted by variant type: missense, frameshift, or synonymous (Synon). 100 bootstrapped samples of WT variants are also shown for comparison. Scores are also shown for variants with >15% aggregated cells in HEK 293T previously measured by Anderson *et al*^39^. Lastly, AUC scores are plotted for ClinVar classifications of: variant of uncertain significance (VUS) and likely pathogenic/pathogenic (LP/P). *** indicates Mann-Whitney p-value < 0.001. (D) Scatterplot showing the AUROC score for each variant against the morphological impact score of that variant. Variants are colored by variant type. Best fit line shown in red. (E) Scatterplot showing the AUROC score for each variant against the number of training cells used for that variant. Variants are colored by variant type. Best fit line shown in red. (F) Receiver operating characteristic (ROC) curves are plotted for univariate zero-shot models trained on VIS-seq *LMNA* variant morphological impact or AUROC scores (solid lines, see Methods for details) predicting ClinVar pathogenicity or aggregation^39^ Area under the curve (AUC) scores are also shown for each model.

**Supplementary Figure 4:**
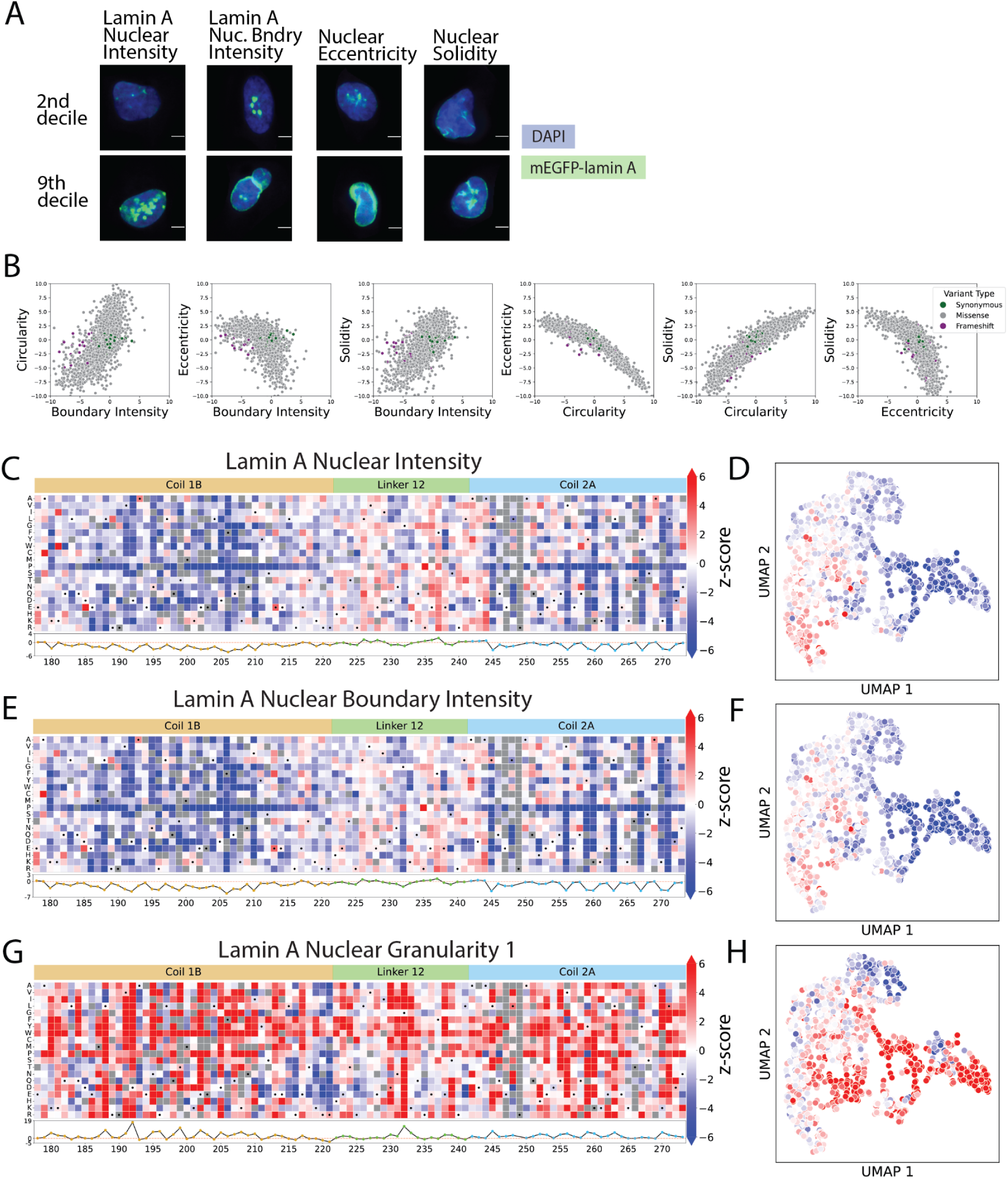
*LMNA* VIS-seq landmark feature heatmaps. (A) Randomly-selected cells from the second and ninth deciles in lamin A nuclear intensity, lamin A nuclear boundary intensity, nuclear eccentricity and nuclear solidity scores are shown. mEGFP-tagged lamin A channel is shown in green, and DAPI in blue. Scale bar indicates 5 μm. (B) *LMNA* feature z-scores for mEGFP-lamin A boundary intensity, nuclear eccentricity, nuclear solidity, and nuclear eccentricity are plotted against each other for all profiled variants. Z-scores are versus the synonymous variant distribution. Variant type is colored according to: synonymous variants (green), missense (grey), and frameshift variants (purple). (C) Heatmap for missense substitution effects on *LMNA* variant nuclear intensity z-scores. Grey boxes indicate missing variants and boxes with black dots indicate synonymous substitutions. Blue indicates low and red indicates high z-score for feature. *LMNA* subdomains are shown above the heatmap, and position-averaged z-score are plotted below the heatmap. (D) Lamin A nuclear intensity z-scores plotted on UMAP visualization of *LMNA* variant profiles, colored as in (C). (E) Heatmap for missense substitution effects on (median over cells) *LMNA* nuclear boundary intensity scores, colored and annotated as in (C). (F) Lamin A nuclear boundary intensity z-scores plotted on UMAP visualization of *LMNA* variant profiles, colored as in (C). (G) Heatmap for missense substitution effects on lamin A nuclear granularity 1 z-scores, colored and annotated as in (C). (H) Lamin A nuclear granularity 1 z-scores plotted on UMAP visualization of *LMNA* variant profiles, colored as in (C).

**Supplementary Figure 5:**
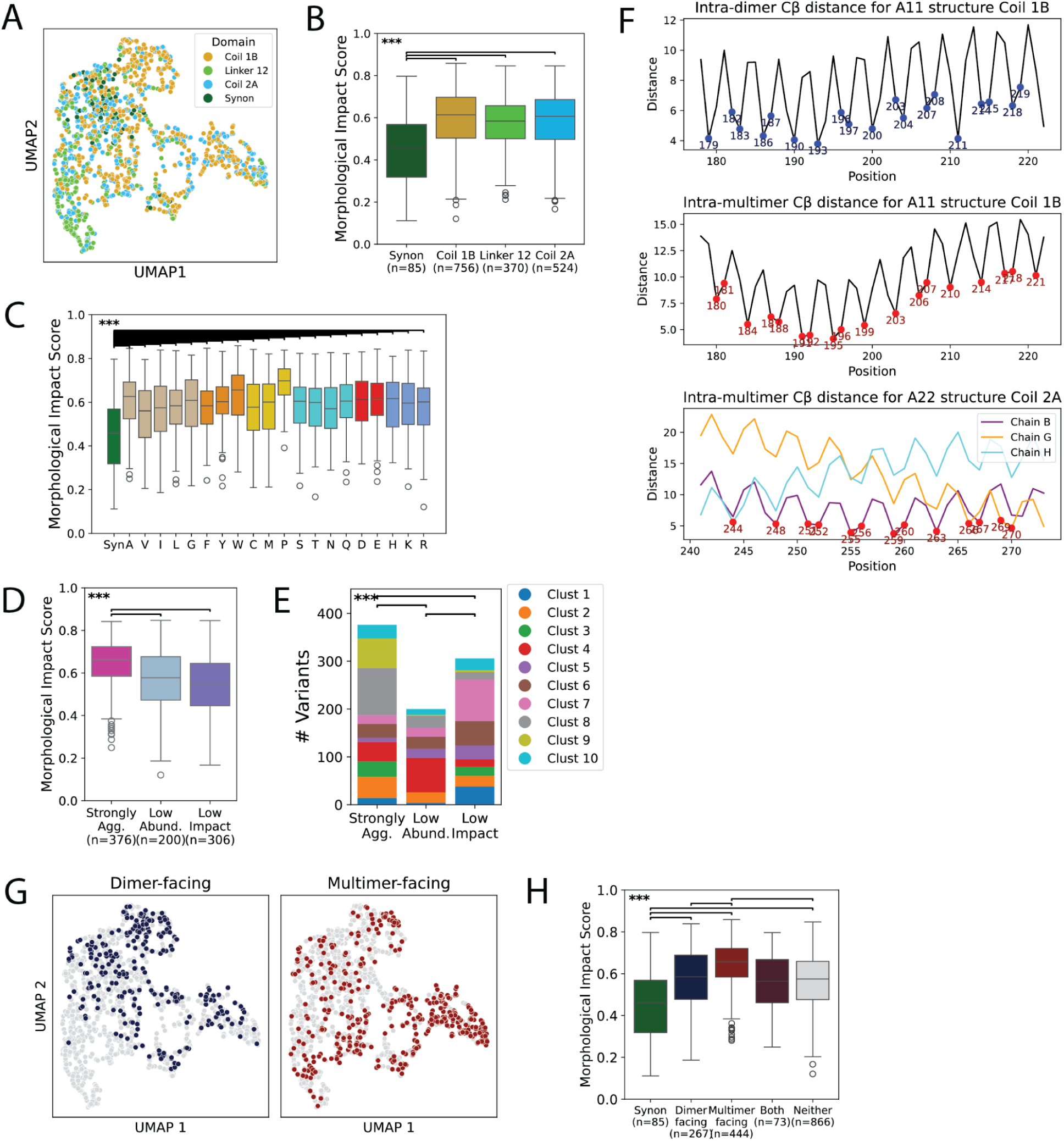
VIS-seq profiles separate lamin A residues by structural and functional properties. (A) UMAP visualization of *LMNA* synonymous (green) or missense variant profiles colored by lamin A domain. (B) Morphological impact score of missense substitutions plotted by domain, compared with synonymous variants (green). *** indicates Mann-Whitney U-test p<0.001. (C) Morphological impact score of lamin A missense substitutions plotted by amino acid, compared with synonymous variants (green). *** indicates Mann-Whitney U-test p<0.001 (D) Morphological impact score of missense substitutions at ⍺-helical position groups as defined in Fig. 3F by clustering amino acid positions. *** indicates Mann-Whitney U-test p<0.001. (E) Profiled ⍺-helical position variants by position cluster, colored by their Louvain cluster of residence as shown on UMAP representation in Fig. 2H. *** indicates *χ*^2^ p<0.001. (F) Intra-dimer (top) and inter-dimer coil 1B (middle) and coil 2A (bottom) minimum beta-carbon distances derived from A11^63^ and A22^64^ multimer structures. Dimer-facing residues are indicated in blue (top) and multimer-facing residues are indicated in red for A11^63^ and A22^64^ structures (middle, bottom, respectively). See Methods for how these residues are defined. (G) missense substitutions at dimer-facing and multimer-facing residues (as defined in (F)) plotted on the UMAP. (H) Morphological impact score of synonymous variants (green) and missense substitutions at dimer-facing positions (blue), multimer-facing positions (red), both (purple) and other non-interacting positions (gray). *** indicates Mann-Whitney U-test p<0.001.

**Supplementary Figure 6:**
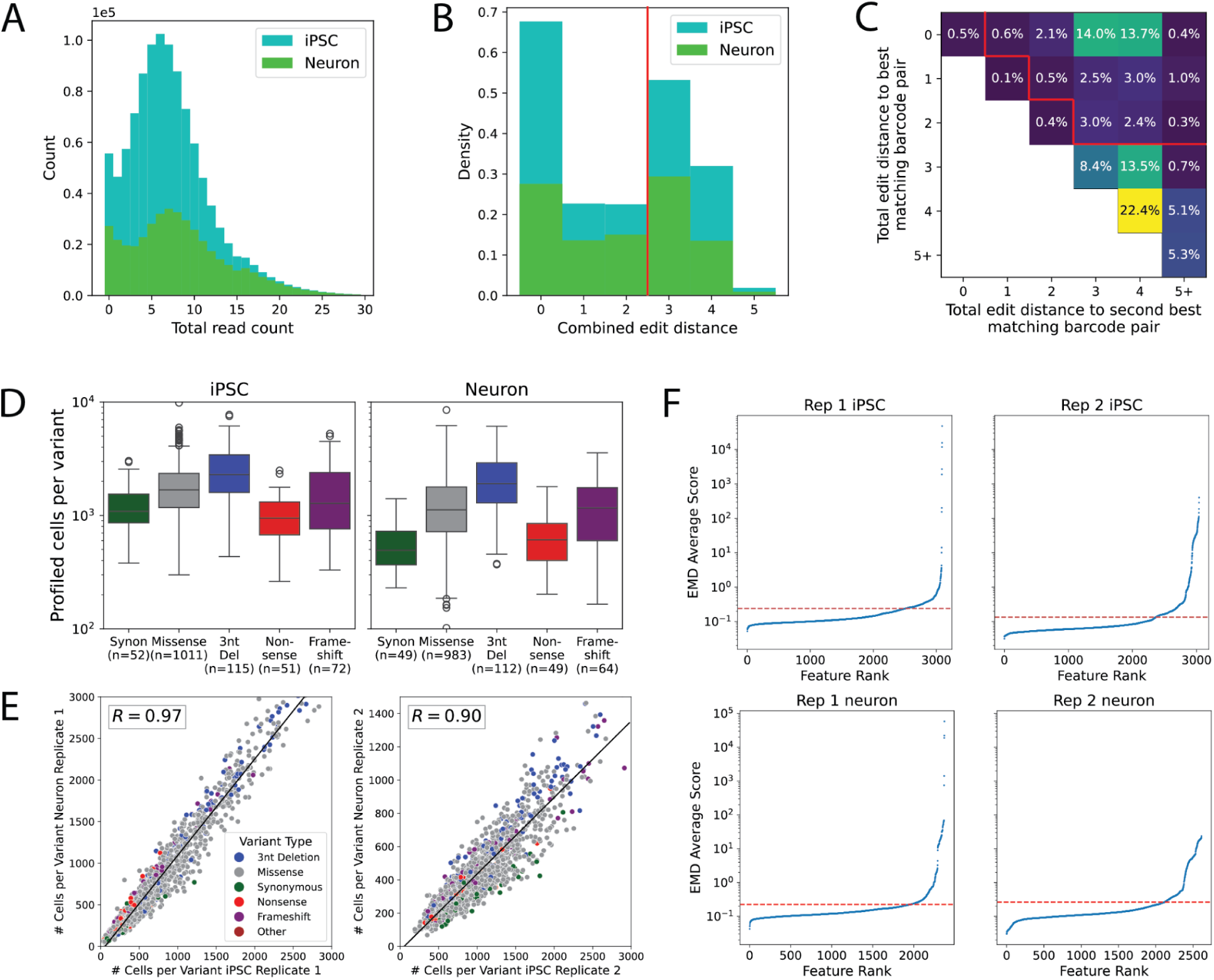
VIS-seq reproducibly called *PTEN* variants in single cells in iPS cells and neurons from two 8bp barcodes. (A) Histogram of total number of 8-base pair reads per genotyped cell in single well of *PTEN* iPS cell (blue) or neuron (green) replicate 2 experiments. (B) Total (summed) edit distance between consensus cell-level 16-base pair double barcode reads and nearest library double barcode in a single well of *PTEN* iPS cell (blue) or neuron (green) replicate 2 experiments. Red line indicates that cells with total edit distance < 3 were used if they matched to a unique library double barcode. (C) Total distance between consensus cell-level 16-base pair double barcode reads and nearest library double barcode (x) or second nearest library double barcode (y) in a single well of *PTEN* iPS cell replicate 2 experiment. Red line indicates that cells with total edit distance < 3 were used if they matched to a unique library double barcode. (D) Numbers of profiled single iPS cells (left) or neurons (right) containing single *PTEN* variants over the two replicate experiments are plotted by variant type. (E) Number of cells genotyped for each *PTEN* variant in iPSC (x) and neuron (y) of replicate 1 (left) and replicate 2 (right) of VIS-seq screens, with Pearson’s r shown. (F) EMD reproducibility scores derived from 30 random partitions of wild-type *PTEN* expressing iPS cells (top) or neurons (bottom) ranked by feature for replicate 1 (left) and replicate 2 (right). Low scores indicate high reproducibility. Threshold is drawn at 1.5 times the IQR added to the first quartile. Feature EMDs above this threshold in either replicate were removed in the feature selection step.

**Supplementary Figure 7:**
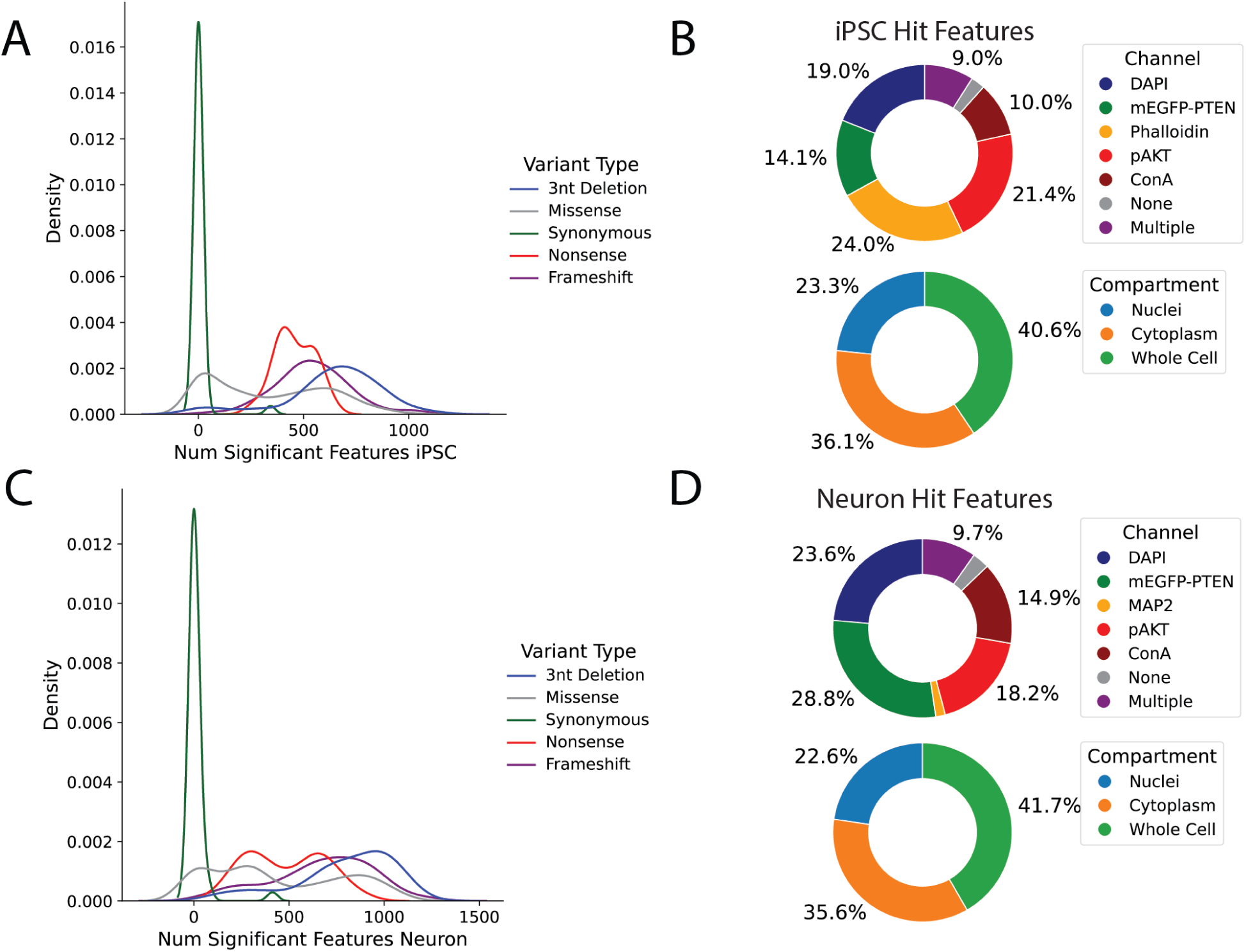
Significant features in VIS-seq *PTEN* experiments. (A) Number of significant features in iPSC for each variant class determined by KS-test against wild-type cells. Significance indicates Bonferroni-corrected p<0.001. (B) Hit iPS cell features, defined as being significantly different from WT (at Bonferroni-corrected KS-test p<0.001) in >=25 *PTEN* variants in iPS cells, are classified by imaging channel (top) or by compartment (bottom). (C) Number of significant features in neurons for each variant class determined by KS-test against wild-type cells. Significance indicates Bonferroni-corrected p<0.001. (D) Hit neuron features, defined as being significantly different from WT (at Bonferroni-corrected KS-test p<0.001) in >=25 *PTEN* variants in neurons, are classified by imaging channel (top) or by compartment (bottom).

**Supplementary Figure 8:**
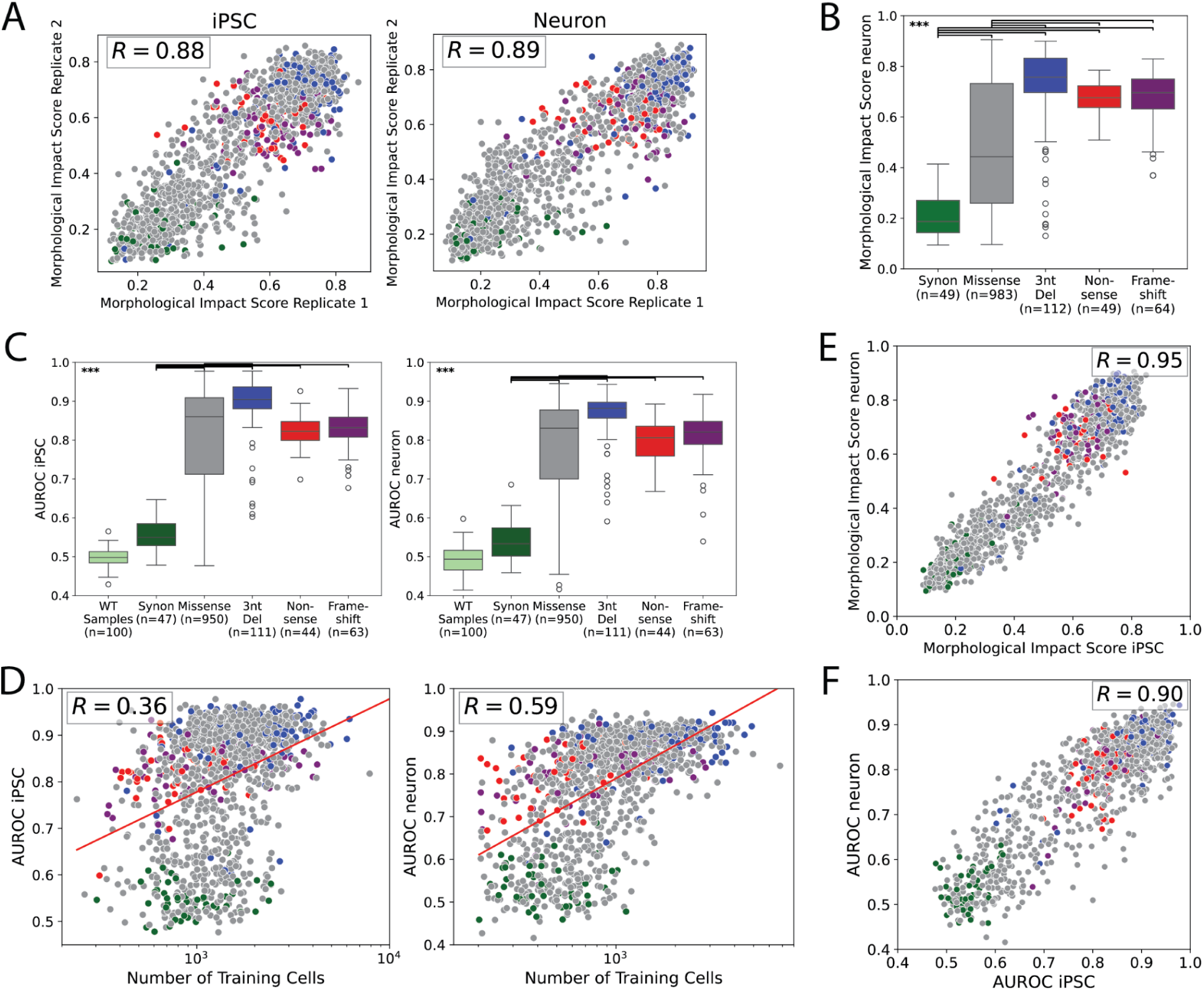
VIS-seq generated reproducible profiles in iPS cells and neurons. (A) Morphological impact scores for variants in each replicate of *PTEN* VIS-seq screen in iPSC (left) and neurons (right), colored by variant type as in (B), with Pearson’s r shown. (B) Morphological impact score for *PTEN* variant profiles in neurons, plotted by variant type. *** indicates Mann-Whitney p-value < 0.001. (C) AUROC scores for *PTEN* variants in iPS cells (left) or neurons (right), plotted by variant type. 100 bootstrapped samples of WT variants are also shown for comparison. *** indicates Mann-Whitney p-value < 0.001. (D) Scatterplot showing the AUROC score for each variant against the number of training iPS cells (left) or neurons (right) used for that variant. Variants are colored by variant type as in (B). Best fit line and Pearson’s r shown. (E) Morphological impact scores for *PTEN* variants in both iPSC (x) and neurons (y) are compared, colored by variant type as in (B). Pearson’s r shown. (F) AUROC scores for *PTEN* variants in both iPSC (x) and neurons (y) are compared, colored by variant type as in (B). Pearson’s r shown.

**Supplementary Figure 9:**
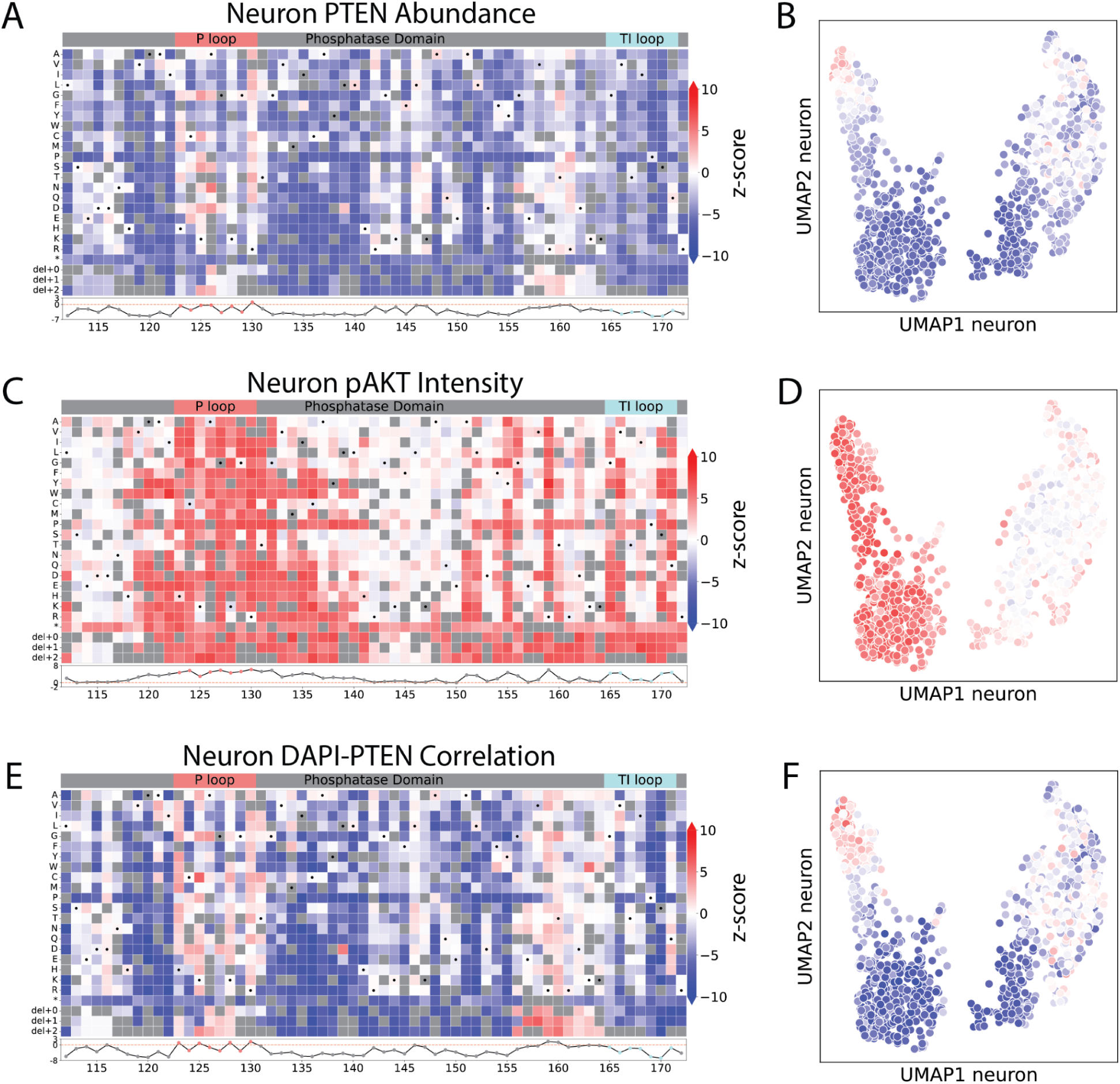
*PTEN* landmark features in neurons. (A) Positional heatmap (top) of PTEN intensity scores for missense, stop gain, or 3-nucleotide deletion variant profiles in neurons. Positional average of scores (bottom) with P-loop and TI-loop residues highlighted. (B) UMAP of *PTEN* variant morphological profiles in neurons colored by PTEN intensity score. Blue indicates low and red indicates high synonymous z-score for feature, colored as in (A). (C) Positional heatmap (top) of pAKT intensity scores for missense, stop gain, or 3-nucleotide deletion variant profiles in neurons. Positional average of scores (bottom) with P-loop and TI-loop residues highlighted. (D) UMAP of *PTEN* variant morphological profiles in neurons colored by pAKT intensity z-score (right). Blue indicates low and red indicates high synonymous z-score for feature, colored as in (C). (E) Positional heatmap (top) of DAPI-PTEN correlation scores for missense, stop gain, or 3-nucleotide deletion variant profiles in neurons. Positional average of scores (bottom) with P-loop and TI-loop residues highlighted. (F) UMAP of *PTEN* variant morphological profiles in neurons colored by DAPI-PTEN correlation. Blue indicates low and red indicates high synonymous z-score for feature, colored as in (E).

**Supplementary Figure 10:**
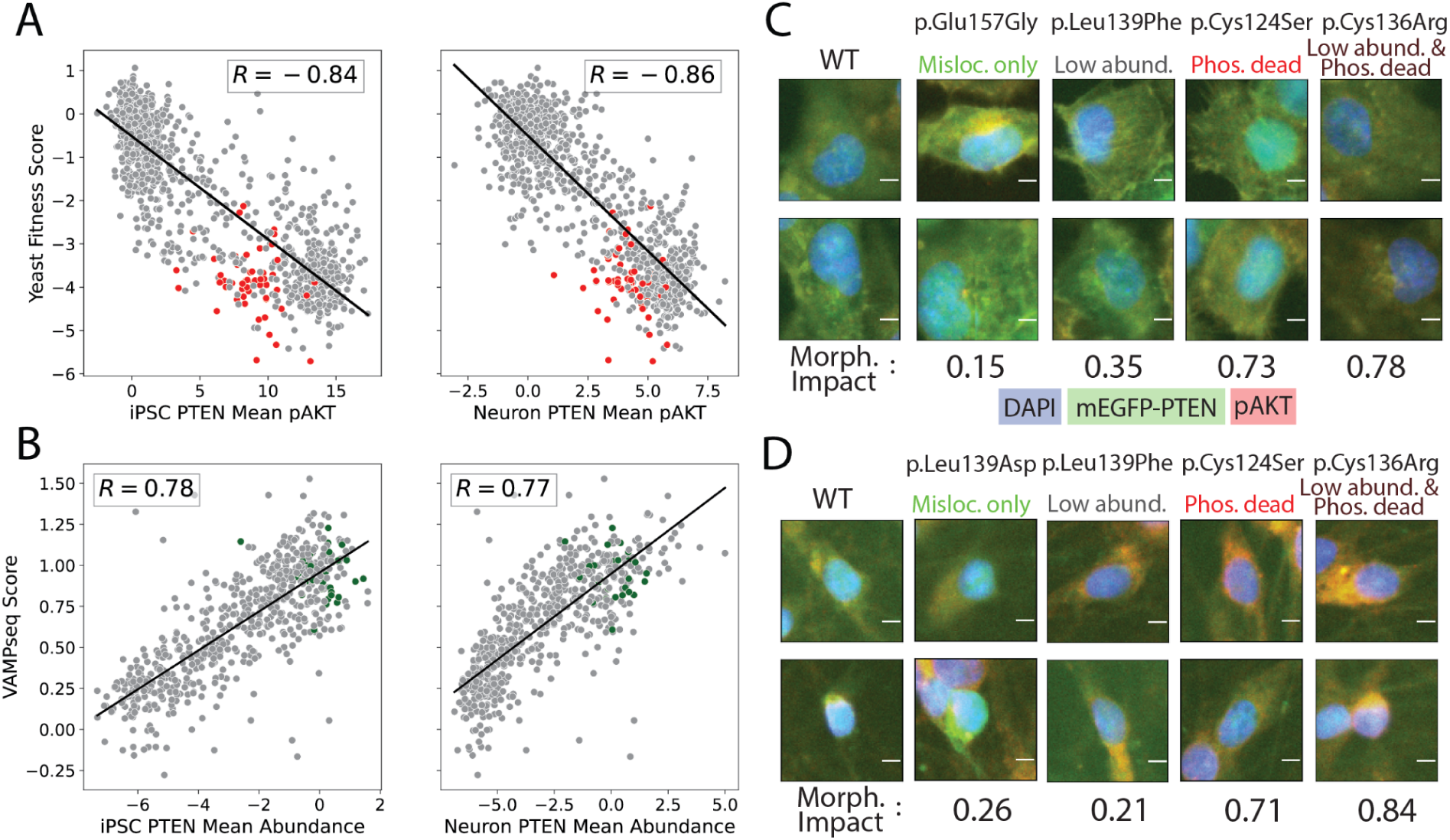
*PTEN* VIS-seq measurements correlate with prior variant effect measurements. (A) pAKT intensity score for *PTEN* variants in iPSC (left) and neurons (right) plotted against yeast fitness score, with best-fit line and Pearson’s r shown. (B) mEGFP-PTEN abundance score for *PTEN* variants in iPSC (left) and neurons (right) plotted against VAMP-seq score, with best-fit line and Pearson’s r shown. (C) Two randomly-selected iPS cells expressing *PTEN* variants are shown, with corresponding morphological impact scores. mEGFP-tagged PTEN channel is shown in green, DAPI in blue, and pAKT in red. Scale bar indicates 5 μm. (D) Two randomly-selected neurons expressing *PTEN* variants are shown, with corresponding morphological impact scores. mEGFP-tagged PTEN channel is shown in green, DAPI in blue, and pAKT in red. Scale bar indicates 5 μm.

**Supplementary Figure 11:**
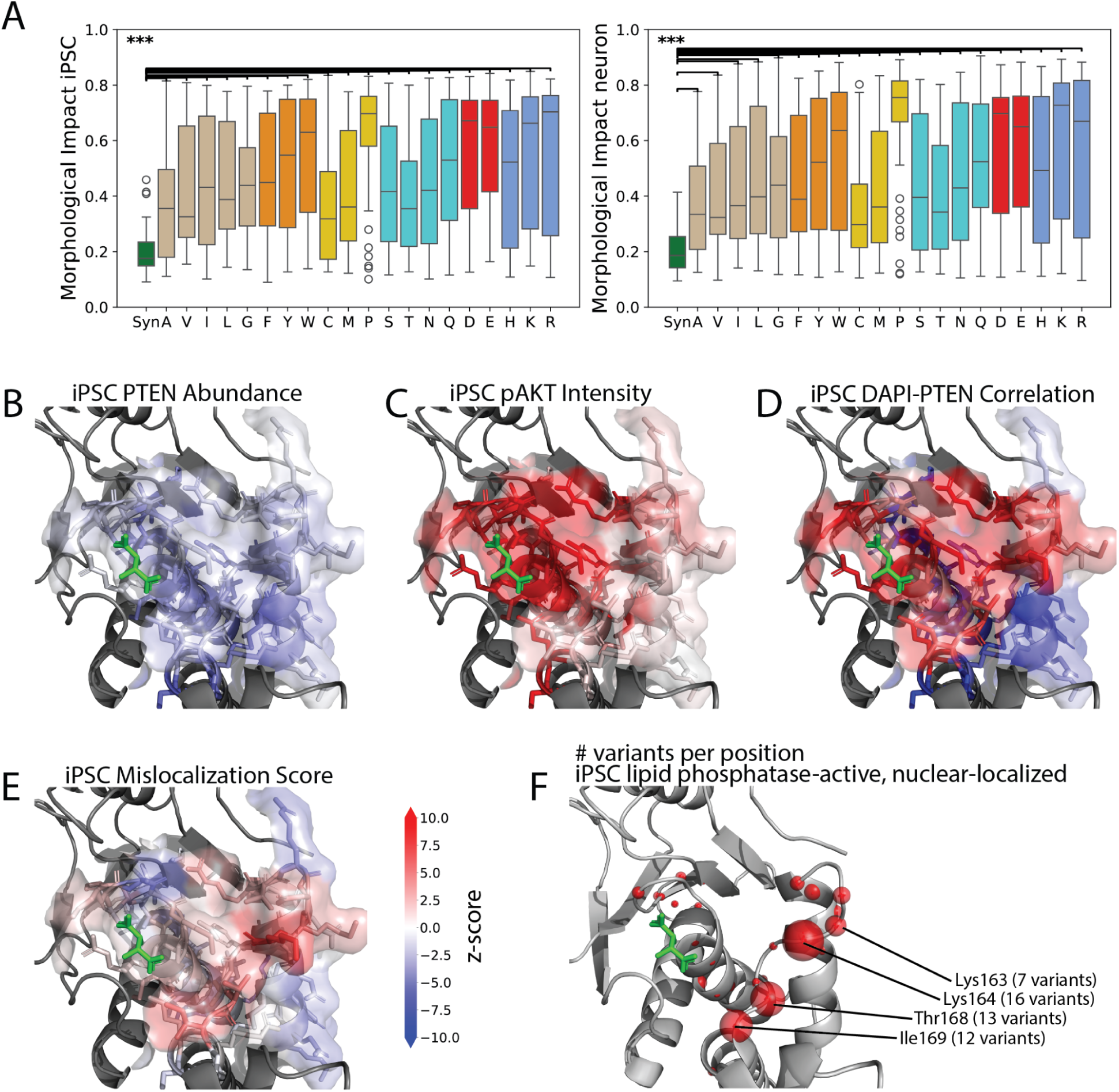
Structural analysis of *PTEN* VIS-seq profiles. (A) Morphological impact score of missense substitutions plotted by amino acid, compared with synonymous variants, in iPSC (left) and neurons (right). *** indicates Mann-Whitney U-test p<0.001 (B) iPSC mEGFP-PTEN intensity score positional averages are used to color the PTEN crystal structure (1D5R)^69^. Solvent-exposed surface is colored by the exposed residue’s z-score. Blue indicates low-intensity positions. The bound tartrate molecule is shown in green. Residue color scale shown to the right of (E). (C) iPSC pAKT intensity score positional averages are used to color the PTEN crystal structure (1D5R)^69^. Red indicates positions with high pAKT intensity and corresponding low lipid phosphatase activity. Residue color scale shown to the right of (E). (D) iPSC DAPI-PTEN correlation score positional averages are used to color the PTEN crystal structure (1D5R)^69^. Blue indicates cytoplasmic-localizing positions and red indicates nuclear-localizing positions. Residue color scale shown to the right of (E). (E) iPSC mislocalization score (see Methods) positional averages are used to color the PTEN crystal structure (1D5R)^69^. Blue indicates aberrantly cytoplasmic-localizing positions and red indicates aberrantly nuclear-localizing positions, after the effects of activity and abundance are removed. Residue color scale shown to the right. (F) PTEN structure (1D5R)^69^ with spheres centered at ⍺-carbon atoms with radii indicating the number of lipid-phosphatase active (defined as pAKT z-score<2.5) and nuclear-localized (defined as DAPI-PTEN correlation z-score>2.5) variants at that position. The top four positions indicating nuclear localization are highlighted, with the number of variants at each of these positions indicated.

**Supplementary Figure 12:**
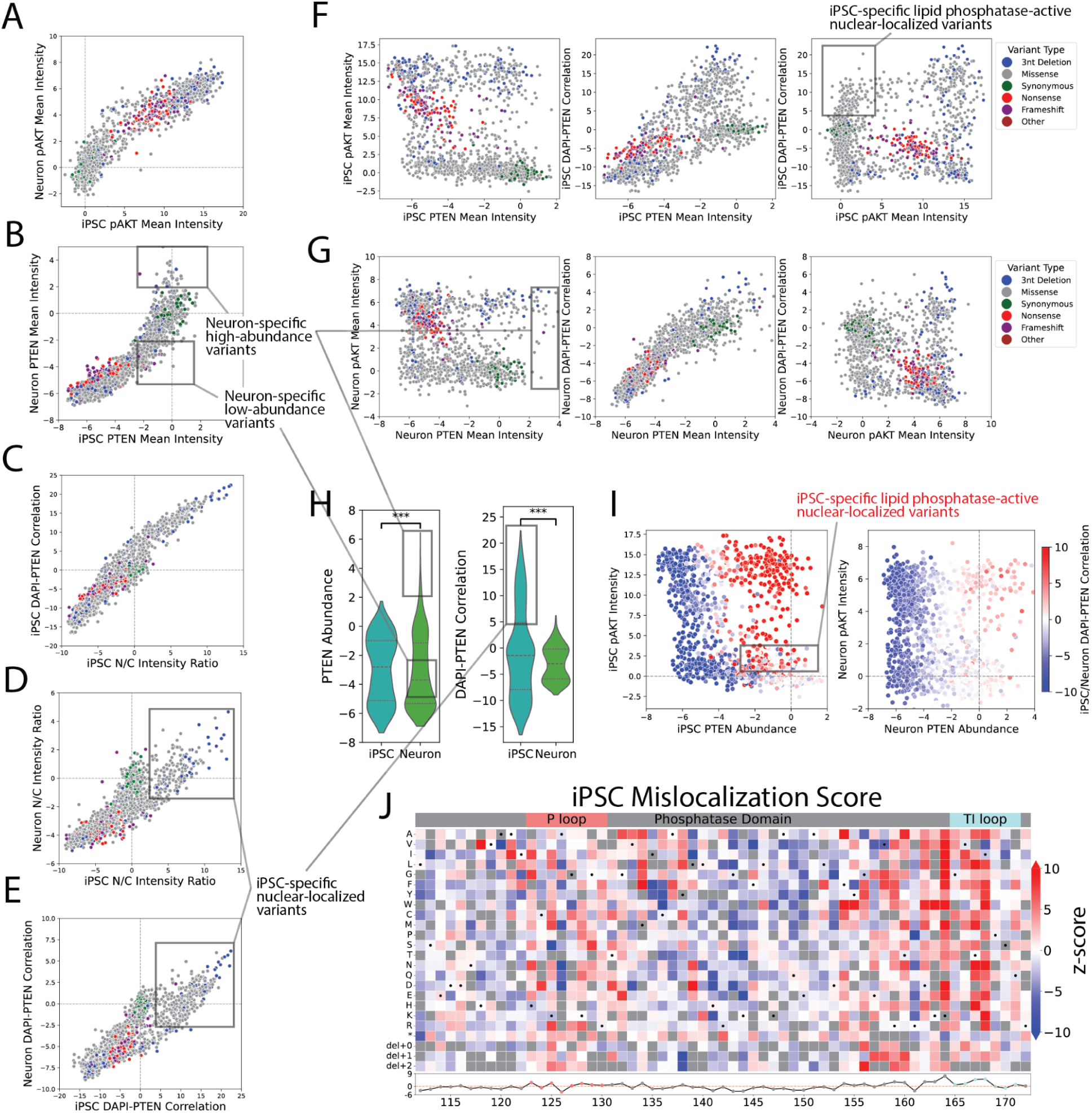
Relationships between *PTEN* landmark features in iPS cells and neurons. (A) iPSC pAKT intensity z-scores are plotted against neuron pAKT intensity z-scores for all profiled variant, colored by variant type as in (F). Z-scores are versus the synonymous variant distribution. (B) iPSC mEGFP-PTEN intensity z-scores are plotted against neuron mEGFP-PTEN intensity z-scores for all profiled variants, colored by variant type as in (F). Both neuron-specific low- and high-abundance variants are indicated. (C) iPSC nucleus to cytoplasm PTEN intensity ratio z-scores are plotted against iPSC DAPI-PTEN correlation z-scores for all profiled variants, colored by variant type as in (F). (D) iPSC nucleus to cytoplasm PTEN intensity ratio z-scores are plotted against neuron nucleus to cytoplasm PTEN intensity ratio z-scores, colored by variant type as in (F). iPSC-specific nuclear-localized variants are indicated. (E) iPSC DAPI-PTEN correlation z-scores are plotted against neuron DAPI-PTEN correlation z-scores for all profiled variants, colored by variant type as in (F). iPSC-specific nuclear-localized variants are indicated. (F) iPSC PTEN landmark feature z-scores (PTEN intensity, pAKT intensity, and DAPI-PTEN correlation) are plotted against each other for all profiled variants. Variant type is colored according to: synonymous variants (green), missense (grey), 3-nt deletions (blue), nonsense (red), and frameshift variants (purple). iPSC-specific lipid phosphatase-active nuclear-localized variants are indicated. (G) Neuron PTEN landmark feature z-scores are plotted against each other for all profiled variant, colored by variant type as in (F). Neuron-specific high-abundance variants are indicated. (H) Violin plots of iPS cell and neuron PTEN abundance (left) and DAPI-PTEN correlation z-scores (right). Neuron-specific low- and high-abundance variants and iPSC-specific nuclear-localized variants are indicated. *** indicates KS p-value<0.001. (I) iPSC (left) or neuron (right) PTEN abundance z-scores are plotted against pAKT intensity z-scores, colored by either iPSC DAPI-PTEN correlation (left) or neuron DAPI-PTEN correlation (right) z-scores. iPSC-specific phosphatase-active, nuclearly-localized variants are indicated on the iPSC plot, as red-colored variants (DAPI-PTEN correlation z-score > 2.5) with pAKT intensity z-score < 2.5. (J) The mislocalization score is defined as the variant-level residuals when DAPI-PTEN correlation is regressed against PTEN intensity and pAKT intensity (see Methods). Positional heatmap (top) of mislocalization scores for missense, stop gain, or 3-nucleotide deletion variant profiles in iPSC. Positional average of scores (bottom) with P-loop and TI-loop residues highlighted. Blue indicates low score and red indicates high score.

**Supplementary Figure 13:**
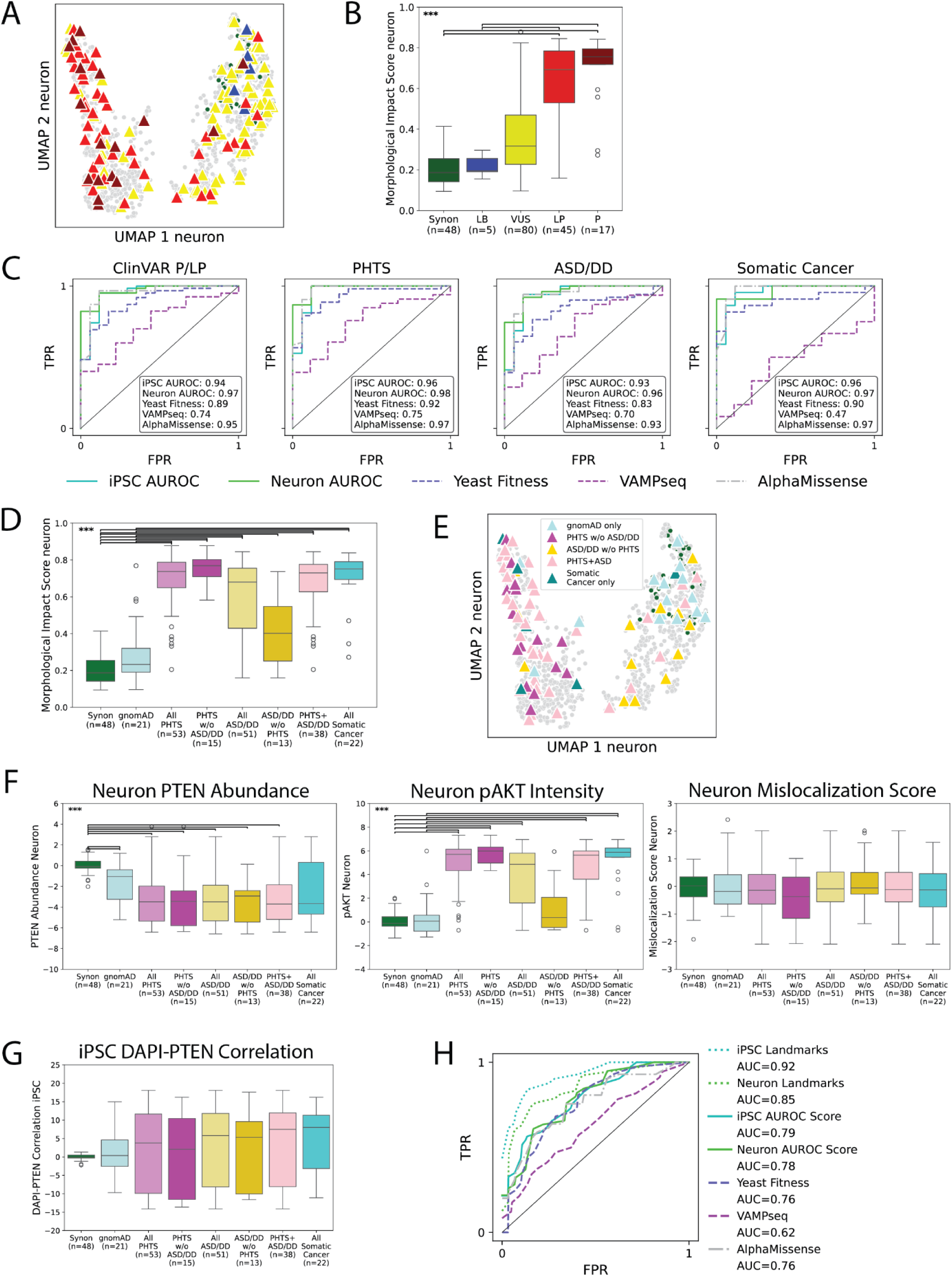
*PTEN* VIS-seq profiles predict pathogenicity and clinical phenotype. (A) UMAP visualization of neuron *PTEN* variant profiles. Triangles indicate variants in ClinVar classified as likely benign (LB, blue), likely pathogenic (LP, red), pathogenic (P, dark red), or variant of uncertain significance (VUS, yellow). All profiled variants are plotted in the background colored green (synonymous) or grey (otherwise) for comparison. (B) Morphological impact scores for *PTEN* neuron profiles are plotted by ClinVar label. Synonymous variants (green) are included for comparison. *** indicates Mann-Whiney U p<0.001. (C) Receiver operating characteristic (ROC) curves are plotted for univariate zero-shot models predicting ClinVar pathogenicity or each clinical phenotype (ASD = autism spectrum disorder, DD = developmental delay, PHTS = *PTEN* hamartoma tumor syndrome; see Methods for curation criteria) from iPSC and neuron AUROC score (this publication, solid lines), yeast fitness scores^83^ (dashed line), VAMPseq scores^5^ (dashed line), or AlphaMissense scores^91^ (dot-dashed line). Area under the curve (AUC) scores are shown in the box for each model. (D) Morphological impact scores for *PTEN* neuron profiles are plotted by variant association with clinical phenotypes. gnomAD v4.1 (light blue) variants and synonymous variants (green) are plotted for comparison. *** indicates Mann-Whiney U p<0.001. (E) UMAP visualization of iPS cell *PTEN* variant profiles. Triangles indicate association with clinical phenotypes. gnomAD v4.1 (light blue) variants are also plotted. All profiled variants are plotted in the background colored green (synonymous) or grey (otherwise) for comparison. (F) Feature scores for *PTEN* variants in neurons are plotted by variant association with clinical phenotypes. gnomAD v4.1 (light blue) variants and synonymous variants (green) are plotted for comparison. Features plotted include PTEN intensity (left), pAKT intensity (center) and mislocalization score (right, see Methods).*** indicates Mann-Whitney U-test p<0.001 (G) Feature scores for *PTEN* variants in iPSC are plotted by variant association with landmark feature iPS cell DAPI-PTEN correlation z-scores relative to the synonymous variant distribution. gnomAD v4.1 (light blue) variants and synonymous variants (green) are plotted for comparison. (H) Receiver operating characteristic (ROC) curves produced by macro-averaging sensitivity and specificity over classes for models trained on iPS and neuron VIS-seq landmark features (dotted lines) as well as scores from (C) classifying gnomAD controls from PHTS-associated variants from ASD/DD-associated variants. AUC is shown on the right for each model.

**Supplementary Figure 14:**
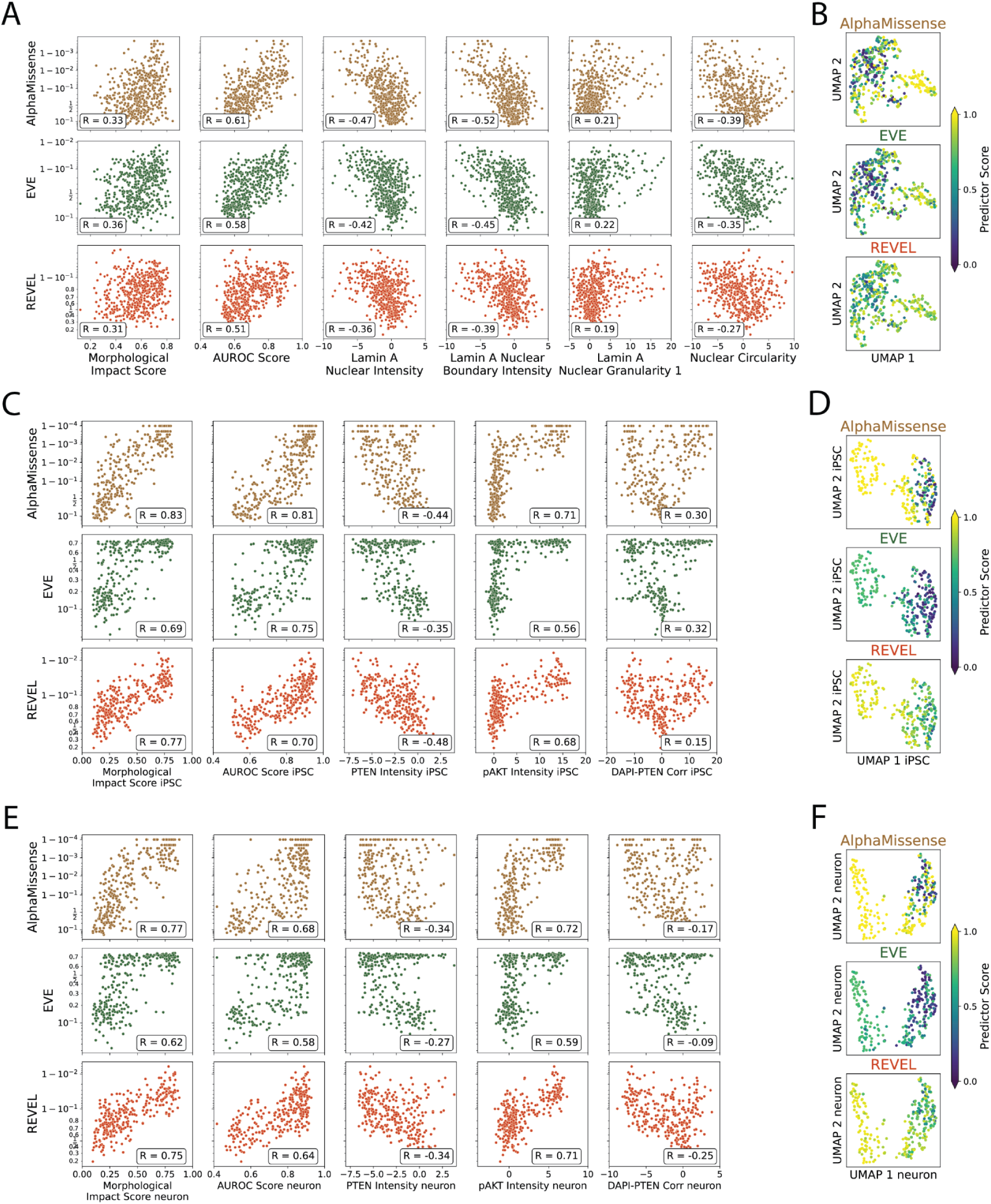
VIS-seq profiles elaborate computational variant effect predictions. (A) Predictor scores for AlphaMissense^91^, EVE^92^, and REVEL^93^ are plotted against *LMNA* VIS-seq morphological impact score and landmark features. Pearson’s r between logit-transformed predictor scores and VIS-seq scores are shown on the bottom right. (B) Predictor scores for AlphaMissense^91^, EVE^92^, and REVEL^93^ are colored on the UMAP visualization of *LMNA* VIS-seq profiles. (C) Predictor scores for AlphaMissense^91^, EVE^92^, and REVEL^93^ are plotted against *PTEN* VIS-seq iPSC morphological impact score and landmark features. Pearson’s r between logit-transformed predictor scores and VIS-seq scores are shown on the bottom left. (D) Predictor scores for AlphaMissense^91^, EVE^92^, and REVEL^93^ are colored on the UMAP visualization of *PTEN* iPSC VIS-seq profiles. (E) Predictor scores for AlphaMissense^91^, EVE^92^, and REVEL^93^ are plotted against *PTEN* VIS-seq neuron morphological impact score and landmark features. Pearson’s r between logit-transformed predictor scores and VIS-seq scores are shown on the bottom left. (F) Predictor scores for AlphaMissense^91^, EVE^92^, and REVEL^93^ are colored on the UMAP visualization of *PTEN* neuron VIS-seq profiles.

## Notes

https://mavedb.org/experiments/urn:mavedb:00001243-a

https://mavedb.org/experiments/urn:mavedb:00001244-b

https://mavedb.org/experiments/urn:mavedb:00001244-a

https://zenodo.org/records/15787685

https://visseq.gs.washington.edu/

